# When Darkness Becomes a Ray of Light in the Dark Times: Understanding the COVID-19 via the Comparative Analysis of the Dark Proteomes of SARS-CoV-2, Human SARS and Bat SARS-Like Coronaviruses

**DOI:** 10.1101/2020.03.13.990598

**Authors:** Rajanish Giri, Taniya Bhardwaj, Meenakshi Shegane, Bhuvaneshwari R. Gehi, Prateek Kumar, Kundlik Gadhave, Christopher J. Oldfield, Vladimir N. Uversky

**Affiliations:** Indian Institute of Technology Mandi, School of Basic Sciences, VPO Kamand, Himachal Pradesh 175005, India; Department of Computer Science, Virginia Commonwealth University, Richmond, VA, USA; Department of Molecular Medicine and Byrd Alzheimer’s Research Institute, Morsani College of Medicine, University of South Florida, Tampa, Florida, United States of America; Laboratory of New Methods in Biology, Institute for Biological Instrumentation of the Russian Academy of Sciences, Federal Research Center “Pushchino Scientific Center for Biological Research of the Russian Academy of Sciences”, Pushchino, Moscow region, 142290 Russia

**Keywords:** COVID-19, SARS-CoV-2, Coronavirus, Bat CoV, Human SARS, Intrinsically Disordered Proteins, Molecular Recognition Features

## Abstract

Recently emerged coronavirus designated as SARS-CoV-2 (also known as 2019 novel coronavirus (2019-nCoV) or Wuhan coronavirus) is a causative agent of coronavirus disease 2019 (COVID-19), which is rapidly spreading throughout the world now. More than 9,00,000 cases of SARS-CoV-2 infection and more than 47,000 COVID-19-associated mortalities have been reported worldwide till the writing of this article, and these numbers are increasing every passing hour. World Health Organization (WHO) has declared the SARS-CoV-2 spread as a global public health emergency and admitted that the COVID-19 is a pandemic now. The multiple sequence alignment data correlated with the already published reports on the SARS-CoV-2 evolution and indicated that this virus is closely related to the bat Severe Acute Respiratory Syndrome-like coronavirus (bat SARS-like CoV) and the well-studied Human SARS coronavirus (SARS CoV). The disordered regions in viral proteins are associated with the viral infectivity and pathogenicity. Therefore, in this study, we have exploited a set of complementary computational approaches to examine the dark proteomes of SARS-CoV-2, bat SARS-like, and human SARS CoVs by analysing the prevalence of intrinsic disorder in their proteins. According to our findings, SARS-CoV-2 proteome contains very significant levels of structural order. In fact, except for Nucleocapsid, Nsp8, and ORF6, the vast majority of SARS-CoV-2 proteins are mostly ordered proteins containing less intrinsically disordered protein regions (IDPRs). However, IDPRs found in SARS-CoV-2 proteins are functionally important. For example, cleavage sites in its replicase 1ab polyprotein are found to be highly disordered, and almost all SARS-CoV-2 proteins were shown to contain molecular recognition features (MoRFs), which are intrinsic disorder-based protein-protein interaction sites that are commonly utilized by proteins for interaction with specific partners. The results of our extensive investigation of the dark side of the SARS-CoV-2 proteome will have important implications for the structural and non-structural biology of SARS or SARS-like coronaviruses.

**Significance:** The infection caused by a novel coronavirus (SARS-CoV-2) that causes severe respiratory disease with pneumonia-like symptoms in humans is responsible for the current COVID-19 pandemic. No in-depth information on structures and functions of SARS-CoV-2 proteins is currently available in the public domain, and no effective anti-viral drugs and/or vaccines are designed for the treatment of this infection. Our study provides the first comparative analysis of the order- and disorder-based features of the SARS-CoV-2 proteome relative to human SARS and bat CoV that may be useful for structure-based drug discovery.

## Introduction

Emerging Coronavirus disease 2019 (COVID-19) is a recent pandemic, which is recently declared as a public health emergency by the World Health Organization (WHO). Since its first appearance in visitors of the Wuhan’s seafood and meat market, China, reported in December 2019, COVID-19 has now large scale socio-economic impact [1]. According to the WHO, till 2^nd^ April 2020, the infection spread over at least 170 countries and territories, where there were more than 0.93 million confirmed cases, with more than 47,000 patients died due to COVID-19. One should also keep in mind that these data on the COVID-19 spread and related casualties are rapidly becoming outdated, almost with the speed of typing of these sentences. Symptoms of COVID-19, which is a respiratory illness, include fever, cough, sore throat, shortness of breath, as well as mild gastrointestinal (GI) symptoms in some patients, and, in more serious cases, the infection can cause severe pneumonia and even death [2].

According to the International Committee on Taxonomy of Viruses (ICTV), SARS-CoV-2 comes under the coronavirinae sub-family of *coronaviridae* family of order nidovirales. Viruses of the nidovirales order are enveloped, non-segmented positive-sense, single-stranded RNA viruses [3]. The major variations among *Nidovirus* family occur in the number, type, and sizes of the structural proteins [3]. The family *coronaviridae* comprises of vertebrate infecting viruses that transmit horizontally mainly through oral/fecal route and cause gastrointestinal and respiratory problems to the host [4]. Sub-family *coronavirinae* consists of four genera, namely: alpha, beta, gamma, and delta coronaviruses based on the phylogenetic clustering of viruses [5,6]. *Coronavirinae* having the largest genomes among the RNA viruses, incorporate their ~30kb genomes inside the enveloped capsid. These viruses have variations among genome due to significant differences in the structure and morphology of their nucleocapsids [7].

SARS-Coronavirus genomic RNA includes a 5’ cap, leader sequence, UTR, a replicase gene, genes for structural and accessory proteins, 3’ UTR, and a poly-A tail (**Figure 1**). Two-third of genome codes for replicase gene (~20kb) containing genes for all non-structural viral proteins, while the remaining part of the genome (~10kb) contains genes for accessory proteins interspersed between the genes responsible for coding structural proteins [7]. For the expression of structural and accessory proteins, additional transcriptional regulatory sequences (TRS) are present within the viral genome. 5’ and 3’ UTRs contain stem-loop regions required for viral RNA synthesis [8]. The ~20kb (replicase gene) ssRNA is translated first into two long polyproteins: replicase polyprotein 1a and 1ab inside host cells. The newly formed polyproteins, after cleavage by two viral proteases, result in 16 non-structural proteins that perform a wide range of functions for viruses inside the host cell. They also induce ER-derived double-membrane vesicles (DMVs) for viral replication and transcription. Structural proteins shape the outer cover of the virion, while accessory proteins are mostly involved in host immune evasion [9,10].

**Figure 1.**
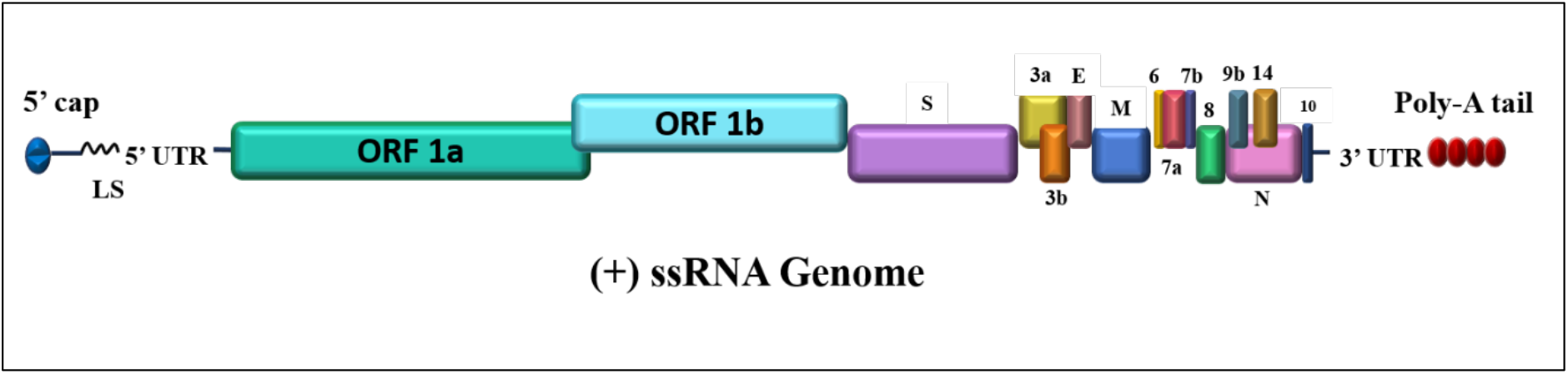
Genome architecture of SARS-CoV-2. SARS-CoV-2 contains a positive single-stranded mRNA as genetic material. The 5’ capped mRNA has a leader sequence (LS), poly-A tail at 3’ end, and 5’ and 3’ UTR. It contains the following genes: ORF1a, ORF1b, Spike (S), ORF3a, ORF3b, Envelope (E), Membrane (M), ORF6, ORF7a, ORF7b, ORF8, ORF9b, ORF14, Nucleocapsid (N), and ORF10.

In this study, we analyzed the dark side of the SARS-CoV-2 proteome (i.e., a part of a proteome that includes proteins or protein regions, which are not amenable to experimental structure determination by existing means and inaccessible to homology modelling; i.e., intrinsically disordered proteins (IDPs) and intrinsically disordered protein regions (IDPRs)), in order to better understand an interplay between the ordered and disordered components of the proteome. In classical structure-function-paradigm, it is believed that a unique, stable, and well-defined 3-dimensional structure is a prerequisite for a protein to accomplish its unique biological function. Although this notion dominated scientific minds for over the hundred years, eventually an idea of the presence of functional intrinsic disorder in proteins came to the attention of the structural biologists. According to this “heretic” viewpoint, a noticeable amount of biologically active proteins (of protein regions) fail to fold into the well-defined structures and instead remain disordered, existing as highly dynamic ensembles of rapidly interconverting conformations under the physiological conditions. These proteins and protein regions are known now as intrinsically disordered proteins (IDPs) and intrinsically disordered protein regions (IDPRs), respectively. The propensity of being functional intrinsically disordered proteins (similar to the propensity of forming unique biologically active structures of ordered proteins) is determined by the amino acid sequences [11–13]. IDPs exhibit their biological functions in numerous biological processes commonly associated with cellular signalling, gene regulation, and control by interacting with their physiological partners [14–18]. These functions of IDPs and IDPs are regulated by their protein-protein, protein-RNA, protein-DNA interactions [19,20]. Molecular recognition features (MoRFs) are the regions in IDPs implicated in regulation of IDPs function by protein-protein interactions and serve as the primary stage in molecular recognition.

Zhang and colleagues have reported the genomic sequence of SARS-CoV-2 with GenBank accession number NC_045512 having 29,903 nucleotides. The virus was isolated from the bronchoalveolar lavage fluid of a patient, went through a circle or renaming, from novel Wuhan seafood market pneumonia virus to deadly Wuhan coronavirus, to the 2019 novel coronavirus (2019-nCoV) or the Wuhan-2019 novel coronavirus (Wuhan-2019-nCoV, and was eventually named SARS-CoV-2 by the WHO [21].

It is known that the IDPs/IDPRs are present in all three kingdoms of life, and viral proteins often contain unstructured regions that have been strongly correlated with their virulence [22–25]. In this report, we investigated the disordered side of the SARS-CoV-2 proteome using a complementary set of computational approaches to check the prevalence of IDPRs in various SARS-CoV-2 proteins and to shed some light on their disorder-related functions. We also have comprehensively analyzed IDPRs among the closely related viruses, such as human SARS CoV and bat SARS-like CoV. Furthermore, we have also identified protein functions related to protein-protein interactions, RNA binding, and DNA binding from all three viruses. Since these three viruses are closely related, our study provides important means for a better understanding of the sequence and structural peculiarities of their evolution. We believe that this study will help the structural and non-structural biologists to design and perform experiments for a more in-depth understanding of this virus and its pathogenicity. This also will have long-term implications for developing new drugs or vaccines against this currently unpreventable infection.

## Materials and Methods

### Sequence retrieval and multiple sequence alignment

The protein sequences of Bat CoV (SARS-like) and Human SARS CoV were retrieved from UniProt (UniProt IDs for individual proteins are listed in **Table 1**). The translated sequences of SARS-CoV-2 proteins [GenBank database [26] (Accession ID: NC_045512.2)] were obtained from GenBank. We used these sequences for performing multiple sequence alignment (MSA) and predicting the IDPRs. We have used Clustal Omega [27] for protein sequence alignment and Esprit 3.0 [28] for constructing the aligned images.

**Table 1:**
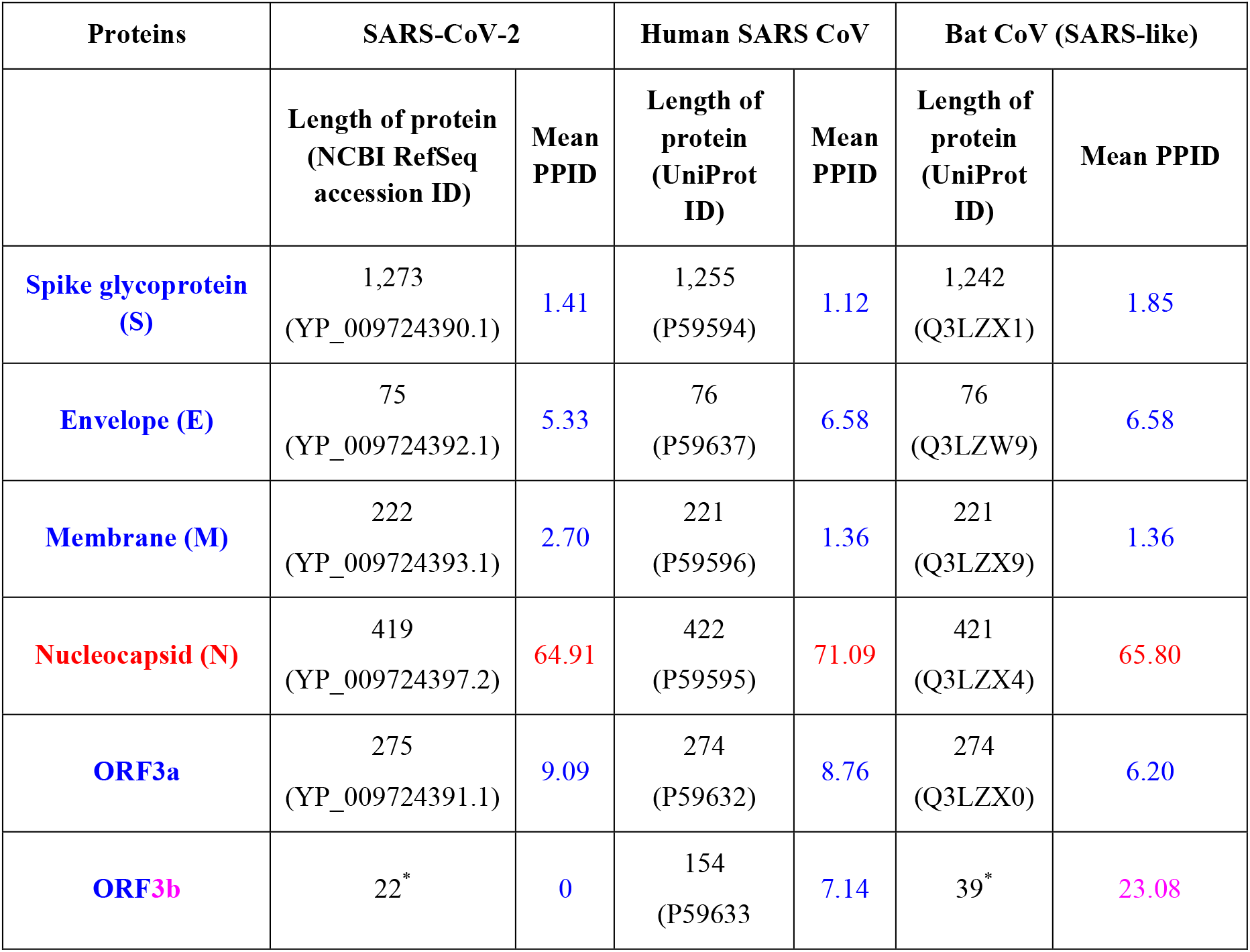
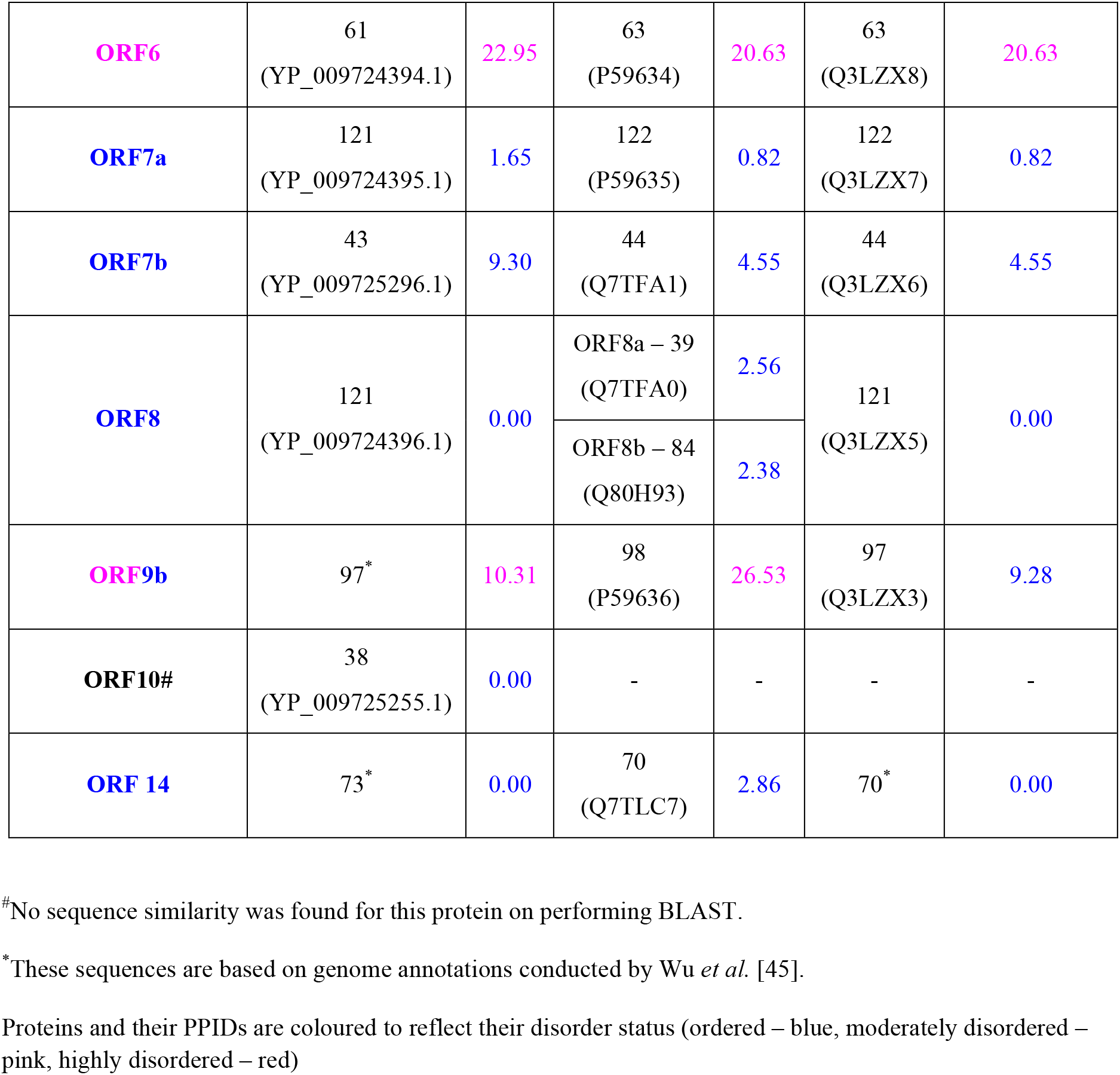
Mean PPID scores of structural and accessory proteins from SARS-CoV-2, Human SARS, and Bat CoVs.

| Proteins | SARS-CoV-2 |  | Human SARS CoV |  | Bat CoV (SARS-like) |  |
| --- | --- | --- | --- | --- | --- | --- |
|  | Length of protein<br>(NCBI RefSeq<br>accession ID) | Mean<br>PPID | Length of<br>protein<br>(UniProt<br>ID) | Mean<br>PPID | Length of<br>protein<br>(UniProt<br>ID) | Mean PPID |
| **Spike glycoprotein**<br>**(S)** | 1,273<br>(YP_009724390.1) | 1.41 | 1,255<br>(P59594) | 1.12 | 1,242<br>(Q3LZX1) | 1.85 |
| **Envelope (E)** | 75<br>(YP_009724392.1) | 5.33 | 76<br>(P59637) | 6.58 | 76<br>(Q3LZW9) | 6.58 |
| **Membrane (M)** | 222<br>(YP_009724393.1) | 2.70 | 221<br>(P59596) | 1.36 | 221<br>(Q3LZX9) | 1.36 |
| **Nucleocapsid (N)** | 419<br>(YP_009724397.2) | 64.91 | 422<br>(P59595) | 71.09 | 421<br>(Q3LZX4) | 65.80 |
| **ORF3a** | 275<br>(YP_009724391.1) | 9.09 | 274<br>(P59632) | 8.76 | 274<br>(Q3LZX0) | 6.20 |
| **ORF3b** | 22* | 0 | 154<br>(P59633) | 7.14 | 39* | 23.08 |
| **ORF6** | 61<br>(YP_009724394.1) | 22.95 | 63<br>(P59634) | 20.63 | 63<br>(Q3LZX8) | 20.63 |
| **ORF7a** | 121<br>(YP_009724395.1) | 1.65 | 122<br>(P59635) | 0.82 | 122<br>(Q3LZX7) | 0.82 |
| **ORF7b** | 43<br>(YP_009725296.1) | 9.30 | 44<br>(Q7TFA1) | 4.55 | 44<br>(Q3LZX6) | 4.55 |
| **ORF8** | 121<br>(YP_009724396.1) | 0.00 | ORF8a – 39<br>(Q7TFA0) | 2.56 | 121<br>(Q3LZX5) | 0.00 |
|  |  |  | ORF8b – 84<br>(Q80H93) | 2.38 |  |  |
| **ORF9b** | 97* | 10.31 | 98<br>(P59636) | 26.53 | 97<br>(Q3LZX3) | 9.28 |
| **ORF10#** | 38<br>(YP_009725255.1) | 0.00 | - | - | - | - |
| **ORF 14** | 73* | 0.00 | 70<br>(Q7TLC7) | 2.86 | 70* | 0.00 |

#No sequence similarity was found for this protein on performing BLAST.
\*These sequences are based on genome annotations conducted by Wu *et al.* [45].

### Per-residue predictions of intrinsic disorder predisposition

For the prediction of the intrinsic disorder predisposition of CoV proteomes, we have used multiple predictors, such as members of the PONDR^®^ (Predictor of Natural Disordered Regions) family including PONDR^®^VLS2 [29], PONDR^®^VL3 [30], PONDR^®^FIT [31], and PONDR^®^ VLXT [32], as well as the IUPred platform for predicting long (≥30 residues) and short IDPRs (,30 residues) [33]. These computational tools predict residues/regions, which do not have the tendency to form an ordered structure. Residues with disorder scores exceeding the threshold value of 0.5 are considered as intrinsically disordered residues, whereas residues with the predicted disorder scores between 0.2 and 0.5 are considered flexible. Complete predicted percent of intrinsic disorder (PPID) in a query protein was calculated for every protein of all the three viruses from outputs of six predictors. The detailed methodology has been given in our previous reports [34,35].

### Combined CH-CDF analysis to predict disorder predisposition of proteins

Charge hydropathy plot [36] and PONDR^®^ VLXT-based cumulative distribution function are two binary predictors of disorder (i.e., tool evaluating entire protein as mostly ordered or mostly disordered), which are available on the PONDR web server (https://www.pondr.com/). Combining the result from these binary predictors helps one to classify the proteins into different groups, depending on the flavours of their global disorder [37].

### Molecular recognition features (MoRFs) determination in CoV proteomes

The authentic online bioinformatics predictors that use a different set of algorithms for the prediction of MoRFs were used: These include MoRFchibi_web [38], ANCHOR [39,40], MoRFPred [41], and DISOPRED3 [42]. The protein residues with ANCHOR, MoRFPred, and DISOPRED3 score above the threshold value of 0.5 and MoRFchibi_web score above the threshold value of 0.725 are considered MoRF regions.

### Identification of DNA and RNA binding regions in CoV proteomes

Often IDPs and IDPRs facilitate interactions with RNAs and DNAs and regulates many cellular functions [43]. Thus, for predicting the DNA binding residues in CoV proteins, we have used two online servers: DRNAPred [19] and DisoRDPbind [43]. For RNA binding residues, we used Pprint (Prediction of Protein RNA-Interaction) [44] and DisoRDPbind [43].

## Results and Discussion

### Comprehensive computational analysis of the intrinsic disorder in proteins from SARS-CoV-2, human SARS and bat CoV (SARS-like)

The mean values of the predicted percentage of intrinsic disorder scores (mean PPIDs), that were obtained by averaging the predicted disorder scores from six disorder predictors (**Supplementary Table 1–3**) for each protein of SARS-CoV-2 as well as Human SARS, and Bat CoV are represented in **Table 1**.

**Figures 2A, 2B**, and **2C** are 2D-disorder plots generated for SARS-CoV-2, Human SARS and Bat CoV proteins, respectively, and represent the PPID_PONDR-FIT_ *vs*. PPID_Mean_ plots. Based on their predicted levels of intrinsic disorder, proteins can be classified as highly ordered (PPID < 10%), moderately disordered (10% ≤ PPID < 30%) and highly disordered (PPID ≥ 30%) [46]. From the data in **Table 1, Figures 2A, 2B**, and **2C**, as well as the PPID based classification, we conclude that the Nucleocapsid protein from all three strains of coronavirus possesses the highest percentage of the disorder and is classified as highly disordered protein. ORF3b protein in Bat Cov, ORF6 protein in SARS-CoV-2, Human SARS, and Bat CoV, and ORF9b protein in Human SARS and SARS CoV belong to the class of moderately disordered proteins. While the structured proteins, namely, Spike glycoprotein (S), an Envelope protein (E) and Membrane protein (M) as well as accessory proteins ORF3a, ORF7a, ORF8 (ORF8a and ORF8b in case of Human SARS) of all three strains of coronavirus are ordered proteins. ORF14 and ORF10 proteins also belong to the class of ordered proteins.

**Figure 2.**
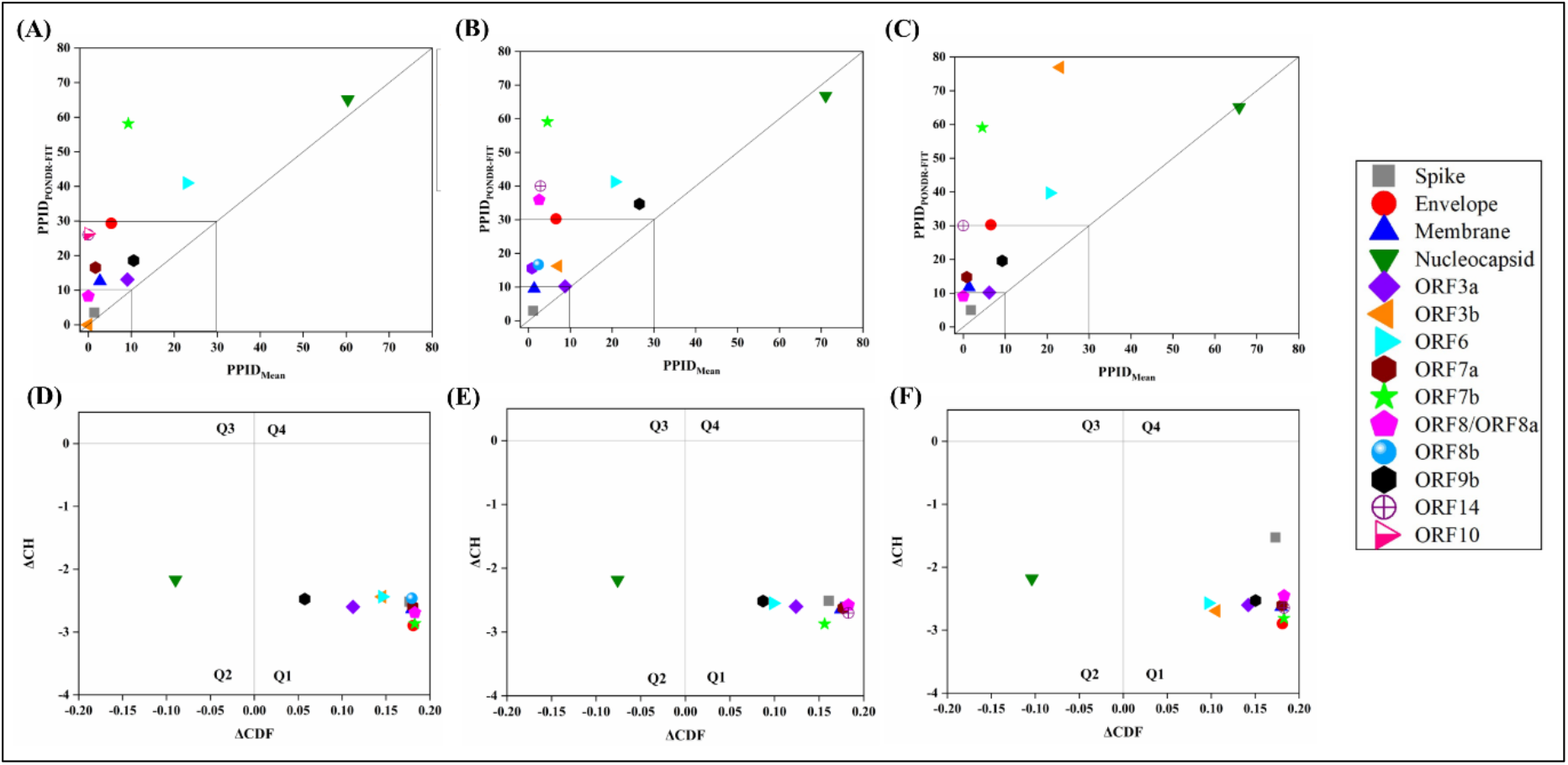
Analysis of overall disorder status of proteins of SARS-CoV–2, Human SARS, and Bat CoV (SARS-like): 2D plot representing PPID_PONDR-FIT_ *vs*. PPID_Mean_ in **(A)** SARS-CoV-2 **(B)** Human SARS and **(C)** Bat CoV. In CH-CDF plot of the proteins of **(D)** SARS-CoV-2 **(E)** Human SARS and **(F)** Bat CoV, the Y coordinate of each protein spot signifies distance of corresponding protein from the boundary in CH plot and the X coordinate value corresponds to the average distance of the CDF curve for the respective protein from the CDF boundary.

In order to further investigate the nature of the disorder in proteins of SARS-CoV-2, Human SARS, and Bat CoV, we utilized the combined CH-CDF tool that uses the outputs of two binary classifiers of disorder, Charge hydropathy (CH) plot and Cumulative distribution function (CDF) plot. This helped in retrieving more detailed characterization of the global disorder predisposition of the query proteins and their classification according to the disorder “favours”. The CH plot is a linear classifier that differentiates between proteins that are predisposed to possess extended disordered conformations that include random coils and pre-molten globules from proteins that have compact conformations (ordered proteins and molten globule-like proteins). The other binary predictor, CDF is a nonlinear classifier that uses the PONDR^®^VLXT scores to discriminate ordered globular proteins from all disordered conformations, which include native molten globules, pre-molten globules, and random coils. The CH-CDF plot can be divided into four quadrants: Q1 (bottom right quadrant) is an area of CH-CDF phase space that is expected to include ordered proteins; Q2 (bottom left quadrant) includes proteins predicted to be disordered by CDF and compact by CH (i.e., native molten globules and hybrid proteins containing high levels of both ordered and disordered regions); Q3 (top left quadrant) contains proteins that are predicted to be disordered by both CH and CDF analysis (i.e., highly disordered proteins with the extended disorder); and Q4 (top right quadrant) possesses proteins disordered according to CH but ordered according to CDF analysis [34]. **Figures 2D, 2E** and **2F** represent the CH-CDF analysis of proteins of SARS-CoV-2, Human SARS, and Bat CoV and show that all the proteins are located within the two quadrants Q1 and Q2. The CH-CDF analysis leads to the conclusion that all proteins of SARS-CoV-2, Human SARS, and Bat CoV are ordered except Nucleocapsid protein, which is predicted to be disordered by CDF but ordered by CH and hence lies in Q2.

Molecular recognition features (MoRFs) are short interaction-prone disordered regions found within IDPs/IDPRs that commence a disorder-to-order transition upon binding to their partners [47,48]. These regions are important for protein-protein interactions and may initiate an early step in molecular recognition [48]. In this study, we have analyzed and compared MoRFs (protein-binding regions) in SARS-CoV-2 with Human SARS and Bat CoV. The results of this analysis are summarized in **Table 2**, which clearly shows that most of the SARS-CoV-2 proteins contain at least one MoRF, indicating that disorder does play an important role in the functionality of these viral proteins. All of the SARS-CoV-2 proteins have been predicted to contain MoRFs except ORF7b and Nsp13 proteins for which none of the predictors have located any MoRF. MoRFs in Human SARS and Bat CoV proteomes are listed in **Supplementary Tables 7 and 8**. Similar to SARS-CoV-2 proteome, Bat CoV proteins ORF7b, and Nsp13 were not predicted to have any MoRF by any of the servers used. In Human SARS proteome the proteins ORF7b, Nsp13, Nsp2, and Nsp15 proteins does not show the presence of any MoRF. Interestingly, structural N protein from SARS-CoV-2, Human SARS, and Bat CoV shows high number of short and long MoRF regions, signifying its central role in virus pathogenesis.

**Table 2:** Predicted MoRFs regions in SARS-CoV-2 proteins

| Proteins | MoRFchibi_web | ANCHOR | MoRFPred | DISOPRED3 |
| --- | --- | --- | --- | --- |
| **Spike glycoprotein (S)** | 1265-1272 | - | 2-6, 819-823 | 1-10 |
| **Envelope (E)** | 46-75 | - | - | 26-30 |
| **Membrane (M)** | 187-221 | - | 214-222 | 117-132 |
| **Nucleocapsid (N)** | 1-20, 82-98, 102-118, 403-409 | 1-75, 104-113, 152-185, 201-236, 264-310, 347-361, 388-419 | 11-21, 48-58, 106-115, 218-228, 268-274, 354-358, 397-408 | 1-10,26-47,179-208,230-249,366-378,380-385,413-419 |
| **ORF 3a** | 1-6, 261-268 | - | 7-12, 126, 230-233, 259-263 | 1-19 |
| **ORF 3b** | NA | - | NA | - |
| **ORF6** | 14-21, 26-61 | - | - | - |
| **ORF7a** | 29-34, 39-51, 77-89 | - | - | 1-10 |
| **ORF7b** | - | - | - | - |
| **ORF8** | 1-5, 26-52, 69-91 | - | - | 1-10 |
| **ORF9b** | 3-29, 34-62, 79-84, | - | 89-97 | - |
| **ORF10** | 3-7 | - | - | - |
| **ORF14** | 1-45 | - | 6-12 | - |
| **Nsp1** | 64-75, 93-97, 99-179 | 173-180 | 137-145, 172-179 | 1-6 |
| **Nsp2** | 1-7 |  | - | - |
| **Nsp3** | - | 105-171, 189- | 101-107, 184-194, | - |
|  |  | 196 | 536-541, 980-984 |  |
| **Nsp4** | - | - | 495-500 | 1-25 |
| **Nsp5** | 3-8 | - | - | - |
| **Nsp6** | - | - | - | 1-19 |
| **Nsp7** | 1-30, 39-58, 65-83 | - | - | - |
| **Nsp8** | 181-185 | - | 89-93 | - |
| **Nsp9** | 40-44, 51-56, 62-77, 83-102 | - | - | - |
| **Nsp10** | 25-32, 91-99, 133-138 | - | 11-18 | - |
| **Nsp12** | 31-36 | - | 11-15 | 1-6 |
| **Nsp13** | - | - | - | - |
| **Nsp14** | - | - | 8-13, 441-445 | - |
| **Nsp15** | - | - | 9-13 | - |
| **Nsp16** | - | - | 151-156 | - |

### Nucleotide-binding propensity in proteins of coronaviruses

In addition to protein-protein interactions/protein-binding functions, IDPs and IDRs also mediates functions by facilitating their interactions with nucleotides such as DNA and RNA [20,49]. Therefore, we have used a combination of two different online servers for locating protein residues that are showing the propensity to bind with DNA as well as RNA. Nucleotide-binding residues in proteins of three studied coronaviruses are listed in **Supplementary Table 9-11**. These results mostly contain the residues in viral proteins that suitably bind viral RNA and host DNA. Interestingly, all the viral proteins of SARS-CoV-2, Human SARS, and Bat CoV have shown the propensity to bind to nucleic acid (DNA/RNA). In particular, structural (S, M, and N proteins) and non-structural (Nsp 2, 3, 4, 5, 6, 12, 13, 14, 15, and 16) proteins of all three viruses displays large number RNA binding residues. However, ORF3a, ORF3b, ORF6, ORF7a, ORF7b, ORF8, ORF9b, ORF10, and ORF14 shows less RNA binding and more DNA binding residues.

### Intrinsic disorder analysis of structural proteins of Coronaviruses

Coronaviruses encode four structural proteins, namely, Spike (S), Envelope (E) glycoprotein, Membrane (M), and Nucleocapsid (N) proteins, which are translated from the last ~10kb nucleotides and form the outer cover of the CoVs, encapsulating their single-stranded genomic RNA.

#### Spike (S) glycoprotein

S protein is a large multifunctional protein forming the exterior of the CoV particles [50,51]. It forms surface homotrimers and contains two distinct ectodomain regions known as S1 and S2. In some CoVs, the S protein is actually cleaved into these subunits, which are joined non-covalently, whereas an additional proteolytic cleavage within the N-terminal part of the S2 subunit that takes place upon virus endocytosis generates spike proteins S2’. Subunit S1 initiates viral infection by binding to the host cell receptors, S2 acts as a class I viral fusion protein that mediates fusion of the virion and cellular membranes and thereby promotes the viral entry into the host cells, whereas S2’ serves as a viral fusion peptide [52,53]. Spike binds to the virion M protein through its C-terminal transmembrane region [54]. Belonging to a class I viral fusion protein, S protein binds to specific surface receptor angiotensin-converting enzyme 2 (ACE2) on host cell plasma membrane through its N-terminal receptor-binding domain (RBD) and mediates viral entry into host cells [55].

The S protein consists of an N-terminal signal peptide, a long extracellular domain, a single-pass transmembrane domain, and a short intracellular domain [56]. A 3.60 Å resolution structure (PDB ID: 6ACC) of S protein from Human SARS complexed with its host binding partner ACE2 has been obtained by cryo-electron microscopy (cryo-EM). In this PDB structure, few residues (1-17, 240-243, 661-673, 812-831 and 1120-1203) are missing [57], suggesting their flexible nature. Also, the structure of the S protein (3.5 Å) from SARS-CoV-2 has been recently deduced by Wrapp *et. al.* using electron microscopy (PDB ID: 6VSB) [58] (**Figure 3A**). In this structure, residues 1-26, 67-78, 96-98, 143-155, 177-186, 247-260, 329-334, 444-448, 455-490, 501-502, 621-639, 673-686, 812-814, 829-851, 1147-1288 were missing, suggesting that the corresponding regions of a protein are characterized by high conformational flexibility. The biophysical analysis reported in previous study has also revealed that the S protein from SARS-CoV-2 has a higher binding affinity to ACE2 than S protein from Human SARS [58].

**Figure 3.**
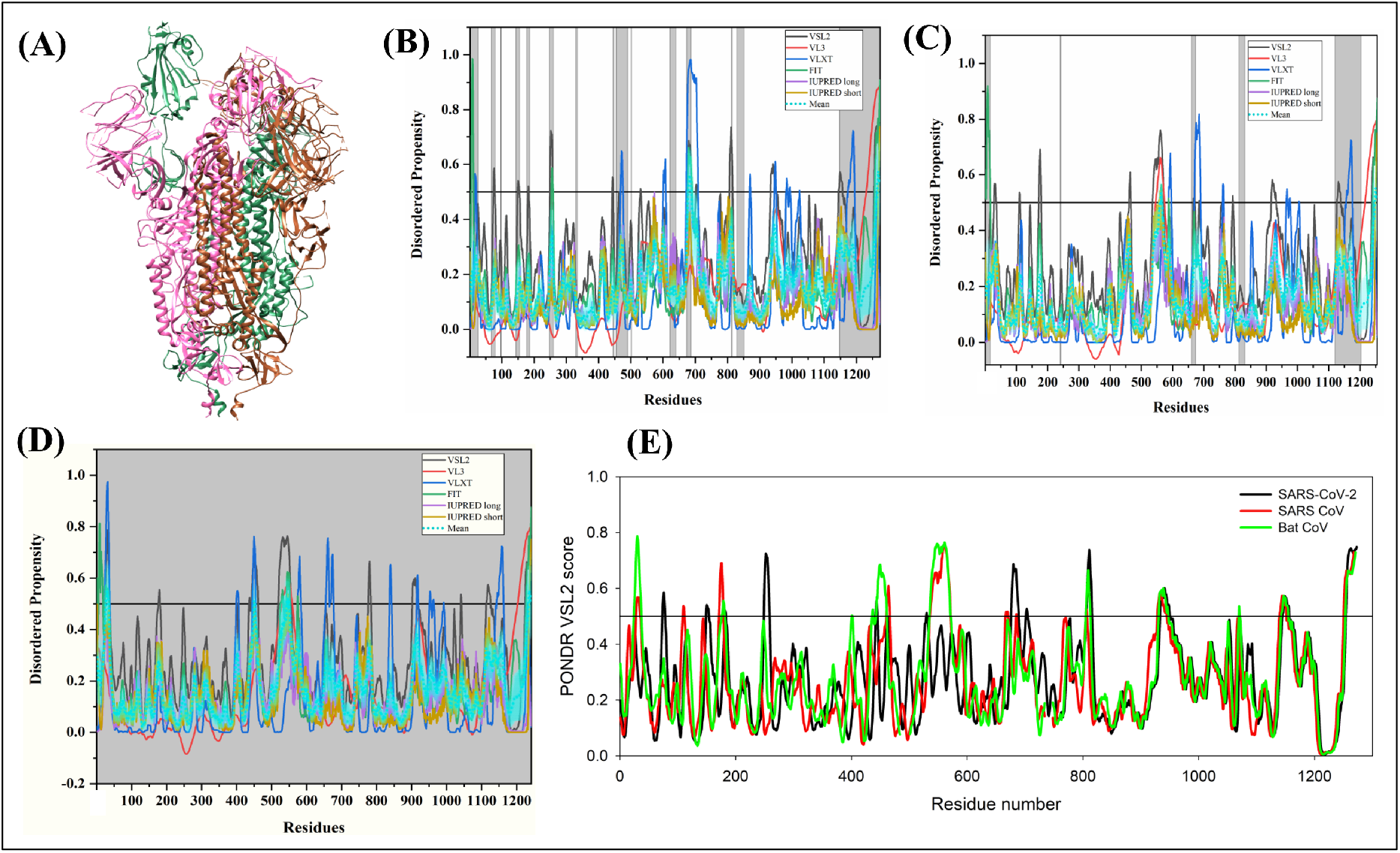
Structure and intrinsic disorder propensity of Spike glycoprotein from CoVs. **(A)** A 3.50 Å resolution structure of spike glycoprotein from SARS-CoV-2 obtained by cryo-EM (PDB ID: 6VSB). This homotrimeric structure includes three chains, A (violet-red), B (dark khaki), and C (turquoise). Evaluation of intrinsic disorder predisposition of S protein from SARS-CoV-2 **(B)**, Human SARS **(C)**, and Bat CoV **(D)**. Graphs in **(B), (C)** and **(D)** show the disorder profiles generated by PONDR^®^ VSL2 (black line), PONDR^®^ VL3 (red line), PONDR^®^ VLXT (blue line), PONDR^®^ FIT (green line), IUPred long (purple line) and IUPred short (golden line). The mean disorder propensity which has been calculated by averaging the disorder scores from all six predictors is represented by a short-dot line (sky-blue line) in the graph. The light sky-blue shadow region signifies the mean error distribution. The residues missing in the PDB structure or the residues for which PDB structure is unavailable are represented by the grey-coloured area in the corresponding graphs. **(E)** Aligned disorder profiles generated for spike glycoprotein from SARS-CoV-2 (black line), Human SARS (red line), and Bat CoV (green line) based on the outputs of the PONDR^®^ VSL2.

MSA analysis among all three coronaviruses demonstrates that S protein of SARS-CoV-2 has a 77.71% sequence identity with Bat CoV and 77.14% identity with Human SARS **(Supplementary Figure S1A)**. All three S proteins are found to have a conserved C-terminal region. However, the N-terminal regions of S proteins display noticeable differences. Given that there is significant sequence variation RBD located at the N-terminal region of S protein, this might be the reason behind variation in its virulence and its receptor-mediated binding and entry into the host cell.

According to our intrinsic disorder propensity analysis, S protein from all three CoVs analysed in this study are highly structured, as their predicted disorder propensity lies below 10% (**Table 1**). In fact, the mean PPID scores of SARS-CoV-2, Human SARS CoV, and Bat CoV are calculated to be 1.41%, 1.12%, and 1.85%, respectively. **Figures 3B, 3C**, and **3D** represent the intrinsic disorder profiles of S proteins from SARS-CoV-2, Human SARS and Bat CoV obtained from six disorder predictors. Finally, **Figure 3E** shows aligned disorder profiles of S proteins from these CoVs and illustrates remarkable similarity in their disorder propensity, especially in the C-terminal region.

It is of interest to map known functional regions of S proteins to their corresponding disorder profiles. The maturation of S protein requires specific posttranslational modification (PTM), proteolytic cleavage that happens at two stages. First, host cell furin or another cellular protease nicks the S precursor to generate S1 and S2 proteins, whereas the second cleavage that takes place after the viral attachment to host cell receptors leads to the release of a fusion peptide generating the S2’ subunit. In Human SARS CoV, the first and second cleavage site is located at residues R_667_ and R_797_, respectively, whereas in Bat CoV, the corresponding cleavage sites are residues R_654_ and R_784_. As it follows from **Figure 3**, these cleavage sites are located within the IDPRs. In Human SARS CoV S protein, fusion peptide (residues 770-788) is located within a flexible region, is characterized by the mean disorder score of 0.232±0.053. Similarly, in Bat CoV S protein, fusion peptide (residues 757-775) has a mean disorder score of 0.320±0.046. S protein contains two heptad repeat regions that form coiled-coil structure during viral and target cell membrane fusion, assuming a trimer-of-hairpins structure needed for the functional positioning of the fusion peptide. In Human SARS CoV S protein, heptad repeat regions are formed by residues 902-952 and 1145-1184, which have mean disorder scores of 0.458±0.067 and 0.353±0.062, respectively. The analogous situation is observed for the S protein from Bat CoV, where these heptad repeat regions are positioned at residues 889-939 (0.44±0.11) and 1132-1171 (0.353±0.062). Another functional region found in S proteins is the receptor-binding domain (residues 306-527 and 310-514 in Human SARS CoV and Bat CoV, respectively) containing a receptor-binding motif responsible for interaction with human ACE2. In human S protein of Human SARS CoV this motif (residues 424-494) is not only characterized by structural flexibility, possessing a mean disorder score of 0.30±0.16, but also contains a disordered region (residues 461-466). Since S protein is known as spike glycoprotein, it contains numerous glycosylation sites. Due to rather close similarity of disorder profiles of S proteins analysed here, we can assume that all the aforementioned indications of the functional importance of disorder and flexible regions in S proteins from SARS CoV and Bat CoV are also applicable to SARS-CoV-2 S protein. Finally, **Table 2** shows that S protein from SARS-CoV-2 contain one MoRF region at its C-terminal (residues 1265-1272) by MoRFchibi_web, two MoRF regions ((residues 2-6) & (residues 819-823)) by MoRFPred, and one MoRF region at N-terminal (residues 1-10) by DISOPRED3. These results indicating that intrinsic disorder is important for its interaction with binding partners. Interestingly, the N-terminal region of S protein (residues 1-10) from all three viruses are observed to be a disorder-based protein binding region by two predictors (MoRFPred and DISOPRED3). N-terminal MoRF displays its role in viral interaction with host receptor and C-terminal MoRF displays its role in M protein interaction and viral assembly. Moreover, MoRF region mainly lies in the N- and C-terminal regions suggesting a possible role during cleavage as well. In addition to protein-binding regions, S protein also shows many nucleotide-binding residues. **Tables 9, 10, and 11** shows that numerous RNA binding residues predicted by PPRint in all three viruses and a single RNA binding residue were predicted by DisoRDPbind in human SARS. Further, DRNApred and DisoRDPbind predicted the presence of many DNA binding residues in S protein of all three viruses. These results signify the role of S protein functions related to molecular recognition (protein-protein interaction, RNA binding, and DNA binding) such as interaction with host cell membrane and further viral infection. Therefore, identified IDPs/IDPRs and residues/regions from S protein crucial for molecular recognition can be targeted for disorder-based drug discovery.

#### Envelope (E) small membrane protein

Envelope (E) protein is a small, multifunctional inner membrane protein that plays an important role in the assembly and morphogenesis of virions in the cell [59–61]. E protein consists of two ectodomains associated with N- and C-terminal regions, and a transmembrane domain. It homo-oligomerize to form pentameric membrane destabilizing transmembrane (TM) hairpins to form a pore necessary for its ion channel activity [62]. **Figure 4A** shows the NMR-structure (PDB ID: 2MM4) of Human SARS envelope glycoprotein of 8-65 residues [63].

**Figure 4.**
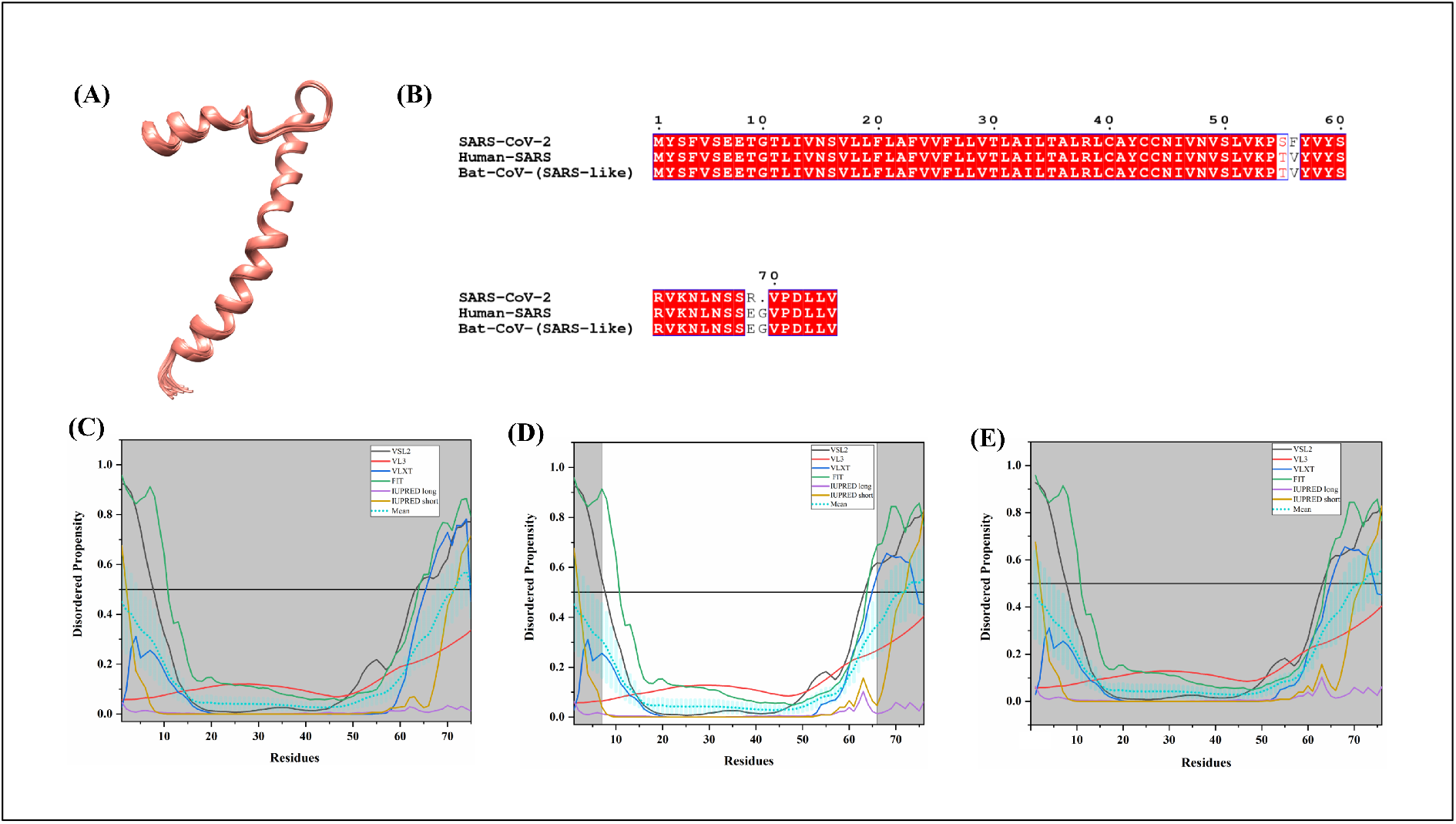
Analysis of structural features and intrinsic disorder predisposition of Envelope glycoprotein. **(A)** The solution NMR generated structure of envelope glycoprotein of Human SARS (PDB ID: 2MM4). The structure is shown for the region consists of residues 8-65. **(B)** MSA profile of envelope glycoproteins of SARS-CoV-2, Human SARS CoV and Bat CoV. Red shaded sequences represent identical regions of these proteins. (**C-E)** graph represents the intrinsic disorder profiles of E protein from SARS-CoV-2 **(C)**, Human SARS CoV **(D)**, and Bat CoV **(E)**. The colour schemes are the same as in the corresponding plots of **Figure 3**.

MSA results illustrate (**Figure 4B**) that this protein is highly conserved, with only three amino acid substitutions in E protein of SARS-CoV-2 conferring its 96% sequence similarity with Human SARS and Bat CoV. Bat CoV shares 100% sequence identity with Human SARS. Mean PPID calculated for SARS-CoV-2, Human SARS, and Bat CoV E proteins are 5.33%, 6.58%, and 6.58% respectively (**Table 1**). The E protein is found to have a reasonably well-predicted structure. Our predictions suggest that the residues of N- and C-terminals are displaying a higher tendency for the disorder. The last 18 hydrophilic residues (residues 59-76) have been reported to adopt a random-coil conformation with and without the addition of lipid membranes [64]. Literature suggests that the last four amino acids of the C-terminal region of E protein containing a PZD-binding motif are involved in protein-protein interactions with a tight junction protein PALS1. Our results support literature as we identified long N-terminal region of approximately 30 residues long as disorder-based protein binding region in all three viruses (see **Table 2, Supplementary Table 7** and **8**). PALS1 is involved in maintaining the polarity of epithelial cells in mammals [65]. Respective graphs in **Figures 4C, 4D**, and **4E** show the predicted intrinsic disorder profiles for E proteins of SARS-CoV-2, Human SARS, and Bat CoV. We speculate that the disordered region content may be facilitating the interactions with other proteins as well. In agreement with this hypothesis, **Table 2** shows that in E protein from SARS-CoV-2, the C-terminal domain serves as protein-binding region. We found that the residues from 45-75 is a long MoRF in E proteins of all three viruses as predicted by MoRFchibi_web (**Table 2, Supplementary Table 7** and **8**). As aforementioned, these randomly-coiled binding-residues at C-terminus may gain structure while assisting the protein-protein interaction mediated by E protein. One more MoRF region (residues 26-30) in the transmembrane domain was observed by DISOPRED3 in the E protein of all three viruses. Since these residues are the part of ion channel, we speculate that these residues do specific interactions and may be guiding the specifi functions of ion channel activity.

Few RNA binding residues by PPRint and DisoRDPbind and several DNA binding residues by DRNApred are predicted for E protein in all three viruses.

#### Membrane (M) glycoprotein

Membrane (M) glycoprotein plays an important role in virion assembly by interacting with the nucleocapsid (N) and E proteins [66–68]. Protein M interacts specifically with a short viral packaging signal containing coronavirus RNA in the absence of N protein, thereby highlighting an important nucleocapsid-independent viral RNA packaging mechanism inside the host cells [69]. It gains high-mannose N-glycans in ER, which are subsequently modified into complex N-glycans in the Golgi complex. Glycosylation of M protein is observed to be not essential for virion fusion in cell culture [70,71].

Cryo-EM and Tomography data indicate that M forms two distinct conformations, a compact M protein having high flexibility and low spike density, and an elongated M protein having a rigid structure and narrow range of membrane curvature [72]. Some regions of M glycoproteins might serve as important dominant immunogens. Although no structural information is available for the full-length M protein as of yet, a short peptide of the membrane glycoprotein (residues 88-96) from Human SARS CoV was co-crystallized with a complex between A-2 alpha chain of the HLA class I histocompatibility antigen and β2-microglobulin (PDB ID: 3I6G) [73]. **Figure 5A** shows that within this complex, the co-crystallized M protein region exists in an extended conformation.

**Figure 5.**
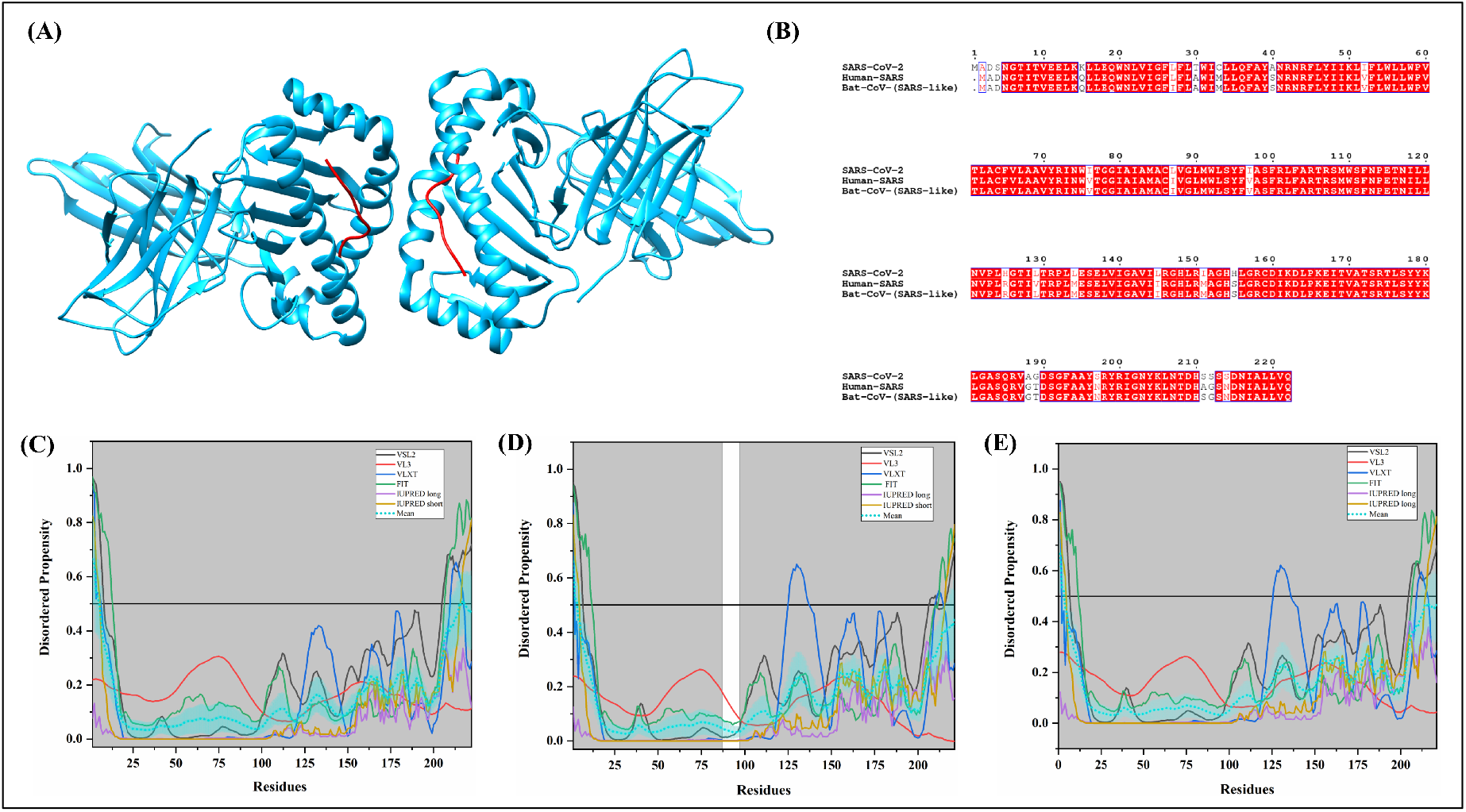
Analysis of intrinsic disorder propensity of Membrane glycoprotein. **(A)** A 2.20 Å resolution crystal structure of a complex between the A-2 alpha chain of HLA class I histocompatibility antigen, β2-microglobulin and the immunogen peptide of the membrane glycoprotein (9 amino acids) from Human SARS CoV (PDB ID: 3I6G). This complex crystallized is a dimer. Therefore, each member is present here in two copies. Chains corresponding to the fragment of the membrane glycoprotein (residues 88-96) are shown by red and pink colours, A-2 alpha chain and β2-microglobulin in this dimer of heterotrimers structure are shown as blue/ice-blue and tan/yellow surfaces. **(B)** MSA profile of Membrane glycoproteins of SARS-CoV-2, Human SARS and Bat CoV (SARS-like). Red shaded sequences represent identical regions of these proteins. **(C-E)** graphs represent the intrinsic disorder profiles of M protein from SARS-CoV-2, **(D)** Human SARS CoV, and **(E)** Bat CoV. The colour schemes are the same as in the corresponding plots of **Figure 3**.

M protein of SARS-CoV-2 has a sequence similarity of 90.1% with Bat CoV and 89.6% with Human SARS M proteins (**Figure 5B**). Our analysis revealed that the intrinsic disorder levels in M proteins of SARS-COV-2, Human SARS CoV, and Bat CoV are relatively low since these proteins show the PPID values of 2.70%, 1.36%, and 1.36% respectively. This is in line with the previous publication by Goh *et al.* on Human SARS HKU4 where they found the mean PPID of 4% using additional predictors such as TopIDP and FoldIndex along with the predictors used in our study [74]. **Figures 5C, 5D**, and **5E** represent per-residue disorder profiles generated for M proteins of SARS-CoV-2, Human SARS CoV, and Bat CoV and show that with the exception to their N- and C-terminal regions, these proteins are mostly ordered. The last 20 residues of MERS-CoV M protein are important for intracellular trafficking and contains a determinant that localizes it into the Golgi network [75]. Our results in **Table 2** illustrates that the disordered C-tail of the M protein is predicted to have disorder based protein-binding region and therefore can serve as a binding site for its specific partner required for its localization inside the host cell. A long MoRF region (residues 186-220) at the C-terminal of M protein in all three viruses were observed by MoRFchibi_web. Two MoRF regions (one at N-terminus (residues 1-16) and one at C-terminus (residues 205-221)) was observed by DISOPRED3 in human SARS and Bat CoV. However, single MoRF (residues 117-132) observed in SARS-CoV-2 by DISOPRED3. MoRFPred also predicts a short MoRF at C-terminus of SARS-CoV-2 (residues 214-222), Human SARS (residues 214-221), and Bat CoV (residues 211-221) (**Table 2, Supplementary Tables 7 and 8**). Furthermore, the M protein from all three viruses displays strong tendency to bind with RNA (as predicted by PPRint and DisoRDPbind) and DNA (as predicted by DRNApred and DisoRDPbind) (see **Supplementary Tables 9, 10, and 11**). Our understanding on M protein of CoVs (IDPs and MoRF at C-terminus and molecular recognition) elucidates its crucial role in interaction with the N and E proteins for viral assembly.

#### Nucleocapsid (N) protein

Nucleocapsid (N) protein is one of the major viral proteins playing several significant roles in transcription, and virion assembly of coronaviruses [76]. It binds to viral genomic RNA forming a ribonucleoprotein core required for the RNA encapsidation during viral particle assembly [77]. SARS-CoV virus-like particles (VLPs) formation has been reported to depend upon either M and E proteins or M and N proteins. For the effective production and release of VLPs, co-expression of E or N proteins with M protein is necessary [78].

N protein of Human SARS consists of two structural domains, the N-terminal RNA-binding domain (NTD: 45-181 residues) and the C-terminal dimerization domain (CTD: 248-365 residues) with a disordered patch in between these domains. N protein has been demonstrated to bind viral RNA using both NTD and CTD [79]. **Figure 6A1** displays the NMR solution structure of the NTD of Human SARS CoV nucleocapsid protein (45-181 residues) (PDB ID: 1SSK) [80]. **Figure 6A2** shows an X-ray crystal structure of the CTD of Human SARS CoV nucleocapsid protein (270-366 residues) (PDB ID: 2GIB) [81]. A model of the domain organization of the N-protein from SARS-CoV-2 is shown in **Figure 6B**.

**Figure 6.**
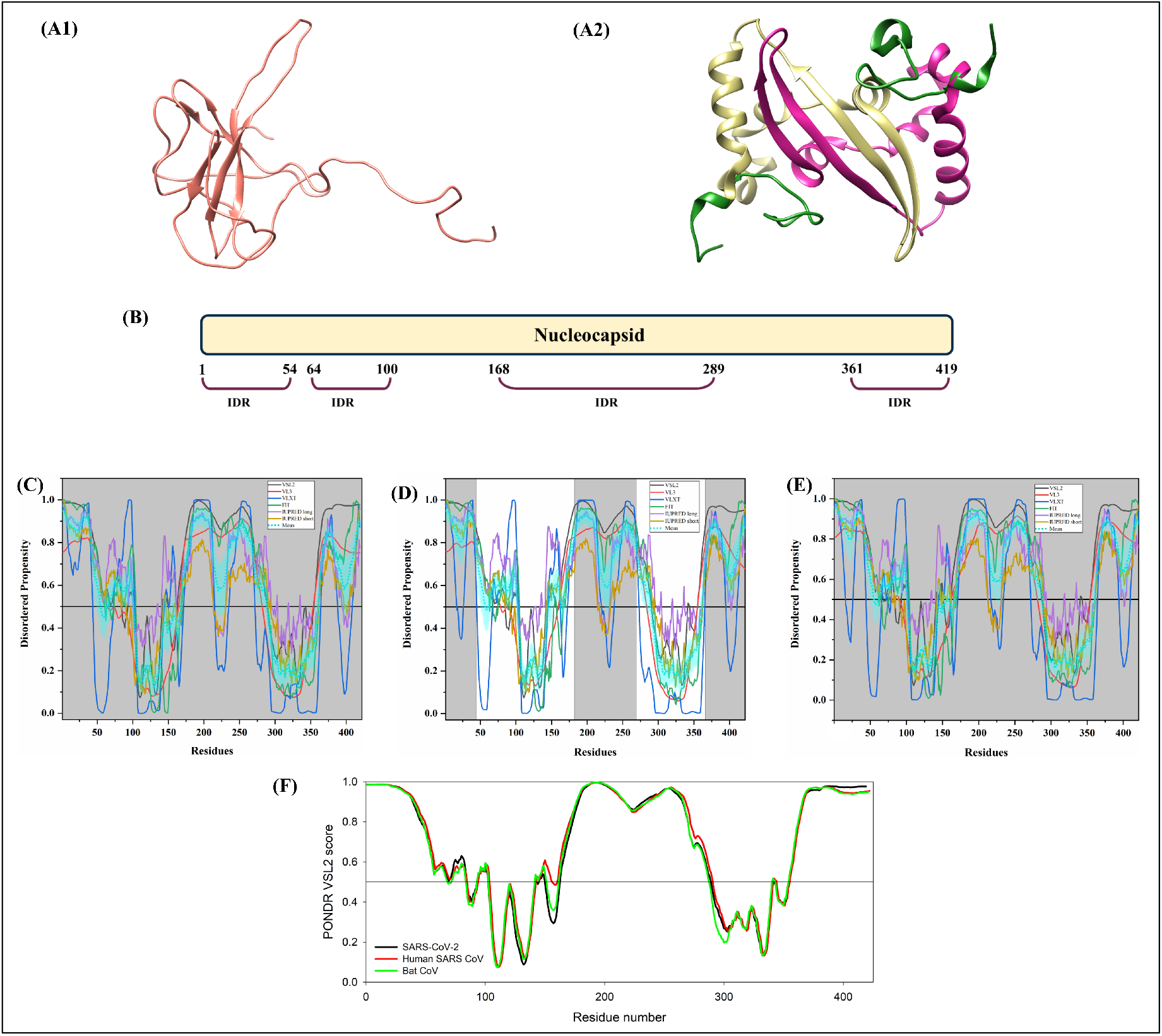
Analysis of structural properties and intrinsic disorder propensity of the coronaviral Nucleocapsid protein. **(A1)** NMR solution structure of the NTD (residues 45-181) of nucleocapsid from Human SARS CoV (PDB ID: 1SSK). **(A2)** X-ray crystal structure with the 1.75Å resolution of the CTD (residues 270-366) of nucleocapsid protein from Human SARS (PDB ID: 2GIB). The structure is a homodimer of chains A (violet-red) and B (dark khaki). Residues 270-289 and 362-366 have shown disorder propensity in the available structures (forest green colour). **(B)** Representation of the predicted disordered regions of SARS-CoV-2 on N protein. **(C-E)** graphs represent the intrinsic disorder profiles of N protein from **(C)** SARS-CoV-2, **(D)** Human SARS CoV, and **(E)** Bat CoV. The colour schemes are the same as in corresponding plots of **Figure 3**. **(F)** Aligned disorder profiles generated for N protein from SARS-CoV-2 (black line), Human SARS CoV (red line), and Bat CoV (green line) based on the outputs of the PONDR^®^ VSL2.

The 419 amino acid-long N protein of SARS-CoV-2 shows a percentage identity of 88.76% with N protein of Bat CoV N protein and 89.74% with Human SARS N protein (**Supplementary Figure S1B)**. Our analysis revealed that the N proteins of coronaviruses contain the highest levels of intrinsic disorder (see **Figure 6** and **Table 1**). In fact, N proteins from SARS-COV-2 Human SARS CoV, and Bat CoV are characterized by the mean PPID of 64.91%, 71.09%, and 65.80%, respectively. In accordance with the previously evaluated intrinsic disorder predisposition [74], N protein is highly disordered in all three SARS viruses analysed in this study (**Table 1**). Graphs in **Figures 6C, 6D**, and **6E** depict the disorder profiles of SARS-CoV-2, Human SARS CoV, and Bat CoV nucleocapsid proteins and show that their N- and C-terminal regions are completely disordered, and all three proteins also contain the central unstructured segment. As expected, the intrinsic disorder predisposition of the N protein of SARS-CoV-2 is remarkably similar to that for the N protein of Human SARS CoV as reported in a previous study [74]. This is further supported by **Figure 6F**, where PONDR^®^ VSL2-generated disorder profiles of these three proteins are overlapped to show almost complete coincidence of their major disorder-related features. It is clear that in N-proteins, the N- and C-termini and a log central segment are completely disordered. **Figure 6C** shows that in the N protein from SARS-CoV-2, residues 1-57, 64-102, 145-162, 166-289, and 362-422 are found to be disordered. Many of these residues are lying within the NTD and CTD regions, and which, due to their structural plasticity, were not crystallized in Human SARS CoV N protein. SARS-CoV-2 has a disordered segment from 168-289 residues while Human SARS has predicted to have an unstructured segment from 145-289 residues. Overall, all three N proteins are found to be highly disordered.

The N protein from Human SARS CoV has one phosphorylation site (residue S_177_) and several regions with compositional biases, such as Ser-rich (residues 181-213), Poly-Leu, Poly-Gln, and Ploy-Lys (residues 220-225, 240-245, and 370-376), all predicted to be disordered. Similarly, in N protein from Bat CoV, S_176_ is phosphorylated, and this protein has Ser-rich, Poly-Leu, and Ploy-Lys regions (residues 176-206, 219-224, and 369-375, respectively), all of which are disordered. It has been reported to interact using the central disordered region with M protein, hnRNP A1, and self N-N interaction [82–84]. The middle flexible region is also responsible for its RNA-binding activity [85]. Deletion of 184-196 residues, 169-308 residues, 161-210 residues of N abolishes its multimerization, RNA-binding capacity, and hnRNP A1 interactions respectively. **Supplementary Table 7** and **8**, and **Table 2** shows that N protein is heavily decorated with numerous MoRFs, suggesting that this protein is a promiscuous binder. Long disorder-based protein bonding regions at N- and C-terminus of N protein of all three viruses were observed by all four predictors (MoRFchibi_web, ANCHOR, MoRFPred, and DISOPRED3). Indeed, this is the single protein where we found many MoRFs as compared with the other structural, non-structural and accessory proteins of CoVs. The MoRFs present in these regions may mediate the above-mentioned interactions of N proteins. **Figure 6A2** represents another important disorder-related functional feature of the N protein. In fact, the CTD homodimer shown there is characterized by highly intertwined morphology, which is typically a result of binding-induced folding [86–88], indicating that a very significant part of CTD gains structure during dimerization. We identified numerous RNA binding residues in all three viruses using PPRint server. This finding supports the function of N protein as it interacts with genomic RNA for a ribonucleoprotein core formation which is crucial step for RNA encapsidation during viral particle assembly. In addition, DRNApred and DisoRDPbind predicts multiple DNA binding residues for N protein in SARS-CoV-2, Human SARS, and Bat CoV. The long flexible (IDPRs) regions at N and C-terminus of SARS-CoV-2 have long protein-binding as well as nucleotide-binding regions that may have important role in its interaction with viral RNA. These flexible regions can be targeted to inhibit interaction of N protein with viral genomic RNA.

### Intrinsic disorder analysis of accessory proteins of Coronaviruses

Literature suggests that some viral proteins are translated from the genes interspersed in between the genes of structural proteins. These proteins are known as accessory proteins, and many of them are proposed to be involved in viral pathogenesis [89].

#### Proteins ORF3a and ORF3b

ORF3a is a multifunctional protein with the molecular weight of ~31 kDa that has been found to localize in different organelles inside the host cells. Also referred to as *U274*, *X1*, and *ORF3*, the gene for this protein is present between the *S* and *E* genes of the SARS-CoV genome [90–92]. The homo-tetrameric complex of ORF3a has been demonstrated to form a potassium-ion channel on the host cell plasma membrane [93]. It performs a major function during virion assembly by co-localizing with E, M, and S viral proteins [94,95]. ORF3b protein can be found in the cytoplasm, nucleolus, and outer membrane of mitochondria of the host cells [96,97]. In Huh 7 cells, its over-expression has been linked with the activation of AP-1 via ERK and JNK pathways [98]. Transfection of ORF3b-EGFP leads to cell growth arrest at the G0/G1 phase of Vero, 293, and COS-7 cells [99]. ORF3a induces apoptosis via caspase 8/9 directed mitochondrial-mediated pathways, while ORF3b is reported to affect only the caspase 3-related pathways [100,101].

On performing MSA, results of which are shown in **Figure 7D**, we found that ORF3a protein from SARS-COV-2 is slightly evolutionary closer to the ORF3a of Bat CoV (73.36%) than to the ORF3a of Human SARS CoV (72.99%). Graphs in **Figures 7A, 7B**, and **7C** depict the propensity for disorder in ORF3a proteins of novel SARS-CoV-2, Human SARS CoV, and Bat CoV (SARS-like), respectively. Mean PPIDs in these ORF3a proteins are 9.1% (SARS-CoV-2), 8.8% (Human SARS), and 6.2% (Bat CoV (SARS-like)). ORF3a of SARS CoV-2 shows protein-binding regions at its N-terminus (by MoRFchibi_web (residues 1-6), MoRFPred (residues 7-12), and DISOPRED3 (residues 1-19)) and at C-terminus (by MoRFchibi_web (residues 261-268) and MoRFPred (residues 259-263)) (**Table 2**). Similarly, ORF3a of Human SARS and Bat CoV also shows MoRFs at N- and C-terminus with the help of MoRFchibi_web and MoRFPred (**Supplementary Tables 7 and 8**). These protein-binding regions in ORF3a may have role in its co-localization with E, M, and S viral proteins. Apart from MoRFs, it also displays several nucleotide-binding residues in all three viruses (see **Supplementary Tables 9, 10, and 11**). In fact, this represents maximum number of RNA and DNA binding residues as compared with all other accessory proteins. These results indicate that the IDPs/IDPRs of this protein could be utilized in molecular recognition (protein-protein, protein-RNA, and protein-DNA interaction).

**Figure 7.**
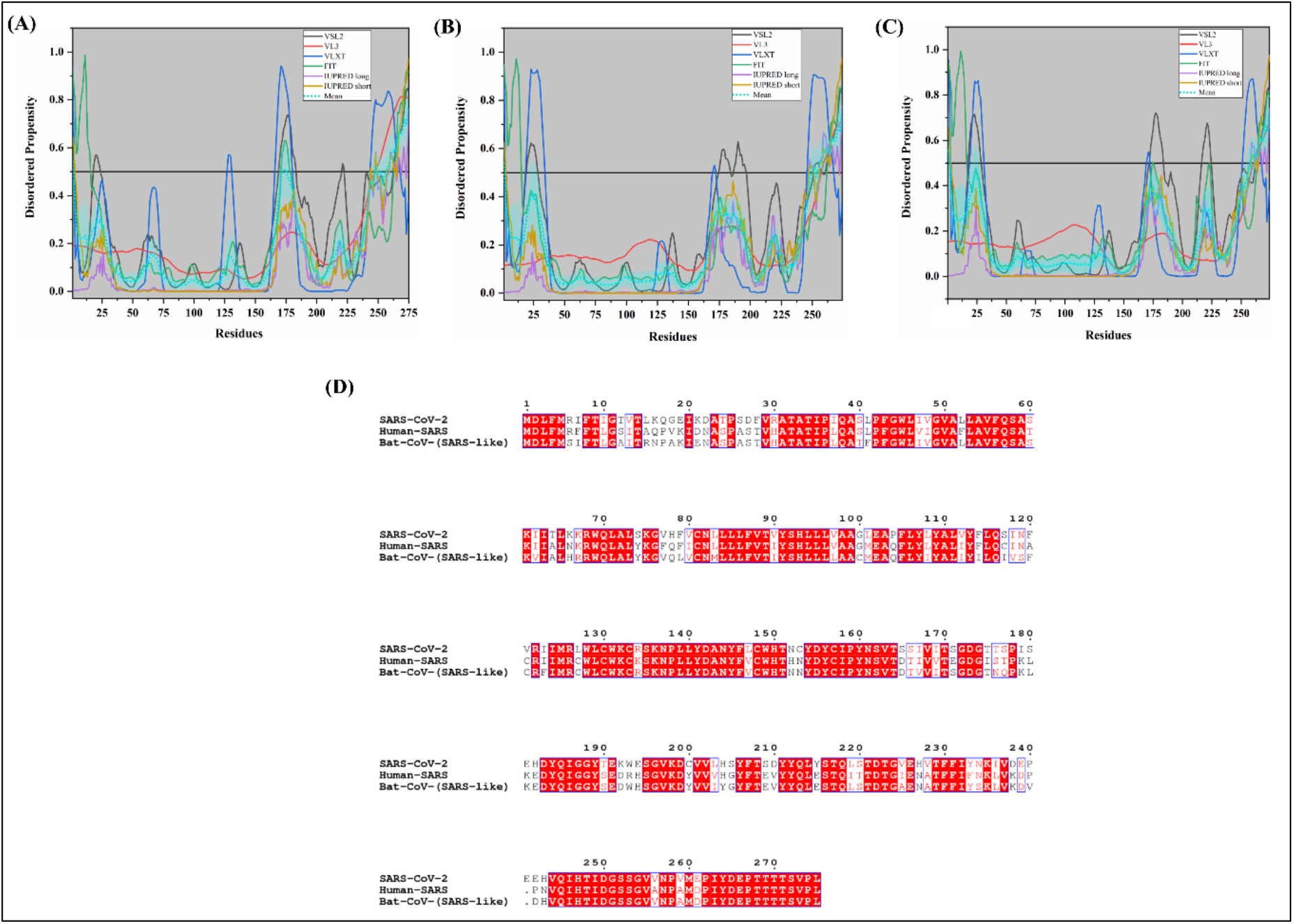
Analysis of intrinsic disorder propensity of ORF3a protein. **(A-C)** graphs represent the intrinsic disorder profiles of ORF3a protein from **(A)** SARS-CoV-2, **(B)** Human SARS CoV, and **(C)** Bat CoV. The colour schemes are the same as in the corresponding plots of **Figure 3. (D)** MSA profile of ORF3a proteins from SARS-CoV-2, Human SARS CoV, and Bat CoV. Red shaded sequences represent identical regions of these proteins, whereas red characters show similar residues.

According to the intrinsic disorder predisposition analysis of ORF3b proteins, their mean PPID values in SARS-CoV-2, Human SARS CoV, and Bat CoV are 0%, 7.1%, and 23.1% respectively, as represented in **Figures 8A, 8B**, and **8C**. MSA results (**Figure 8D**) demonstrate that ORF3b of SARS-CoV-2 is not closer to ORF3b protein of Human SARS and ORF3b protein of Bat-CoV, having a sequence similarity of only 54.6% and 59.1%, respectively. As we can see in **Table 2**, there is not a single MoRF found in ORF3b of SARS-CoV-2. However, for Human SARS we identified three MoRFs (residues 32-37, 41-70, and 125-153) and for Bat CoV one MoRF at N-terminus (residues 1-38) by MoRFchibi_web server.

**Figure 8.**
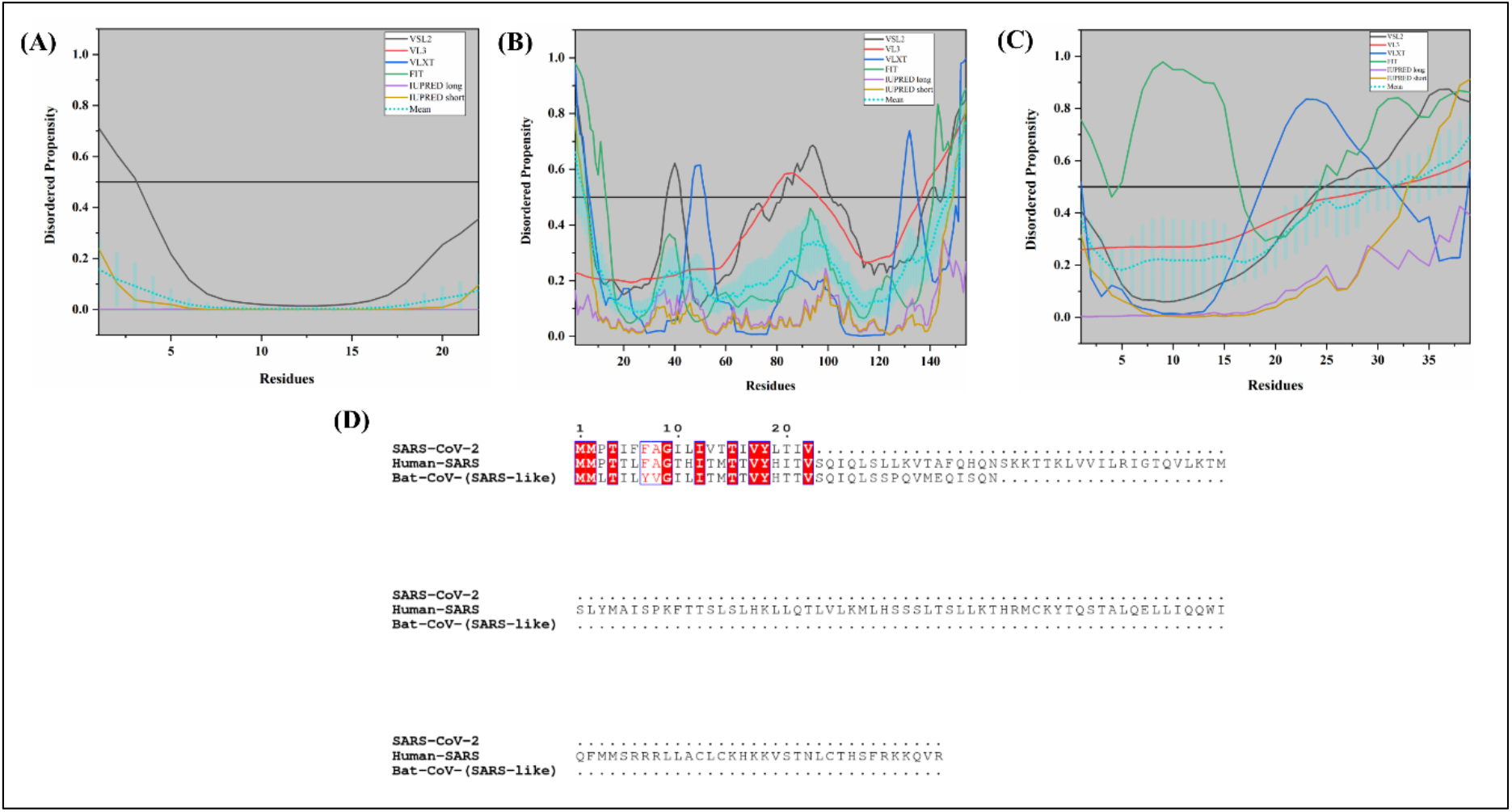
Analysis of intrinsic disorder propensity of ORF 3b protein of Human SARS CoV. **(A)** The graph represents the results of intrinsic disorder analysis in SARS-CoV-2, **(B)** Human SARS, and **(C)** Bat CoV. The colour schemes are the same as in the corresponding plots of **Figure 3. (D)** MSA profile of ORF3b proteins from SARS-CoV-2, Human SARS CoV, and Bat CoV. Red shaded sequences represent identical regions of these proteins, whereas red characters show similar residues.

#### Protein ORF6

ORF6 is a short coronavirus protein with just 63 residues. Also known as P6, this membrane-associated protein serves as an interferon (IFN) antagonist [102]. It downregulates the IFN pathway by blocking a nuclear import protein, karyopherin α2. Using its C-terminal residues, ORF6 disrupts karyopherin import complex in the cytosol and, therefore, hampers the movement of transcription factors like STAT1 into the nucleus [102,103]. It contains a YSEL motif near its C-terminal region, which functions in protein internalization from the plasma membrane into the endosomal vesicles [104]. Another study has also demonstrated the presence of ORF6 in endosomal/lysosomal compartments [104,105].

MSA results demonstrate that (**Figure 9D**), SARS-CoV-2 ORF6 is closer to ORF6 protein of Human SARS CoV, having a sequence similarity of 68.85% than to the ORF6 of Bat CoV (SARS-like) (67.21%). Novel SARS-CoV-2 ORF6 is predicted to be the second most disordered structural protein, with PPID of 22.95%, and with especially disordered C-terminal region.

**Figure 9.**
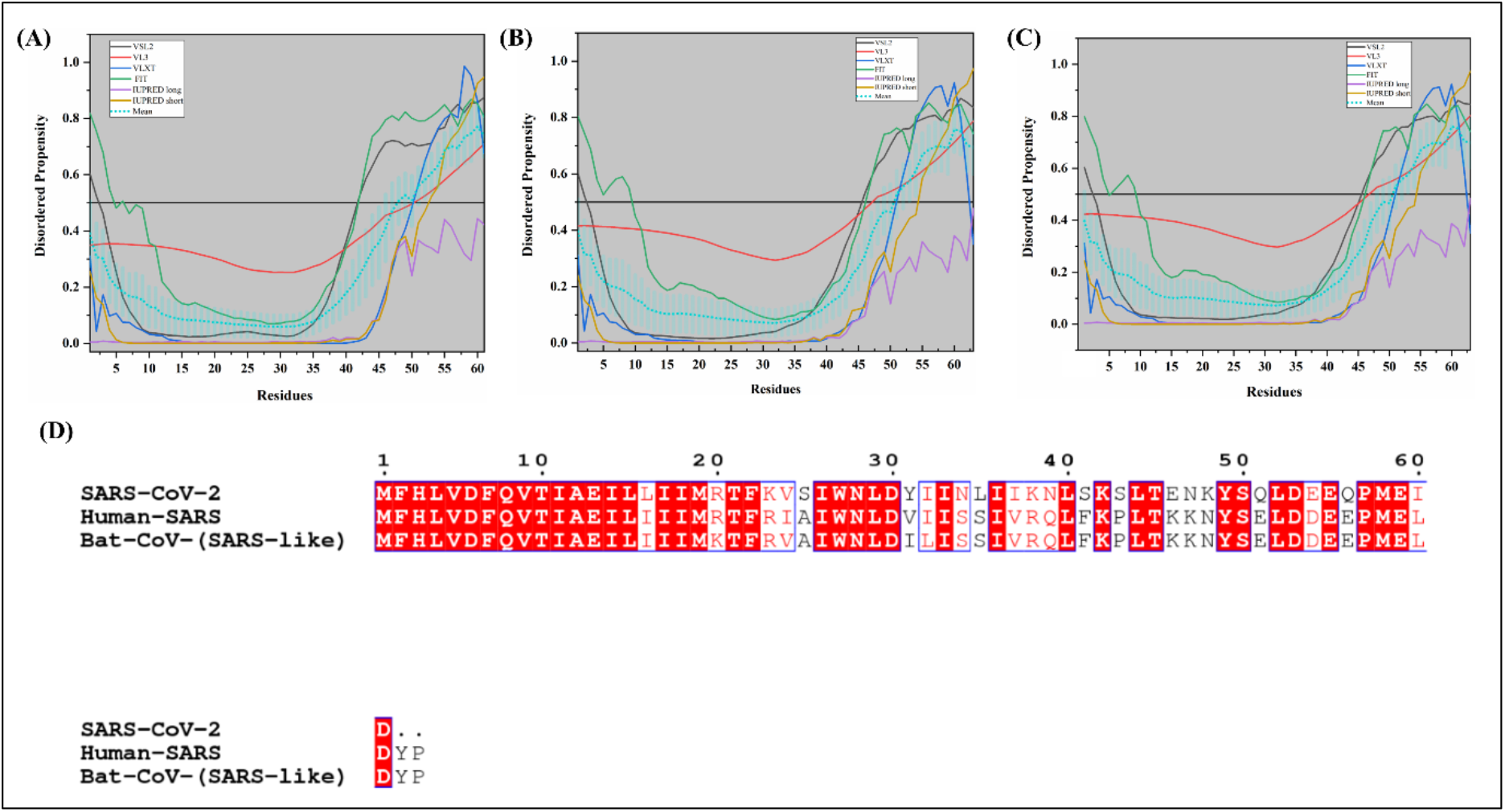
Analysis of intrinsic disorder propensity of ORF6 protein. Graphs represent the intrinsic disorder analysis of ORF6 proteins from SARS-CoV-2 **(A)**, Human SARS **(B)**, and Bat CoV **(C)**. The colour schemes are the same as in the corresponding plots of **Figure 3. (D)** MSA profile of ORF6 of SARS-CoV-2, Human SARS CoV, and Bat CoV. Red shaded sequences represent identical regions of these proteins, whereas red characters show similar residues.

Our analysis of the intrinsic disorder predisposition using six predictors revealed the mean PPID in ORF6 proteins of SARS-CoV-2, Human SARS, and Bat CoV to be 22.95%, 20.63%, and 20.63%, respectively (**Table 1**). Graphs in **Figures 9A, 9B** and **9C** illustrate that ORF6 proteins from all three studied coronaviruses are expected to be moderately disordered proteins with the high disorder content in their C-terminal regions. These disordered regions are important for the biological activities of ORF6. As aforementioned, this hydrophilic region contains lysosomal targeting motif (YSEL) and diacidic motif (DDEE) responsible for binding and recognition during translocation [104]. However, the N-terminal region does not contain a noticeable disorder. The 1-38 residues of the N-terminal region of Human SARS CoV ORF6 was shown to be α-helical and embedded in the membrane, although ORF6 is not a transmembrane protein [106]. A long MoRF region ((residues 26-61 in SARS-CoV-2), (residues 31-63 in Human SARS), and (residues 30-60 in Bat Cov)) is also present at C-terminus of ORF6 proteins which are tabulated in **Table 2**, and **Supplementary Tables 7** and **8**. No predictor other than MoRFchibi_web has located MoRFs in this protein. **Supplementary Table 9, 10, and 11** shows nucleotide-binding residues in ORF6 of all three viruses. It represents very few RNA binding residues by PPRint and few DNA binding residues by DRNApred.

#### ORF7a and ORF7b proteins

Alternatively called U122, ORF7a is a type I transmembrane protein [107,108]. It has been proven to localize in ER, Golgi, and peri-nuclear space. The presence of a KRKTE motif near the C-terminal region is needed for importing this protein from the ER to the Golgi apparatus [107,108]. ORF7a contributes to viral pathogenesis by activating the release of pro-inflammatory cytokines and chemokines, such as IL-8 and RANTES [109,110]. In another study, overexpression of BCL-XL in 293T cells blocked the ORF7a mediated apoptosis [111]. On the other hand, ORF7b is an integral membrane protein that has been shown to localize in the Golgi complex [112,113]. The same reports also confirm the role of ORF7b as an accessory as well as a structural protein in SARS-CoV virion [112,113].

**Figure 10D** represents the 1.8 Å X-ray crystal structure of the 14-96 fragment of the ORF7a from Human SARS CoV (PDB ID: 1XAK) and demonstrates the compact seven-stranded *β* topology of this protein, which is similar to that of the Ig-superfamily members [114]. Importantly, in this crystal structure, residues 82-96 constituted the region with missing electron density, indicating high structural flexibility of this segment. In line with this hypothesis, the NMR solution structure of the 16-99 fragment of the ORF7a from Human SARS CoV (PDB ID: 1YO4) showed that residues 81-99 are highly disordered [115]. At the domain level, the structure of the ORF7a protein includes a signal peptide, a luminal domain, a transmembrane domain, and a short cytoplasmic tail at the C-terminus [95,114].

**Figure 10.**
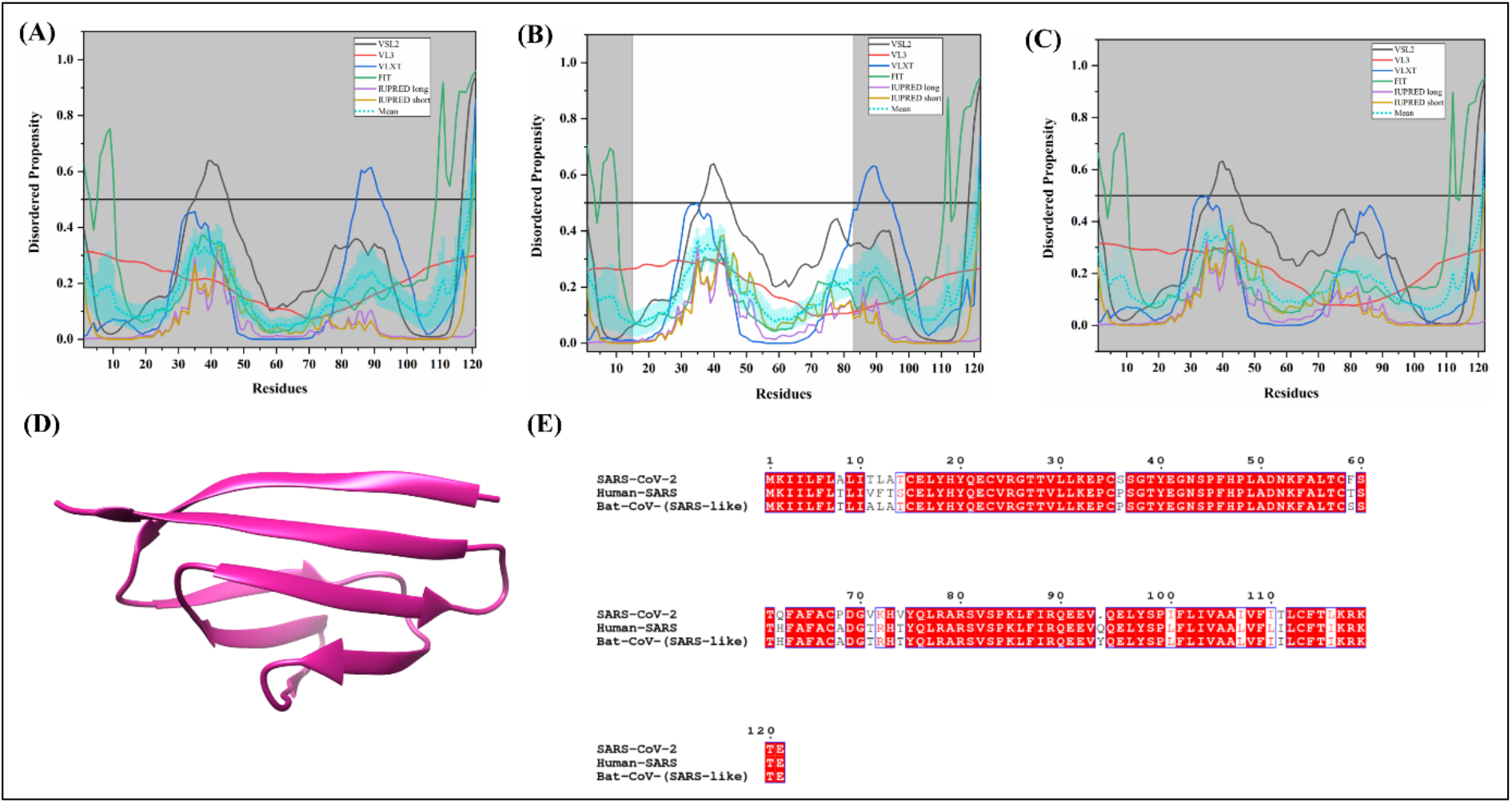
Analysis of intrinsic disorder propensity of ORF7a protein. The graphs represent the results of the intrinsic disorder analysis of ORF7a protein from SARS-CoV-2 **(A)**, Human SARS **(B)**, and Bat CoV **(C)**. The colour schemes are the same as in the corresponding plots of **Figure 3. (D)** A 1.8 Å resolution X-ray structure of ORF7a from Human SARS CoV (PDB ID: 1XAK). It consists of only A chain of 14-96 residues (violet colour). **(E)** MSA profile of ORF7a from SARS-CoV-2, Human SARS CoV, and Bat CoV. Red shaded sequences represent identical regions of these proteins, whereas red characters show similar residues.

We found that 121-residue-long ORF7a protein of SARS-CoV-2 shares 89.26% and 85.95% sequence identity with ORF7a proteins of Bat CoV and Human SARS CoV, respectively (**Figure 10E**). On the other hand, the ORF7b of SARS-CoV-2 is found to be closer to ORF7b of Human SARS than to ORF7b of Bat CoV, showing sequence identities of 81.40% and 79.07%, respectively (see **Figure 11D**).

**Figure 11.**
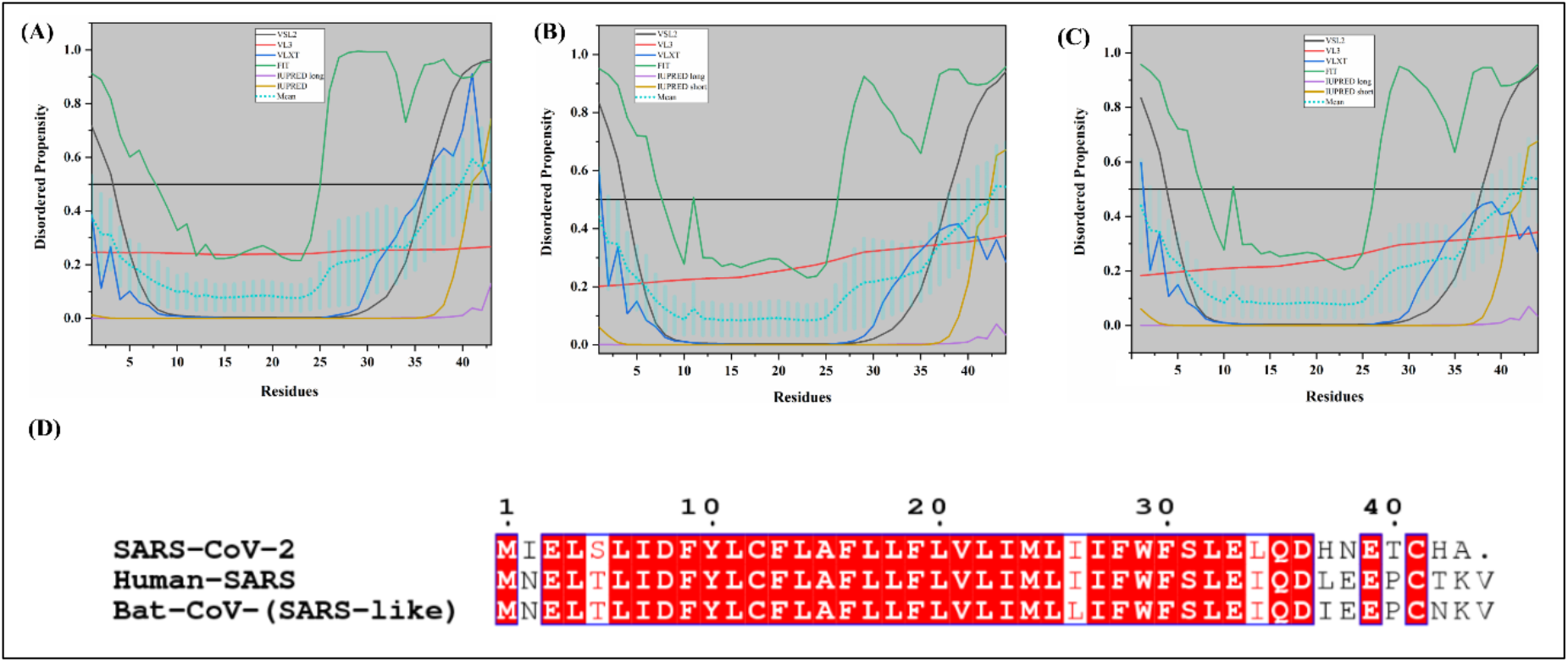
Analysis of intrinsic disorder propensity of ORF7b protein. The graphs represent the results of the intrinsic disorder analysis of the ORF7b protein from SARS-CoV-2 **(A)**, Human SARS CoV **(B)**, and Bat CoV **(C)**. The colour schemes are the same as in the corresponding plots of **Figure 3. (D)** MSA profile of ORF7b from SARS CoV-2, Human SARS CoV, and Bat CoV. Red shaded sequences represent identical regions of these proteins, whereas red characters show similar residues.

As can be observed from **Table 1**, our disorder predisposition analyses resulted in the overall PPID for ORF7a proteins of 1.65% for SARS-CoV-2, 0.82% for Bat CoV and 0.82% for Human SARS CoV. Mean PPIDs estimated for ORF7b proteins are 9.30% for SARS-CoV-2, 4.55% for Bat CoV and 4.55% Human SARS CoV. **Figures 10A, 10B**, and **10C** represent the residues predisposed for disorder in ORF7a proteins of SARS-CoV-2, Human SARS CoV, and Bat CoV, respectively. **Table 2** shows that ORF7a protein is expected to have several MoRFs indicating the potential involvement of this protein in disorder-dependent protein-protein interactions. At the N-terminus, we observed one MoRF region (residues 1-10) with the help of DISOPRED3 in all three viruses. In addition to protein binding regions, ORF7a also contains several RNA and DNA binding residues. Analysis also represents, ORF7b proteins from all three viruses have low disorder content, likewise, they are not predicted to contain any MoRF by any of the predictors used in this study (**Table 2, Supplementary Table 7** and **8**). Although the ORF7b does not contain protein-binding regions, it was found to contain nucleotide (RNS and DNA) binding regions in the protein **Figures 11A, 11B**, and **11C** depict the residues predisposed for disorder in ORF7b proteins of SARS-CoV-2, Human SARS CoV, and Bat CoV, respectively. According to our analysis, both proteins in all three studied coronaviruses have a mostly ordered structure.

#### Proteins ORF8a and ORF8b

In animals and isolates from early human infections, the *ORF8* gene codes for a single ORF8 protein. However, in late infections, more specifically, at middle and late stages, a 29 nucleotide deletion in the *ORF8* gene led to the formation of two distinct proteins, ORF8a and ORF8b containing 39 and 84 residues respectively [116,117].

Both proteins have conformations different from that of the longer ORF8 protein. It has been reported that overexpression of ORF8b resulted in the downregulation of E protein while the proteins ORF8a and ORF8/ORF8ab have no effect on the expression of protein E. Also, ORF8/ORF8ab was found to interact very strongly with proteins S, ORF3a, and ORF7a. ORF8a interacts with S and E proteins, whereas ORF8b protein interacts with E, M, ORF3a and ORF7a proteins [118]. The disorder-based protein binding regions of this protein identified in this study may have important role in interaction with other proteins.

ORF8 protein found in early SARS-CoV-2 isolates having 121 residues and according to our analysis, it shares a 90.05% sequence identity with ORF8 protein of Bat CoV (**Figure 12C**). Furthermore, **Figures 12A** and **12B** show that there is no intrinsic disorder in both ORF8 proteins from SARS-CoV-2 and Bat CoV. Therefore, these two proteins predicted to be completely structured having a mean PPID of 0.00%. In ORF8a and ORF8b proteins of the Human SARS, the predicted disorder is estimated to be 2.56% and 2.38%, respectively **(Table 1)**. Graphs in **Figures 13A** and **13B** illustrate the presence of some disorder near the N- and C-terminals of ORF8a and ORF8b proteins. **Table 2** shows the identified MoRF regions in ORF8 of SARS-CoV-2. It shows three MoRF regions (residues 1-5, 26-52, and 69-91) by MoRFchibi_web and one MoRF region (residues 1-10) by DISOPRED3. In Human SARS, the N-terminus of both ORF8a (residues 1-39) and ORF8b (residues 1-83) was found to be MoRF by MoRFchibi_web server (**Supplementary Table 7**). Further, four protein-binding regions (residues 26-53, 70-91, 98-104, and 113-130) were identified by MoRFchibi_web server in Bat CoV (**Supplementary Table 8**). Apart from protein-binding, ORF8 of SARS-CoV-2, ORF8a and ORF8b of Human SARS, and ORF8 of Bat CoV also comprise several nucleotide-binding residues (see **Supplementary Table 9, 10**, and **11**).

**Figure 12.**
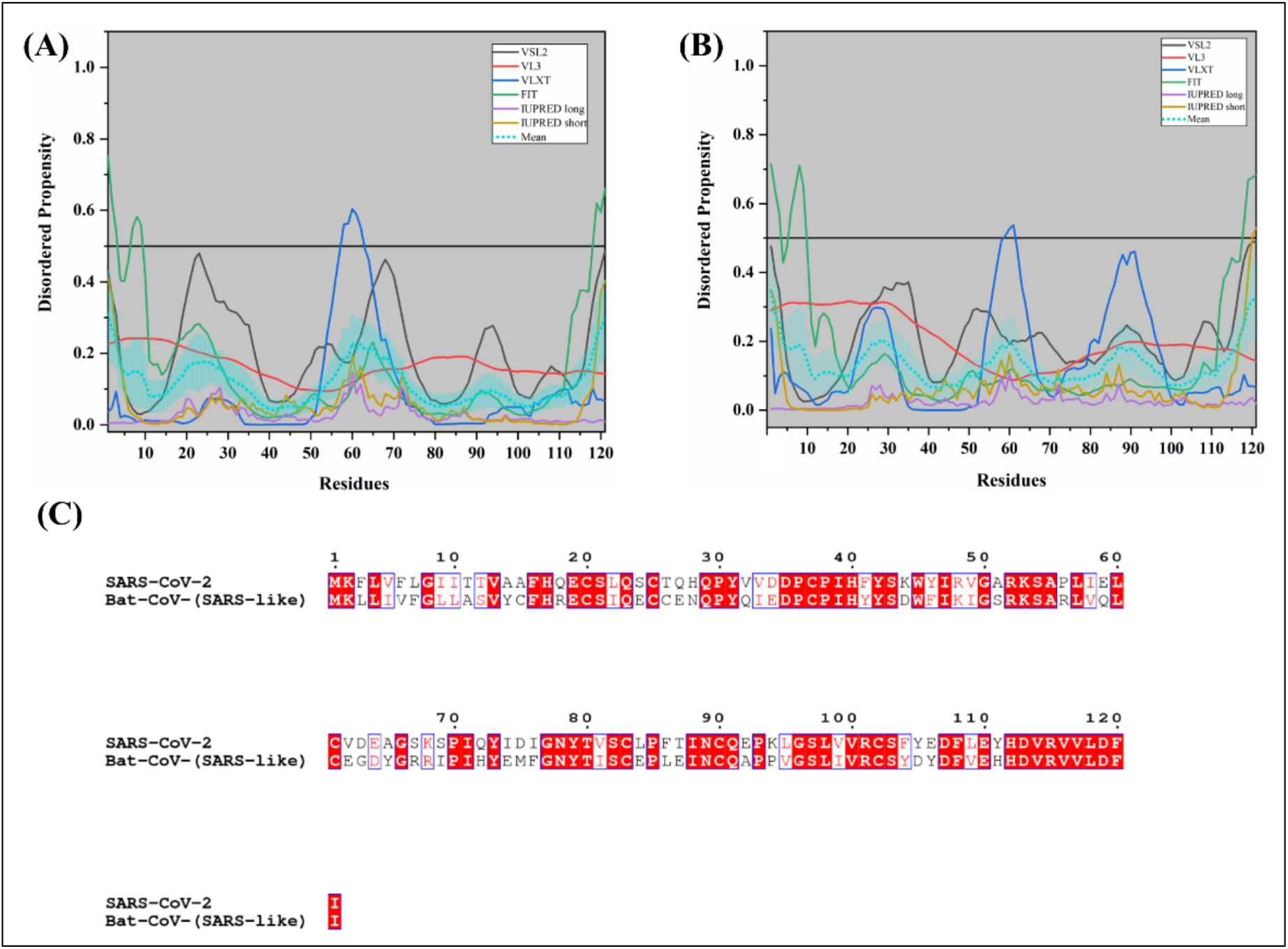
Analysis of intrinsic disorder propensity of ORF8 protein. The graphs represent the results of the intrinsic disorder analysis of the ORF8 protein from SARS-CoV-2 **(A)** and Bat CoV **(B)**. The colour schemes are the same as in the corresponding plots of **Figure 3. (C)** MSA profile of ORF8 protein from SARS-CoV-2 and Bat CoV. Red shaded sequences represent identical regions of these proteins, whereas red characters show similar residues.

**Figure 13.**
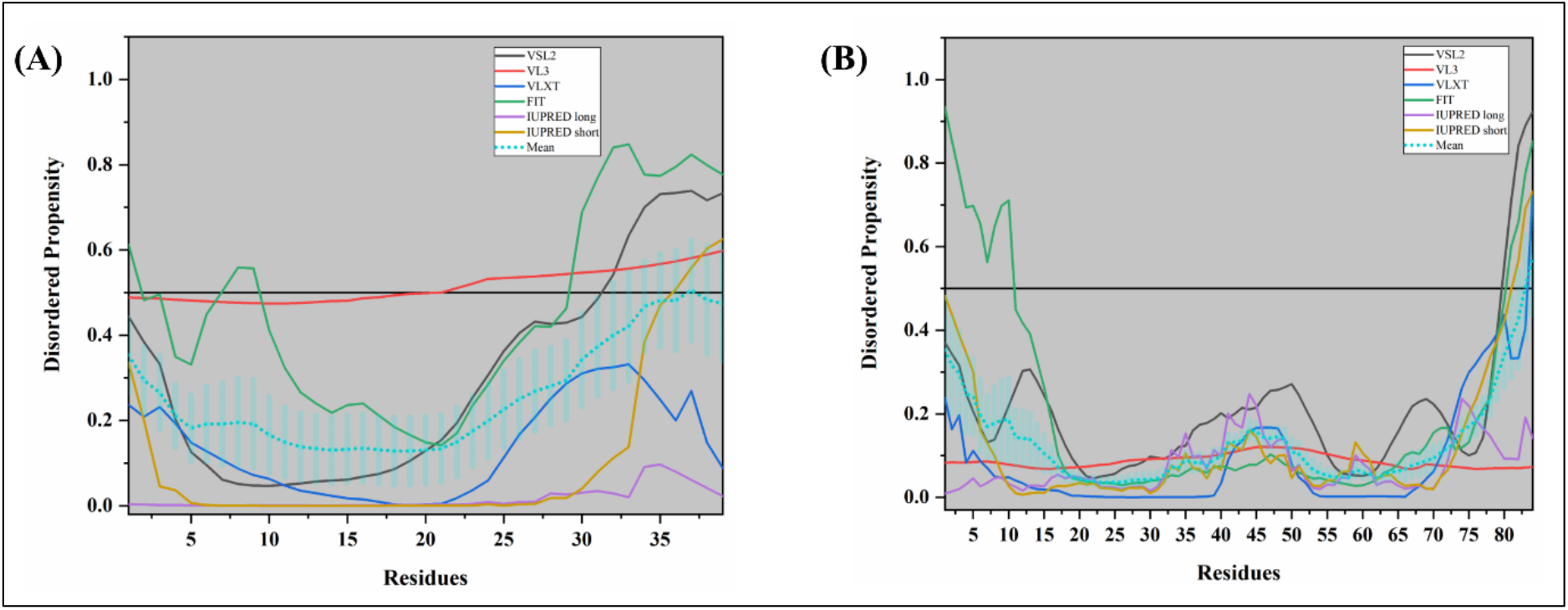
Analysis of intrinsic disorder propensity of ORF8a and ORF8b proteins from Human SARS CoV. **(A)** The graph represents the disorder profile generated for ORF8a protein from Human SARS CoV. **(B)** The graph shows the disorder profile generated for ORF8b protein from Human SARS CoV. The colour schemes are the same as in the corresponding plots of **Figure 3**.

#### ORF9b protein

This protein is expressed from an alternative ORF within the *N* gene through a leaky ribosome binding process [119]. Inside the host cells, ORF9b enters the nucleus, which is a cell cycle-independent process and represents a passive entry. This protein was shown to interact with a nuclear export protein receptor Exportin 1 (Crm1), using which it translocate out of the nucleus [120]. Our MoRFs analysis shows the presence of disorder-based protein binding regions in ORF9b protein which may have role in its interaction with Crm1 and further translocation outside the nucleus. A 2.8 Å resolution crystal structure of ORF9b protein from Human SARS CoV (PDB ID: 2CME) shows the presence of a dimeric tent-like *β*-structure along with the central hydrophobic amino acids (**Figure 14D)**. The published structure has the highly polarized distribution of charges, with positively charged residues on the one side of the tent and negatively charged on the other [121].

**Figure 14.**
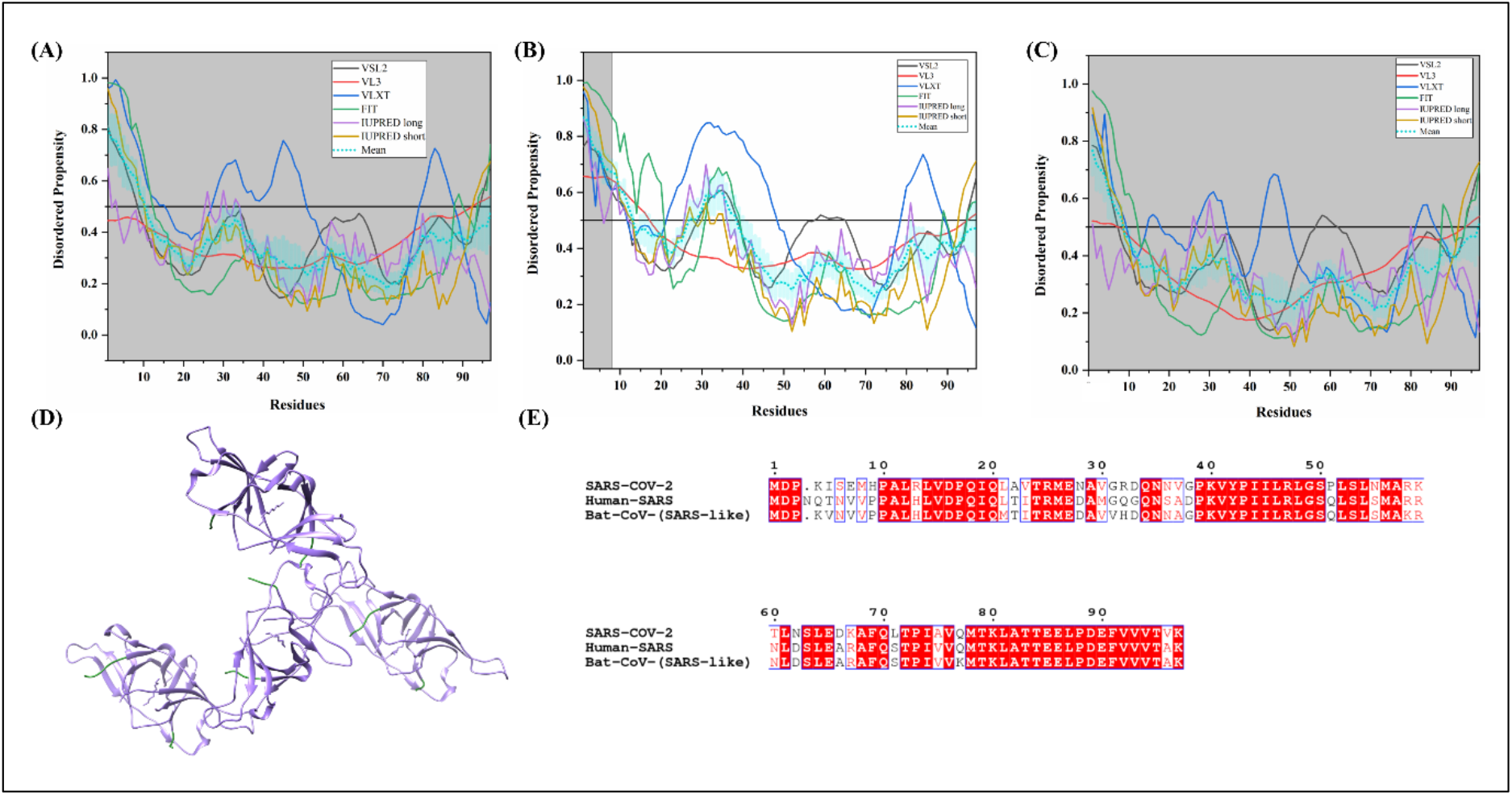
Analysis of intrinsic disorder propensity of ORF9b protein. The graphs represent results of the intrinsic disorder analysis of the ORF9b of SARS-CoV-2 **(A)** and Human SARS CoV **(B)**, as well as Bat CoV of ORF9 protein **(C)**. The colour schemes are the same as in the corresponding plots of **Figure 3. (D)** A 2.8 Å resolution X-ray crystal structure of ORF9b protein from Human SARS CoV (PDB ID: 2CME). This asymmetric unit includes four ORF9b homodimers and consists of A, B, E, and G chains of 9-98 residues (purple colour) and C, D, F, and H chains of 10-98 residues (purple colour). **(E)** MSA of ORF9b of Human SARS CoV and ORF9 of Bat CoV. Red shaded sequences represent identical regions of these proteins, whereas red characters show similar residues.

Based on the sequence availability of accession ID NC_045512.2, the translated protein sequence of ORF9b is not reported for the SARS-CoV-2 as of yet. However, based on the report by Wu and colleagues [45], the sequences of the SARS-CoV-2 are already annotated. Therefore, we took the corresponding amino acid sequences from that study and conducted the intrinsic disorder analysis. According to the MSA, results shown in **Figure 14E**, ORF9b protein from SARS-CoV-2 shares 73.2% identity with Human SARS and 74.23% identity with Bat CoV.

Our IDP analysis (**Table 1**) shows that ORF9b from Human SARS is a moderately unstructured protein with a mean PPID estimated to 26.53%. As depicted in **Figure 14A, 14B**, and **14C**, disorder mainly lies near the N-terminal end 1-10 residues and 28-40 residues near the central region with a well-ordered inner core of Human SARS ORF9b protein. The x-ray crystal structure of ORF9b has a missing electron density of the first 8 residues and 26-37 residues near the central region. This indicates that the corresponding regions are disordered, which are difficult to crystallize due to their highly dynamic structural organization. SARS-CoV-2 ORF9b protein with a mean PPID of 10.31% has an N-terminal (1-10 residues) predicted disordered segment. ORF9b of Bat CoV is shown to have an intrinsic disorder content of 9.28%, comparatively lower than that of the Human SARS ORF9b protein. MoRFs lies in the N-terminal region of ORF9b proteins (**Table 2, Supplementary Table 7** and **8**). In the absence of other viral proteins, its first 41 residues have been demonstrated to induce membranous structures similar to DMVs [106]. The available crystal structure also has the missing electron density in the N-terminal region suggests that these flexible amino acids are likely to interact with host lipids. The first 3-29 residues of SARS-CoV2 are identified as disorder-based protein binding region that may have role in its interaction with host lipids and formation of DMVs. **Supplementary Tables 9, 10**, and **11** represents nucleotide-binding residues for ORF9b of SARS-CoV-2, Human SARS, and Bat CoV.

#### ORF10 protein

The newly emerged SARS-CoV-2 has an ORF10 protein of 38 amino acids. ORF10 of SARS-CoV-2 has a 100% sequence similarity with ORF10 of Bat CoV strain Bat-SL-CoVZC45 [21]. However, we did not conduct the disorder analysis for ORF10 from the Bat-SL-CoVZC45 strain, since all our studies reported here are related to a different strain of Bat CoV (reviewed strain HKU3-1). Therefore, we report here only the results of disorder analysis for the ORF10 protein from SARS-CoV-2, according to which this protein has a mean PPID of 0.00% (see also **Figure 15** for disorder profile of ORF10). This protein contains a MoRF from 3-7 residues at its N-terminus as predicted by MoRFchibi_web. Further, we found its tendency to nucleotides and found the presence of few RNA binding sites, however, it does not contain DNA binding residues.

**Figure 15.**
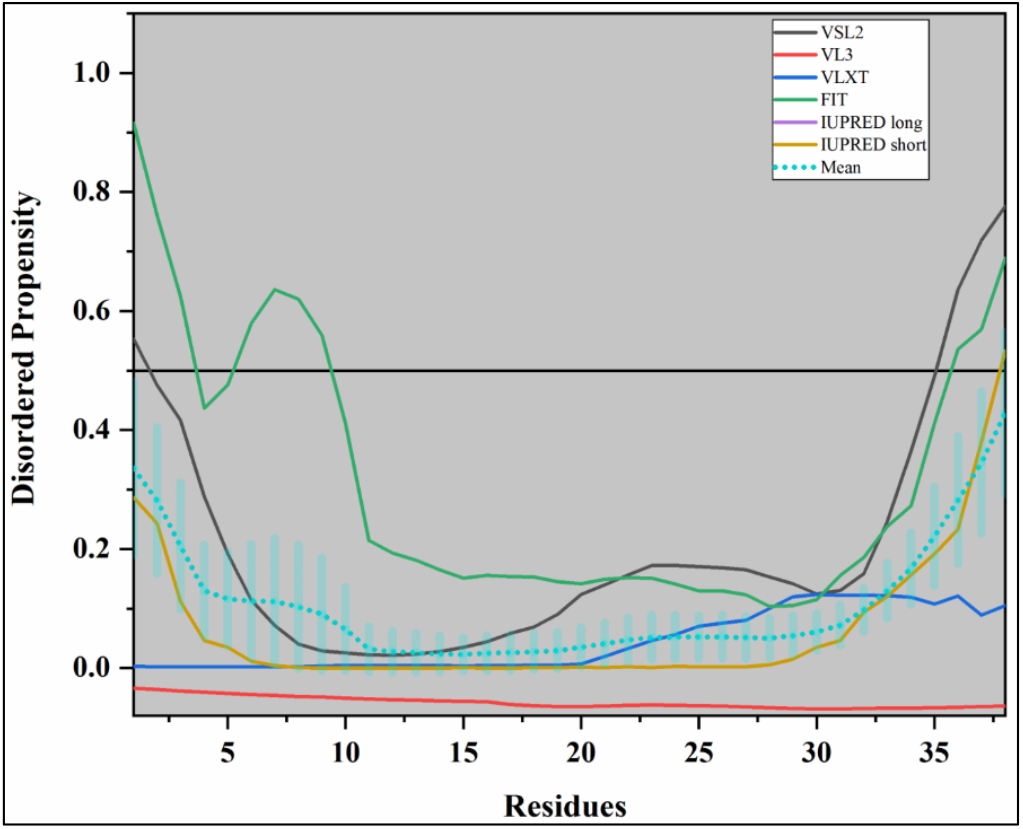
Analysis of intrinsic disorder propensity of ORF10 protein of SARS-CoV-2. The colour schemes are the same as in the corresponding plots of **Figure 3**.

#### Protein ORF14

This is a 70-amino-acid-long uncharacterized protein of unknown function, which is present in Human SARS and Bat CoV. In SARS-CoV-2, ORF14 is a 73-amino-acid-long protein. According to the MSA, ORF14 of SARS-CoV-2 has 77.1% identity with Human-SARS and 72.9% identity with Bat CoV as represented in **Figure 16D**. We have performed the intrinsic disorder analysis to see the peculiarities of the distribution of disorder predisposition in this protein. **Figures 16A, 16B**, and **16C** show the resulting disorder profiles of ORF14 of SARS-CoV-2, Human SARS CoV, and Bat CoV. Although these proteins have calculated mean PPID values of 0.00%, 2.86%, and 0.00% respectively, **Figure 16** shows that they have flexible N- and C-terminal regions. This protein can use intrinsic disorder or structural flexibility for protein-protein interactions since it possesses MoRFs. It mainly contains MoRFs at N- and C-terminal regions as tabulated in (**Table 2, Supplementary Table 7** and **8**). It was also found to contain several RNA and DNA binding residues (**Supplementary Table 9, 10**, and **11)**. These results indicating its vital role in protein function related to molecular recognition such as protein-protein, protein-RNA, and Protein-DNA interaction.

**Figure 16.**
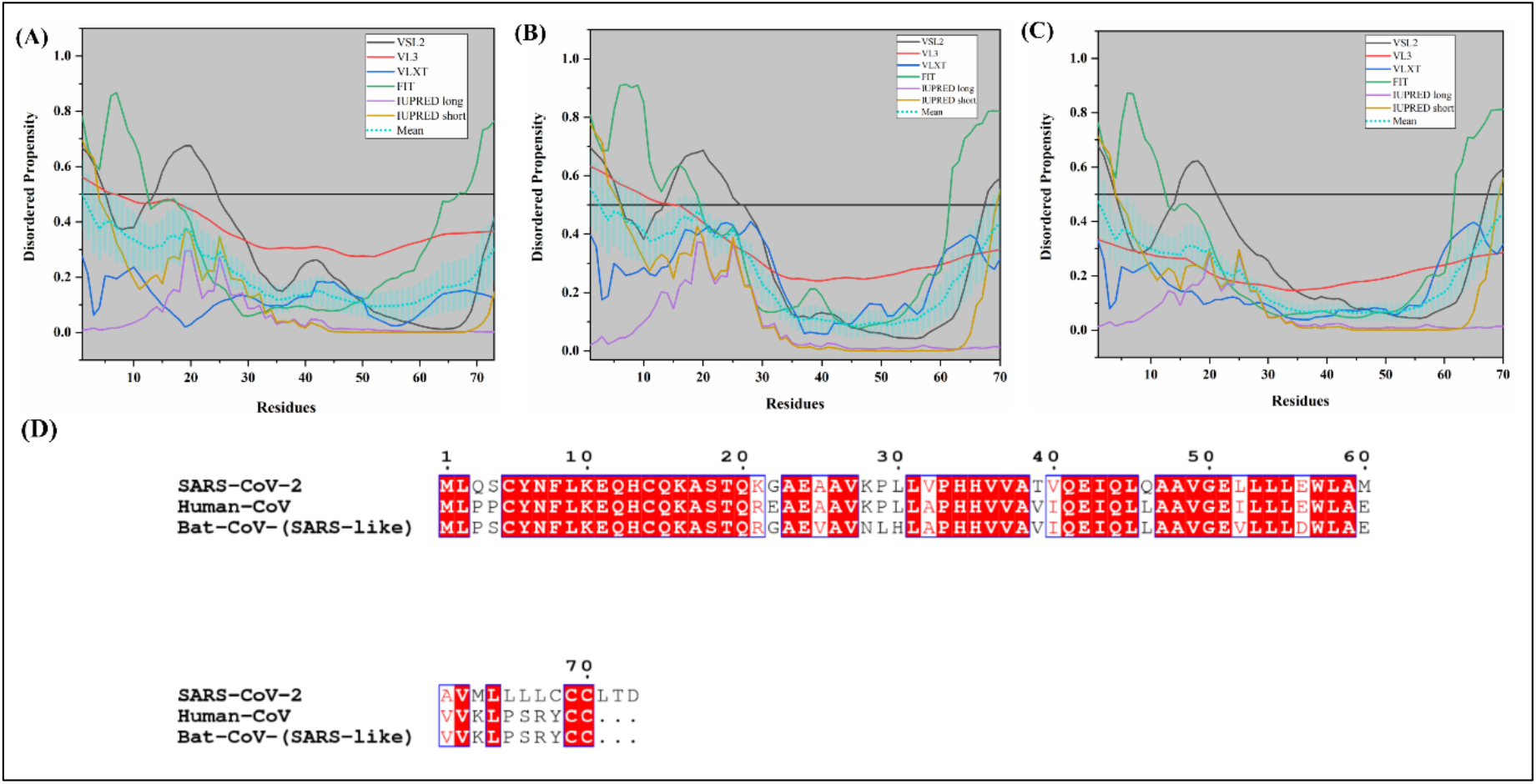
Analysis of intrinsic disorder propensity of ORF 14 protein. The graphs represent the results of the intrinsic disorder analysis of the ORF14 protein from SARS-CoV-2 **(A)**, Human SARS CoV **(B)**, and Bat CoV **(C)**. The colour schemes are the same as in the corresponding plots of **Figure 3. (D)** MSA profile of ORF14 from SARS CoV-2, Human SARS CoV, and Bat CoV. Red shaded sequences represent identical regions of these proteins, whereas red characters show similar residues.

### Intrinsic disorder analysis of non-structural proteins of Coronaviruses

In coronaviruses, due to ribosomal leakage during translation, two-third of the RNA genome is processed into two polyproteins: (i) Replicase polyprotein 1a and (ii) Replicase polyprotein 1ab. Both contain non-structural proteins (Nsp1-10) in addition to different proteins required for viral replication and pathogenesis. Replicase polyprotein 1a contains an additional Nsp11 protein of 13 amino acids, the function of which is not investigated yet. The longer replicase polyprotein 1ab of 7073 amino acids accommodates five other non-structural proteins (Nsp12-16) [122]. These proteins assist in ER membrane-induced vesicle formation, which acts as sites for replication and transcription. In addition to this, non-structural proteins work as proteases, helicases, and mRNA capping and methylation enzymes, crucial for virus survival and replication inside host cells [122,123].

### Global analysis of intrinsic disorder in the replicase polyprotein 1ab

**Table 3** represents the PPID mean scores of 15 non-structural proteins (Nsps) derived from the Replicase polyprotein 1ab in SARS-CoV-2, Human SARS CoV, and Bat CoV. These values were obtained by combining the results from six disorder predictors (see **Supplementary Table S4-S6**). **Figures 17A, 17B**, and **17C** represent the 2D-disorder plots of the Nsps coded by ORF1ab in SARS-CoV-2, Human SARS Cov, and Bat CoV, respectively. Based on the mean PPID scores in **Table 3, Figures 17A, 17B, 17C**, and taking into PPID based classification [46], we conclude that none of the Nsps in SARS-CoV-2, Human SARS CoV, and Bat CoV are highly disordered. The highest disorder was observed for Nsp8 proteins in all three Coronaviruses. Both Nsp1 and Nsp8 are moderately disordered proteins (10% ≤ PPID ≤ 30%). We also observed that Nsp2, Nsp3, Nsp5, Nsp6, Nsp7, Nsp9, Nsp10, Nsp15, and Nsp16 have less than 10% disordered residues and hence, belong to the category of mostly ordered proteins. Other non-structural proteins, namely, Nsp4, Nsp12, Nsp13, and Nsp14 have negligible levels of disorder (PPID < 1%), which tells us that these are highly structured proteins.

**Table 3.** Evaluation of mean predicted percentage disorder in non-structural proteins of novel SARS-CoV-2, Human SARS, and Bat CoV.

| Proteins | SARS-CoV-2 |  | Human SARS<br>(UniProt ID: P0C6X7) |  | Bat CoV (SARS-like)<br>(UniProt ID: P0C6W2) |  |
| --- | --- | --- | --- | --- | --- | --- |
|  | Length of protein<br>(Residues)<br>(NCBI RefSeq<br>accession ID) | Mean<br>PPID | Length of<br>protein<br>(Residues) | Mean<br>PPID | Length of<br>protein<br>(Residues) | Mean<br>PPID |
| **Nsp1** (Host translation inhibitor) | 180<br>(1-180)<br>(YP_009725297.1) | 12.78 | 180<br>(1 – 180) | 14.44 | 179<br>(1-179) | 12.85 |
| **Nsp2** | 638<br>(181-818)<br>(YP_009725298.1) | 5.17 | 638<br>(181 – 818) | 2.04 | 639<br>(180-818) | 2.03 |
| **Nsp3** (papain-like proteases) | 1,945<br>(819-2763)<br>(YP_009725299.1) | 7.40 | 1,922<br>(819 – 2740) | 7.91 | 1,916<br>(819 – 2734) | 7.78 |
| **Nsp4** | 500<br>(2764-3263)<br>(YP_009725300.1) | 0.80 | 500<br>(2741 – 3240) | 0.60 | 500<br>(2735 – 3234) | 0.60 |
| **Nsp5** (3CL-Pro) | 306<br>(3264-3569)<br>(YP_009725301.1) | 1.96 | 306<br>(3241 – 3546) | 1.96 | 306<br>(3235 – 3540) | 1.96 |
| **Nsp6** | 290<br>(3570-3859)<br>(YP_009725302.1) | 1.03 | 290<br>(3547 – 3836) | 1.03 | 290<br>(3541 – 3830) | 1.03 |
| **Nsp7** (multimeric RNA polymerase) | 83<br>(3860-3942)<br>(YP_009725303.1) | 9.64 | 83<br>(3837 – 3919) | 9.64 | 83<br>(3831 – 3913) | 9.64 |
| **Nsp8** (multimeric RNA polymerase) | 198<br>(3943-4140)<br>(YP_009725304.1) | 23.74 | 198<br>(3920 – 4117) | 23.74 | 198<br>(3914 – 4111) | 22.22 |
| **Nsp9** | 113<br>(4141-4253)<br>(YP_009725305.1) | 7.08 | 113<br>(4118 – 4230) | 7.96 | 113<br>(4112 – 4224) | 7.08 |
| **Nsp10** | 139<br>(4254-4392) | 5.04 | 139<br>(4231 – 4369) | 5.04 | 139<br>(4225 – | 5.04 |
|  | (YP_009725306.1) |  |  |  | 4363) |  |
| **Nsp12** (RNA-directed RNA polymerase, RDRP) | 932<br>(4393-5324)<br>(YP_009725307.1) | 0.43 | 932<br>(4370 – 5301) | 0.43 | 932<br>(4364 – 5295) | 0.43 |
| **Nsp13** (Viral helicase) | 601<br>(5325-5925)<br>(YP_009725308.1) | 0.67 | 601<br>(5302 – 5902) | 0.67 | 601<br>(5296 – 5896) | 0.67 |
| **Nsp14** (Guanine-N7 methyltransferase/ExoN-nsp14) | 527<br>(5926-6452)<br>(YP_009725309.1) | 0.38 | 527<br>(5903 – 6429) | 0.38 | 527<br>(5897 – 6423) | 0.57 |
| **Nsp15** (Uridylate-specific endoribonuclease/ NendoU) | 346<br>(6453-6798)<br>(YP_009725310.1) | 1.73 | 346<br>(6430 – 6775) | 2.60 | 346<br>(6424 – 6769) | 2.60 |
| **Nsp16** (2'-O-methyltransferase) | 298<br>(6799-7096)<br>(YP_009725311.1) | 5.37 | 298<br>(6776 – 7073) | 3.02 | 298<br>(6770 – 7067) | 3.02 |

Proteins and their PPIDs are coloured to reflect their disorder status (ordered – blue, moderately disordered – pink, highly disordered (if any) – red)

**Figure 17.**
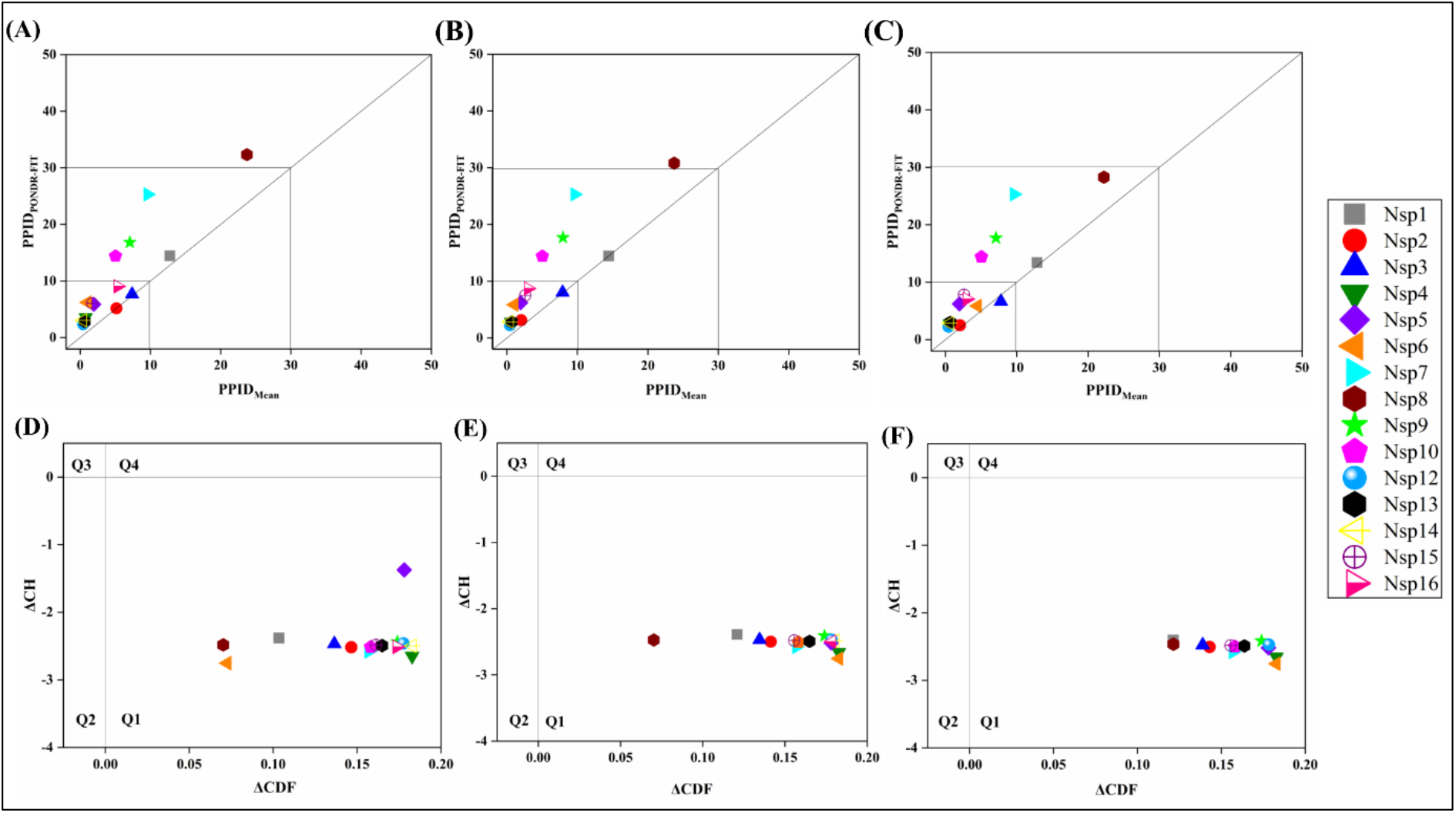
Analysis of the overall intrinsic disorder status of non-structural proteins (Nsps) coded by *ORF1ab* in SARS-CoV-2, Human SARS CoV, and Bat CoV: 2D plot representing PPID_PONDR-FIT_ vs PPID_Mean_ in **(A)** SARS-CoV-2 **(B)** Human SARS and **(C)** Bat CoV (SARS-like). In CH-CDF plot of the proteins of **(D)** SARS-CoV-2 **(E)** Human SARS and **(F)** Bat CoV, the Y coordinate of each protein spot signifies distance of corresponding protein from the boundary in CH plot and the X coordinate value corresponds to the average distance of the CDF curve for the respective protein from the CDF boundary.

The CH-CDF analysis of the Nsps from SARS-CoV-2, Human SARS and Bat CoV have been represented in **Figures 17D, 17E**, and **17F** respectively. It was observed that all the Nsps of the three coronaviruses are located within the quadrant Q1 of the CH-CDF phase space, indicating that all the Nsps are predicted to be mostly ordered.

#### Replicase polyprotein 1ab

The longer replicase polyprotein 1ab is a 7,073 amino acid-long polypeptide, which contains 15 non-structural proteins listed in **Table 3**. Nsp1, Nsp2, and Nsp3 are cleaved using a viral papain-like proteinase (Nsp3/PL-Pro), while the rest of Nsps are cleaved by another viral 3C-like proteinase, Nsp5/3CL-Pro. We mapped the cleavage sites of the replicase 1ab polyprotein from Human SARS CoV to the disorder profile of this polyprotein. **Figure 18** represents the results of this analysis by showing zoomed-in regions surrounding all the cleavage sites with few residues spanning at both terminals. Interestingly, we observed that all the cleavage sites are largely disordered, suggesting that intrinsic disorder may have a crucial role in the maturation of individual non-structural proteins. As the Nsps of Human SARS CoV are evolutionary close to the Nsps of SARS-CoV-2, we hypothesize that the cleavage sites in the SARS-CoV-2 replicase 1ab polyprotein are also intrinsically disordered or flexible. To shed more light on other implications of IDPRs, the structural and functional properties of Nsps and their predicted IDPRs are thoroughly described below.

**Figure 18.**
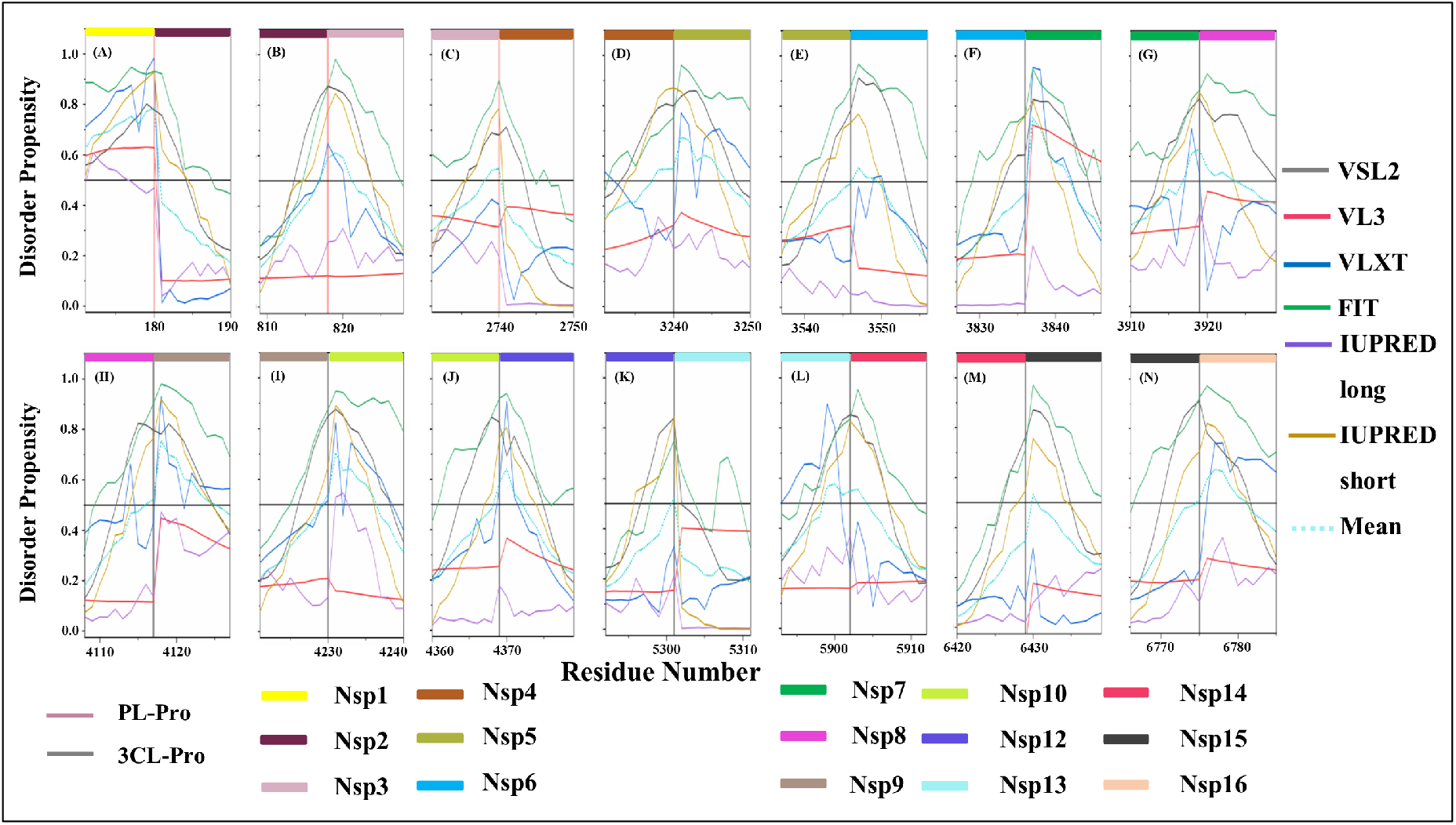
Intrinsic disorder at the cleavage sites of the replicase 1ab polyprotein of Human SARS CoV. Plots **A** to **N** denotes the cleavage sites (magenta coloured bar for PL-Pro protease and grey coloured bar for 3CL-Pro protease) in relation to disordered regions present between the individual proteins (Nsp1-16) of replicase 1ab polyprotein of Human SARS. All proteins are represented by different coloured horizontal bars.

#### Non-structural protein 1 (Nsp1)

This protein acts as a host translation inhibitor as it binds to the 40S subunit of the ribosome and blocks the translation of cap-dependent mRNAs as well as mRNAs that uses the internal ribosome entry site (IRES) [124]. **Figure 19D** shows the NMR solution structure (PDB ID: 2GDT) of Human SARS nsp1 protein (13-128 residues), whereas residues 117-180 were not included in this structural analysis [125].

**Figure 19.**
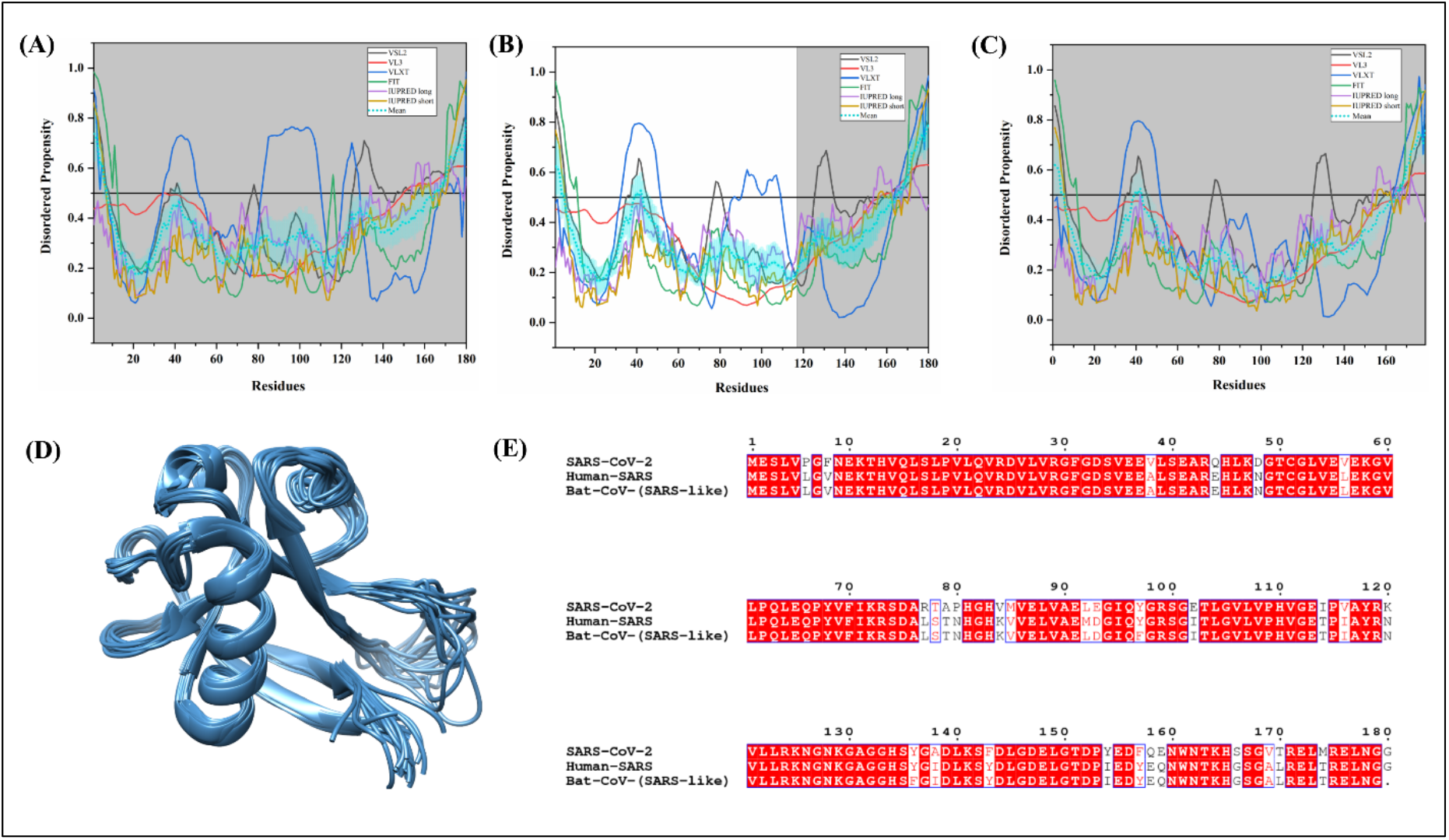
Analysis of intrinsic disorder propensity and structural characterization of Nsp1. Intrinsic disorder profiles generated for Nsp1 from SARS-CoV-2 **(A)**, Human SARS CoV **(B)**, and Bat CoV **(C)**. The colour schemes are the same as in the corresponding plots of **Figure 3. (D)** NMR solution structure of a 13-128 fragment of Human SARS Nsp1 (PDB ID: 2GDT). **(E)** MSA profile of Nsp1 from SARS CoV-2, Human SARS CoV, and Bat CoV. Red shaded sequences represent identical regions of these proteins, whereas red characters show similar residues.

SARS-CoV-2 Nsp1 shares 84.44% and 83.80% sequence identity with Nsp1s of Human SARS CoV and Bat CoV, respectively. Its N-terminal region is found to be more conserved than the rest of the protein sequence (**Figure 19E)**. Mean PPIDs of Nsp1s from SARS-CoV-2, Human SARS CoV, and Bat CoV are 12.78%, 14.44%, and 12.85%, respectively. **Figure 19A, 19B**, and **19C** represent the graphs of predicted per-residue intrinsic disorder propensity of these Nsp1s. According to the analysis, the following regions are predicted to be disordered: SARS-CoV-2 (residues 1-7 and 165-180), Human SARS CoV (residues 1-5 and 165-180), and Bat CoV (residues 1-5 and 165-179). NMR solution structure of Nsp1 from Human SARS revealed the presence of two unstructured segments near the N-terminal (1-12 residues) and C-terminal (129-179 residues) regions [125]. The disordered region (128–180 residues) at C-terminus is important for Nsp1 expression [126]. Based on sequence homology with Human SARS CoV Nsp1, the predicted disordered C-terminal region of SARS-CoV-2 Nsp1 may play a critical role in its expression. Alanine mutants at K164 and H165 in the C-terminal region of Nsp1 protein is reported to abolish its binding with the 40S subunit of the host ribosome [127]. In conjunction with this data, several MoRFs are present in the unstructured segments of Nsp1 proteins. These regions are tabulated in **Table 2**, and **Supplementary Tables 7** and **8**.

#### Non-structural protein 2 (Nsp2)

This protein functions by disrupting the host survival pathway via interaction with the host proteins Prohibitin-1 and Prohibitin-2 [128]. Reverse genetic deletion in the coding sequence of Nsp2 of the SARS virus attenuated little viral growth and replication and allowed the recovery of mutant virulent viruses. This indicates the dispensable nature of the Nsp2 protein for SARS viruses [129].

The sequence identity of the Nsp2 protein from SARS-CoV-2 with Nsp2s of Human SARS CoV and Bat CoV amounts to 68.34% and 68.97%, respectively (**Supplementary Figure S2A**). We have estimated the mean PPIDs of Nsp2s of SARS-CoV-2, Human SARS CoV, and Bat CoV to be 5.17%, 2.04%, and 2.03% respectively (see **Table 3**). The per-residues predisposition for the intrinsic disorder of Nsp2s from SARS-CoV-2, Human SARS CoV, and Bat CoV are depicted in **Figures 20A, 20B**, and **20C**. According to this analysis, the following regions in Nsp2 proteins are predicted to be disordered, residues 570-595 (SARS-CoV-2), residues 110-115 (Human SARS), and residues 112-116 (Bat CoV). As listed in **Table 2**, and **Supplementary Tables 7** and **8**, Human SARS CoV does not contain MoRF while SARS-CoV-2 and Bat CoV have an N-terminally located MoRF region predicted by MoRFchibi_web.

**Figure 20.**
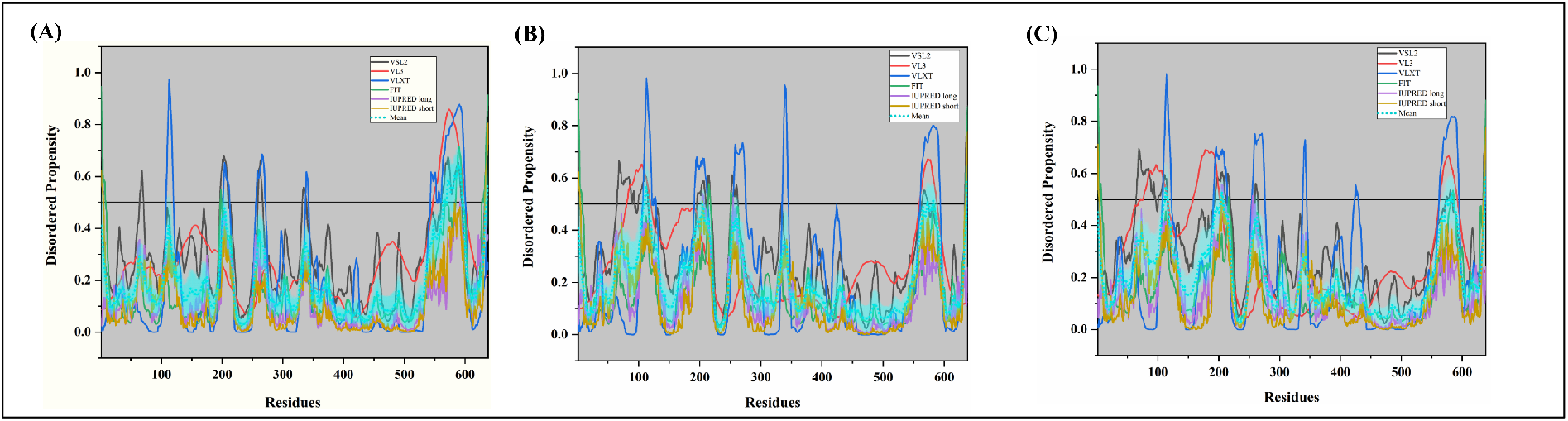
Analysis of the intrinsic disorder propensity of Nsp2. Intrinsic disorder profiles generated for Nsp2 from SARS-CoV-2 **(A)**, Human SARS CoV **(B)**, and Bat CoV **(C)**. The colour schemes are the same as in the corresponding plots of **Figure 3**.

#### Non-structural protein 3 (Nsp3)

Nsp3 is an almost 2,000-residue-long viral papain-like protease (PLP) that affects the phosphorylation and activation of IRF3 and therefore antagonizes the IFN pathway [130]. It was also demonstrated that Nsp3 works by stabilizing NF-κ*β* inhibitor further blocking the NF-κ*β* pathway [130]. **Figure 21D** represents the 1.85 Å resolution X-ray crystal structure of the catalytic core of Nsp3 protein from Human SARS CoV (PDB ID: 2FE8), which was obtained by Andrew and colleagues [131]. This structure consists of the residues 723-1036 of Nsp3. The structure revealed folds similar to a deubiquitinating enzyme in-vitro deubiquitinating activity of which was found to be efficiently high [131].

**Figure 21.**
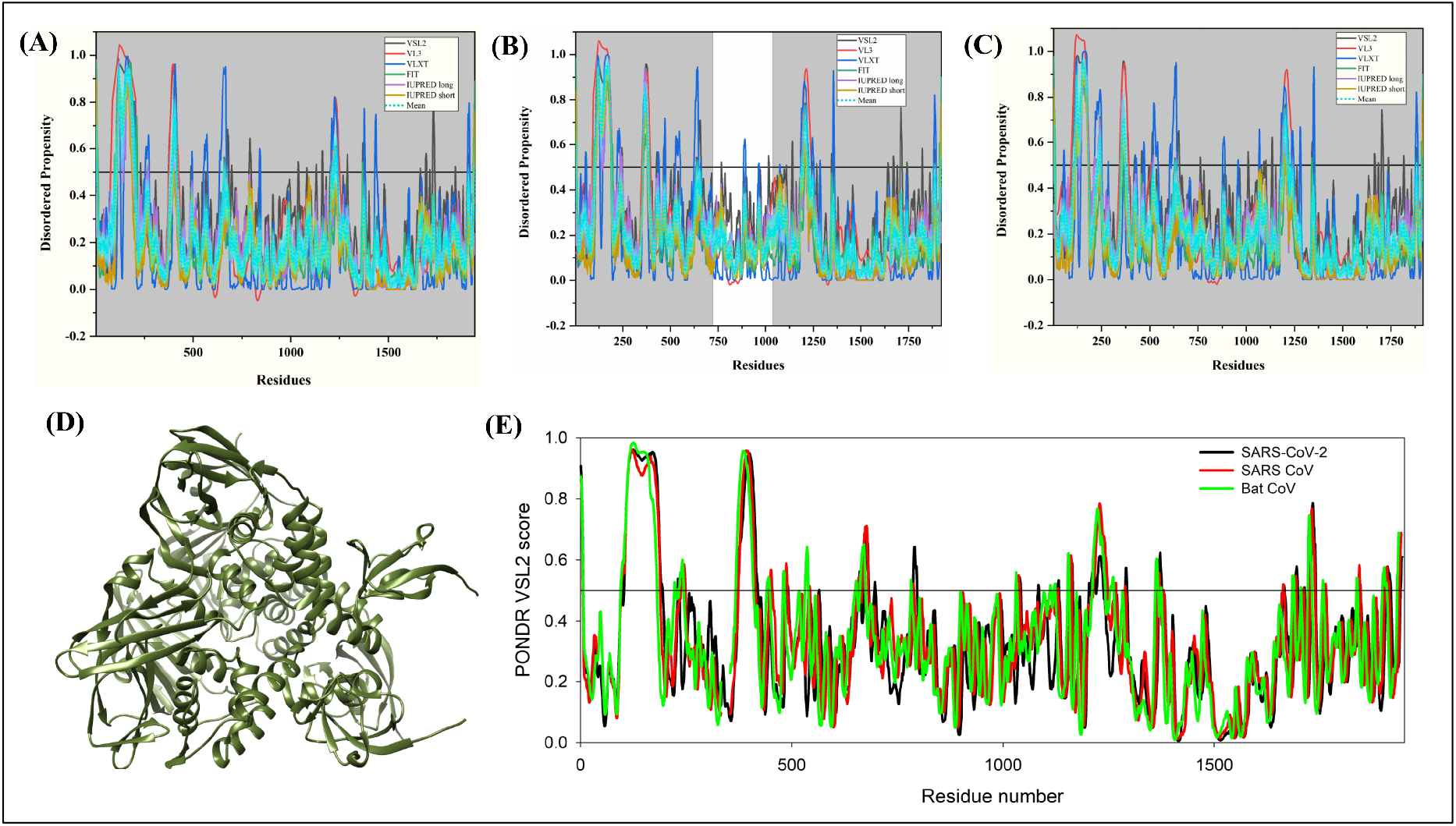
Analysis of intrinsic disorder propensity of Nsp3. Intrinsic disorder profiles generated for Nsp3 from SARS-CoV-2 **(A)**, Human SARS CoV **(B)**, and Bat CoV **(C)**. The colour schemes are the same as in the corresponding plots of **Figure 3. (D)** A 1.85 Å resolution X-ray crystal structure of the region containing residues 723-1036 of Nsp3 of Human SARS CoV (PDB ID: 2FE8). **(E)** Aligned disorder profiles generated for Nsp3 from SARS-CoV-2 (black line), Human SARS CoV (red line), and Bat CoV (green line) based on the outputs of the PONDR^®^ VSL2.

Nsp3 protein of SARS-CoV-2 contains several substituted residues throughout the protein. It is equally close with both Nsp3 proteins of Human SARS and Bat CoV sharing respective 76.69% and 76.31% identity **(Supplementary Figure S2B)**. According to our results, the mean PPIDs of Nsp3 proteins of SARS-CoV-2, Human SARS, and Bat CoV are 7.40%, 7.91%, and 7.78% respectively (**Table 3**). Graphs in **Figures 21A, 21B**, and **21C** portray the tendency of Nsp3 proteins of SARS-CoV-2, Human SARS, and Bat CoV for the intrinsic disorder. Nsp3 proteins of all three studied SARS viruses were found to be highly structured and characterized by rather similar disorder profiles. This is further supported by **Figure 21E**, where PONDR^®^ VSL2-generated disorder profiles of these three proteins are overlapped to show almost complete coincidence of their major disorder-related features. According to the mean disorder analysis (see **Figures 21A, 21B**, and **21C**), Nsp3 proteins are predicted to have the following IDPRs, SARS-CoV-2 (1-5, 105-199, 1221-1238), Human SARS (102-189, 355-384, 1195-1223) and Bat CoV (107-182, 352-376, 1191-1217). The First 112 residues in Nsp3 represent a ubiquitin-like globular fold while 113-183 residues form the flexible acidic domain rich in glutamic acid. It is thought to bind and ubiquitinate viral E protein using the N-terminal acidic domain [132,133]. This unstructured segment has many MoRFs predicted by ANCHOR and MoRFPred servers which may facilitate the protein-protein interaction (**Table 2**). Interestingly, Nsp3 of all three viruses was found with highest number of RNA-binding residues (**Supplementary Tables 9, 10**, and **11**).

#### Non-structural protein 4 (Nsp4)

Nsp4 has been reported to induce the formation of the double-membrane vesicles (DMVs) with the co-expression of full-length Nsp3 and Nsp6 proteins for optimal replication inside host cells [134–136]. It localizes itself in ER-membrane, when expressed alone but is demonstrated to be present in replication units in infected cells. It was observed that Nsp4 protein contains a tetraspanning transmembrane region having its N- and C-terminals in the cytosol [137]. No crystal or NMR solution structure is reported for this protein as of yet.

Nsp4 protein of SARS-CoV-2 has multiple substitutions near the N-terminal region and has a quite conserved C-terminus **(Supplementary Figure S2C)**. It is found to be closer to Nsp4 of Bat CoV (81.40% identity) than to Human SARS Nsp4 (80%). Mean PPIDs of Nsp4s from SARS-CoV-2, Human SARS, and Bat CoV are estimated to be 0.80%, 0.60%, and 0.60% respectively. The low level of intrinsic disorder is further illustrated by **Figures 22A, 22B**, and **22C**. With PPIDs around zero, Nsp4 were classified as highly structured proteins, which, however, contain some flexible regions. Likewise, **Table 2** shows the presence of only N- and C-terminal MoRFs which possibly assist in cleavage of Nsp4 protein from long polyproteins 1a and 1ab.

**Figure 22.**
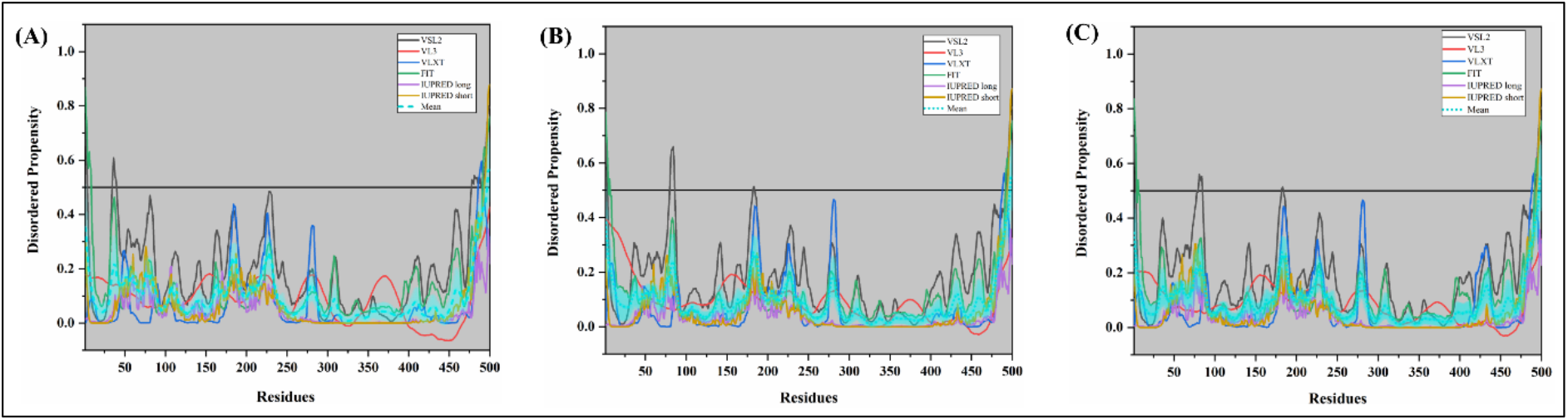
Analysis of intrinsic disorder propensity of Nsp4. Intrinsic disorder profiles generated for Nsp4 from SARS-CoV-2 **(A)**, Human SARS CoV **(B)**, and Bat CoV **(C)**. The colour schemes are the same as in the corresponding plots of **Figure 3**.

#### Non-structural protein 5 (Nsp5)

Also referred to as 3CL-pro, Nsp5 works as a protease that cleaves the replicase polyproteins (1a and 1ab) at 11 major sites [138,139]. X-ray crystal structure with 1.5 Å resolution (PDB ID: 5C5O) obtained for Human SARS CoV Nsp5 is shown in **Figure 23D**. Here, 3CL-protease is bound to a phenyl-beta-alanyl (S, R)-N-declin type inhibitor. Another crystal structure resolved to 1.96 Å revealed a chymotrypsin-like fold and a conserved substrate-binding site connected to a novel α-helical fold [140]. Recently, the X-ray crystal structure (resolution 2.16 Å) was solved for the SARS-CoV-2 Nsp5 in complex with an inhibitor N3 (PDB ID: 6LU7) (**Figure 23E)**.

**Figure 23.**
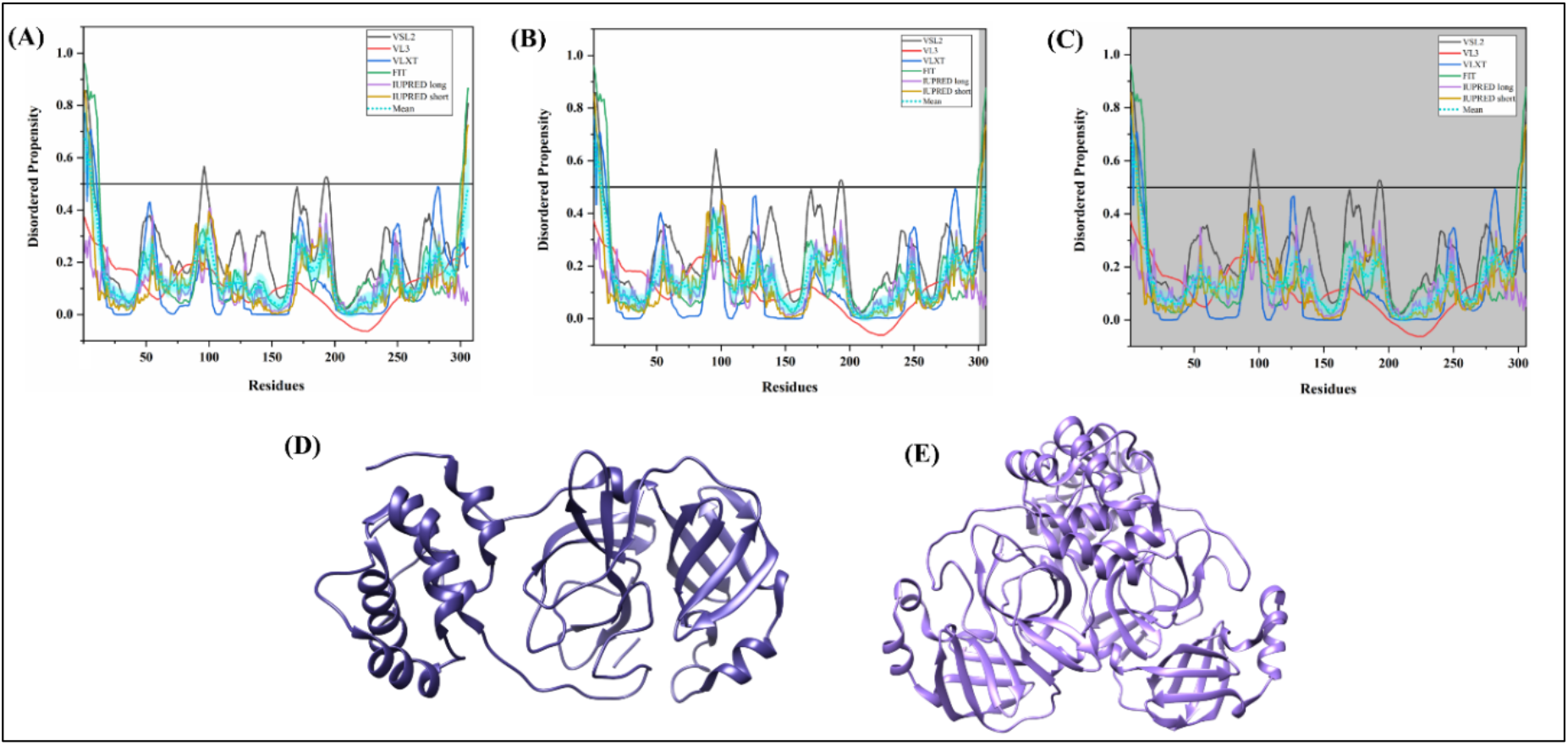
Analysis of intrinsic disorder propensity of Nsp5. Intrinsic disorder profiles generated for Nsp5 from SARS-CoV-2 **(A)**, Human SARS CoV **(B)**, and Bat CoV **(C)**. The colour schemes are the same as in the corresponding plots of **Figure 3. (D)** A 1.50 Å X-ray crystal structure of Nsp5 from Human SARS CoV (PDB ID: 5C5O). **(E)** A 2.16 Å X-ray crystal structure of SARS-CoV-2 Nsp5 (PDB ID: 6LU7).

Nsp5 protein is found to be highly conserved in all three studied CoV viruses. SARS-CoV-2 Nsp5 shares a 96.08% sequence identity with Human SARS Nsp5 and 95.42% with Nsp5 of Bat CoV **(Supplementary Figure S2D)**. Therefore, it not surprising that our analysis demonstrated the identical mean PPID values of 1.96% for Nsp5s from SARS-CoV-2, Human SARS, and Bat CoV (**Table 3**). The predicted per-residue intrinsic disorder propensity of SARS-CoV-2, Human SARS, and Bat CoV Nsp5s are presented in **Figures 23A, 23B**, and **23C**, respectively. As the graphs depict, Nsp5s have several flexible regions and N-terminally IDPR of six residues. Due to the low flexibility of this protein, a single MoRF predicted by MoRFchibi_web is present in the N-terminal region (residues 3-8) in Nsp5s of all three viruses (**Table 2, Supplementary Tables 7** and **8**). Further, the identified nucleotide-binding residues in Nsp5 of all three viruses are tabulated in **Supplementary Tables 9, 10**, and **11**.

#### Non-structural protein 6 (Nsp6)

Nsp6 protein is involved in blocking ER-induced autophagosome/autolysosome vesicle formation that functions in restricting viral production inside host cells. It induces autophagy by activating the omegasome pathway, which is normally utilized by cells in response to starvation. SARS Nsp6 leads to the generation of small autophagosome vesicles thereby limiting their expansion [141].

Nsp6 of SARS-CoV-2 is equally close to Nsp6s from both Human SARS and Bat CoV, having a sequence identity of 87.24% (**Figure 24D**). According to our analysis, mean PPIDs for Nsp6s are calculated to be 1.03%, 1.03%, and 1.03% for SARS-CoV-2, Human SARS CoV, and Bat CoV, respectively. **Figures 24A, 24B**, and **24C** show the corresponding graphs of intrinsic disorder tendency of Nsp6s from SARS-CoV-2, Human SARS CoV, and Bat CoV and demonstrate that these proteins are highly ordered and show low flexibility. As it is a membrane protein, Nsp6 proteins are predicted to have only a single MoRF near the N-terminal region (residues 1-19 in SARS-CoV-2, residues 1-22 in Human SARS, and residues 1-21 in Bat CoV) by the DISOPRED3 server (**Table 2, Supplementary Tables 7** and **8**). The role of these protein-binding regions for the induction of autophagy is need to be elucidated.

**Figure 24.**
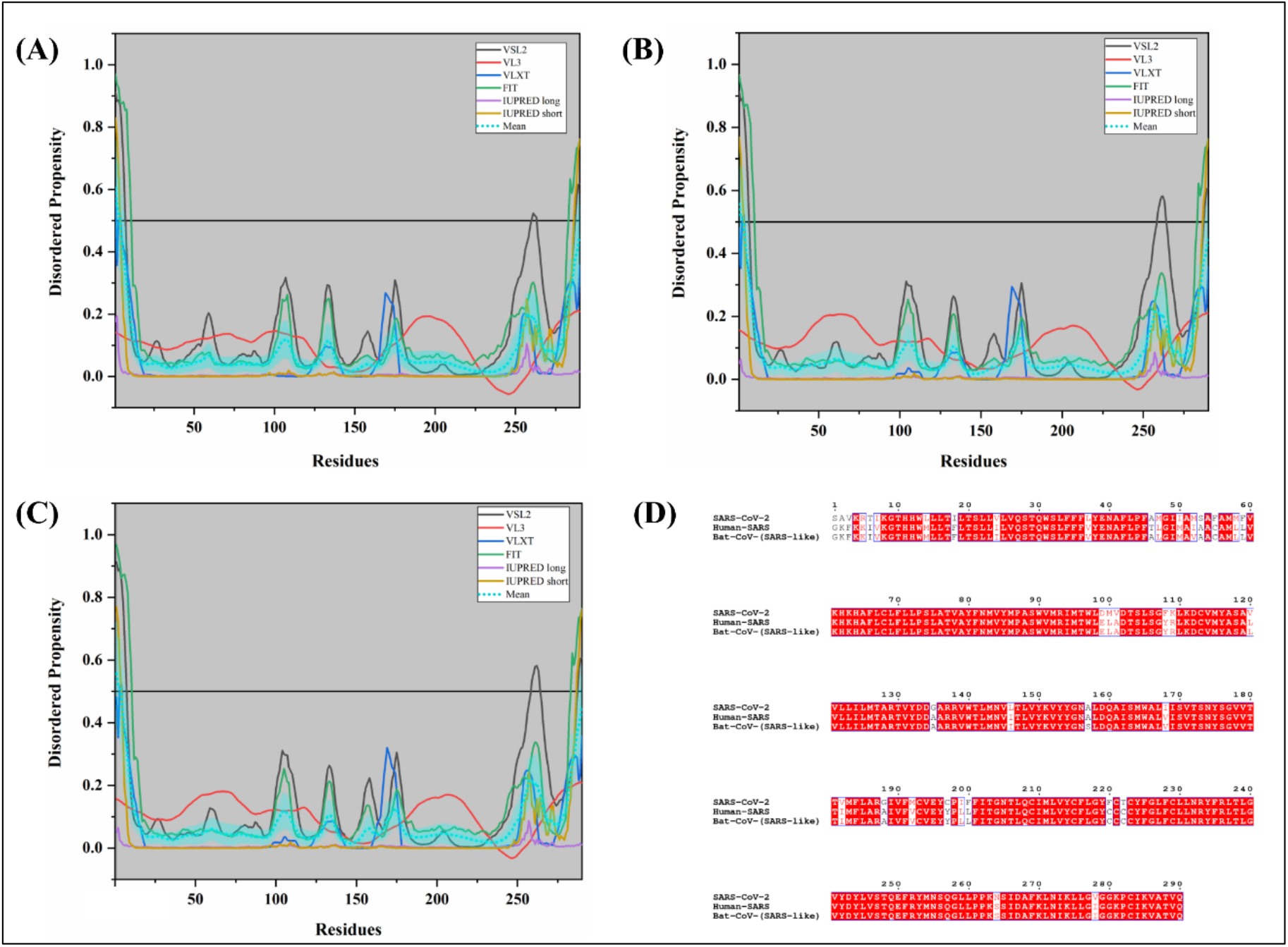
Analysis of intrinsic disorder propensity of Nsp6. Intrinsic disorder profiles generated for Nsp6 from SARS-CoV-2 **(A)**, Human SARS CoV **(B)**, and Bat CoV **(C)**. The colour schemes are the same as in the corresponding plots of **Figure 3. (D)** MSA profile of Nsp6 of SARS-CoV-2, Human SARS, and Bat CoV. Red shaded sequences represent identical regions of these proteins, whereas red characters show similar residues.

#### Non-structural proteins 7 and 8 (Nsp7 and 8)

The ~10 kDa Nsp7 helps in primase-independent *de novo* initiation of viral RNA replication by forming a hexadecameric ring-like structure with Nsp8 protein [142,143]. Both non-structural proteins 7 and 8 contribute 8 molecules to the ring-structured multimeric viral RNA polymerase. Site-directed mutagenesis in Nsp8 revealed a D/ExD/E motif essential for the *in vitro* catalysis [142]. **Figure 25D** depicts the 3.1 Å resolution electron microscopy-based structure (PDB ID: 6NUR) of the RDRP-nsp8-nsp7 complex bound to the Nsp12. The structure identified conserved neutral Nsp7 and Nsp8 binding sites overlapping with finger and thumb domains on Nsp12 of the virus [144].

**Figure 25.**
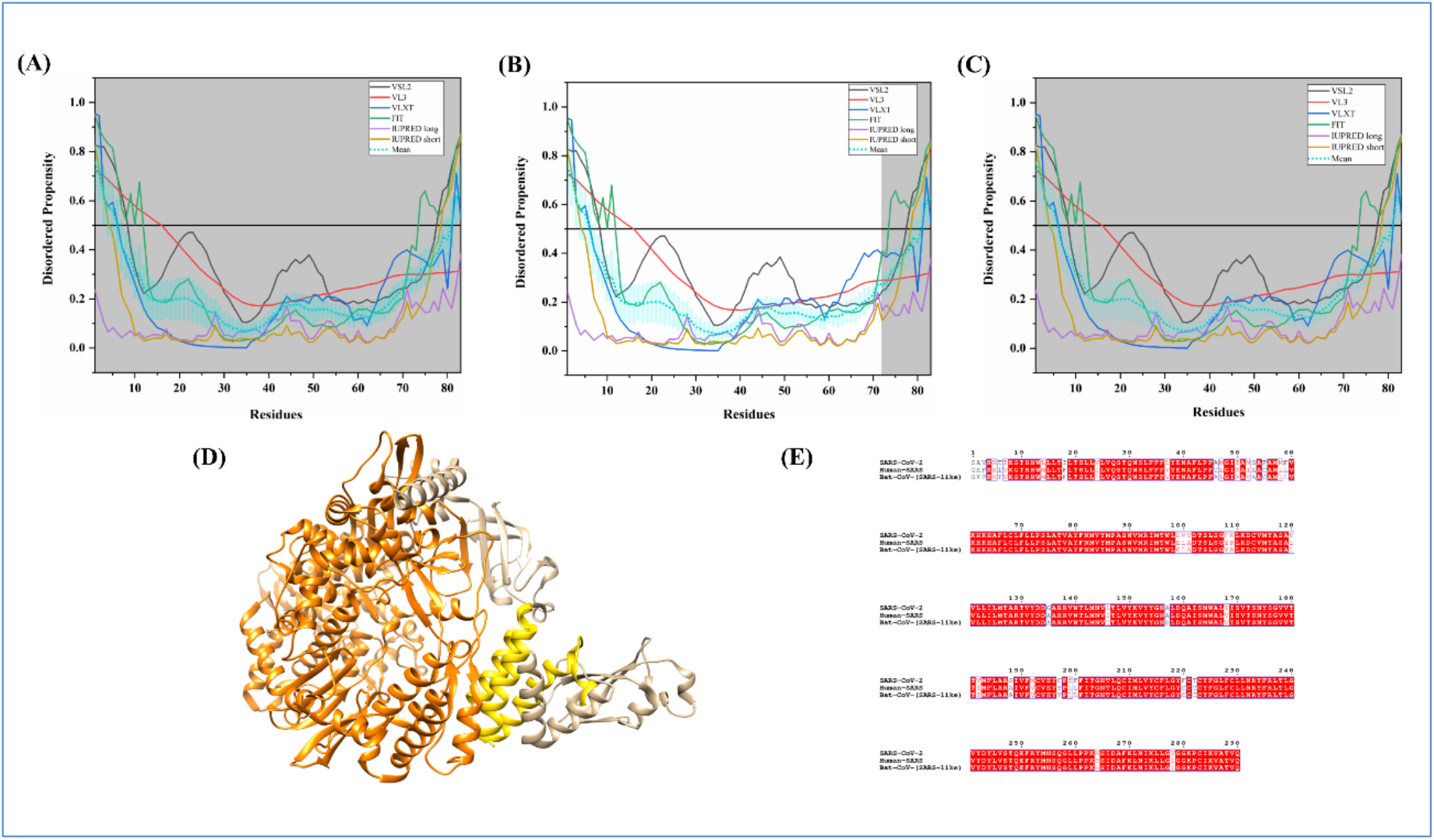
Analysis of intrinsic disorder propensity of Nsp7. Intrinsic disorder profiles generated for Nsp7 from SARS-CoV-2 **(A)**, Human SARS CoV **(B)**, and Bat CoV **(C)**. The colour schemes are the same as in the corresponding plots of **Figure 3. (D)** A 3.10 Å resolution cryo-EM structure of Nsp12-Nsp8-Nsp7 complex (PDB ID: 6NUR). Chain C includes 2-71 residues of Nsp7 (gold colour), chains B and D (dark khaki) represent 77-191 residues of Nsp8 and chain A signifies residues 117-896 and 907-920 of Nsp12 (RNA-directed RNA polymerase) (orange colour) from Human SARS CoV. **(E)** MSA profile of NSP7 of SARS-CoV-2, Human SARS and Bat CoV. Red shaded sequences represent identical regions of these proteins, whereas red characters show similar residues.

We found that Nsp7 of SARS-CoV-2 share 100% sequence identity with Nsp7 of Bat CoV and 98.80% with Nsp7 from Human SARS (**Figure 25E**), while SARS-CoV-2 Nsp8 is closer to Nsp8 of Human SARS (97.47%) than to Nsp8 of Bat CoV (96.46%) (**Figure 26D**).

**Figure 26.**
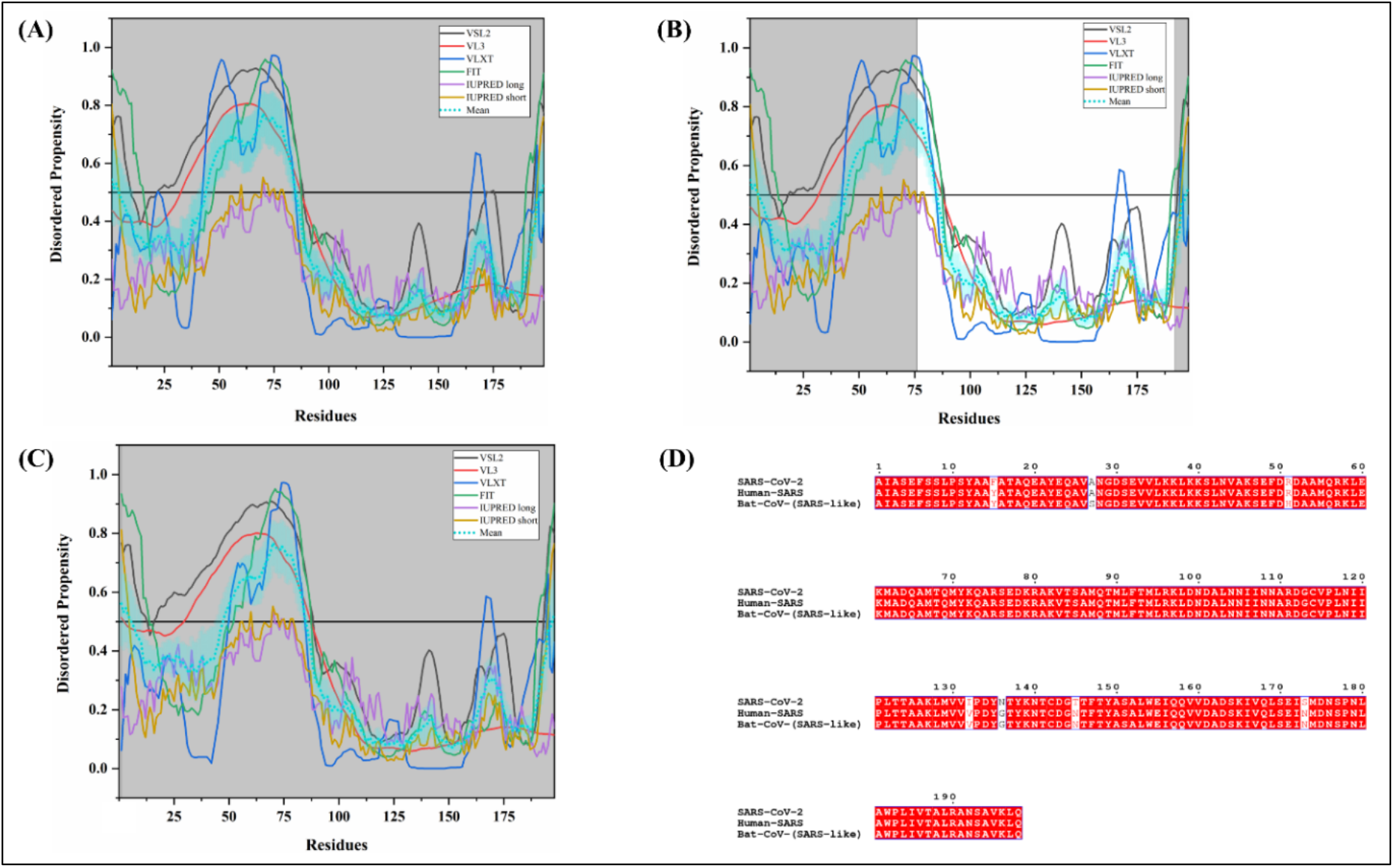
Analysis of intrinsic disorder propensity of Nsp8. Intrinsic disorder profiles generated for Nsp8 from SARS-CoV-2 **(A)**, Human SARS CoV **(B)**, and Bat CoV **(C)**. The colour schemes are the same as in the corresponding plots of **Figure 3. (D)** MSA profile of Nsp8 of SARS-CoV-2, Human SARS and Bat CoV. Red shaded sequences represent identical regions of these proteins, whereas red characters show similar residues.

Due to the high levels of sequence identity, mean PPIDs of all Nsp7s were found to be identical and equal to 9.64%. Both SARS-CoV-2 and Human SARS Nsp8 proteins were calculated to have a mean PPID of 23.74% and, for Nsp8 of Bat CoV mean disorder is predicted to be 22.22%. **Figures 25A, 25B**, and **25C** display the intrinsic disorder profiles for Nsp7s, whereas **Figures 26A, 26B**, and **26C** represent the predicted intrinsic disorder propensity of Nsp8s. As our analysis suggests, Nsp7s might have a well-predicted structure, while Nsp8s are moderately disordered. Nsp8s are predicted to have a long IDPR (residues 44-84) in both SARS-CoV-2 and Human SARS, and a bit shorter IDPR in Bat CoV (residues 48-84). Furthermore, SARS-CoV Nsp7 using its N-terminus residues (V11, C13, V17, and V21) forms a hydrophobic core with Nsp8 residues (M92, M95, L96, M99, and L103). Additionally, H-bonding takes place between Nsp7 Q24 and Nsp8 T89 residues [143]. These amino acids are the part of MoRFs predicted in Nsp7 and Nsp8 proteins. The results are tabulated in both **Table 2**, and **Supplementary Tables 7** and **8**. Three protein binding regions in Nsp7 of SARS-CoV-2 (residues 1-30, 39-58, and 65-83), Human SARS (residues 1-30, 44-58, and 64-83), and Bat CoV (residues 1-30, 39-58, and 65-83) was identified by MoRFchibi_web server. Nsp7 was found with the presence of very few nucleotide-binding regions and Nsp8 contains several DNA as well as RNA binding residues (see **Supplementary Tables 9, 10**, and **11**).

#### Non-structural protein 9 (Nsp9)

Nsp9 protein is a single-stranded RNA-binding protein [145]. It might protect RNA from nucleases by binding and stabilizing viral nucleic acids during replication or transcription [145]. Our results on nucleotide-binding tendency of Nsp9 shows the presence of several RNA binding and few DNA binding residues in Nsp9 of SARS-CoV-2, Human SARS, and Bat CoV (**Supplementary Tables 9, 10**, and **11**). Presumed to evolve from a protease, Nsp9 forms a dimer using its GXXXG motif [146,147]. **Figure 27D** shows a 2.7 Å crystal structure of the homodimer of Human SARS Nsp9 (PDB ID: 1QZ8) that identified a unique and previously unreported for other proteins, oligosaccharide/oligonucleotide fold-like fold [145]. Here, each monomer contains a cone-shaped β-barrel and a C-terminal α-helix arranged into a compact domain [145].

**Figure 27.**
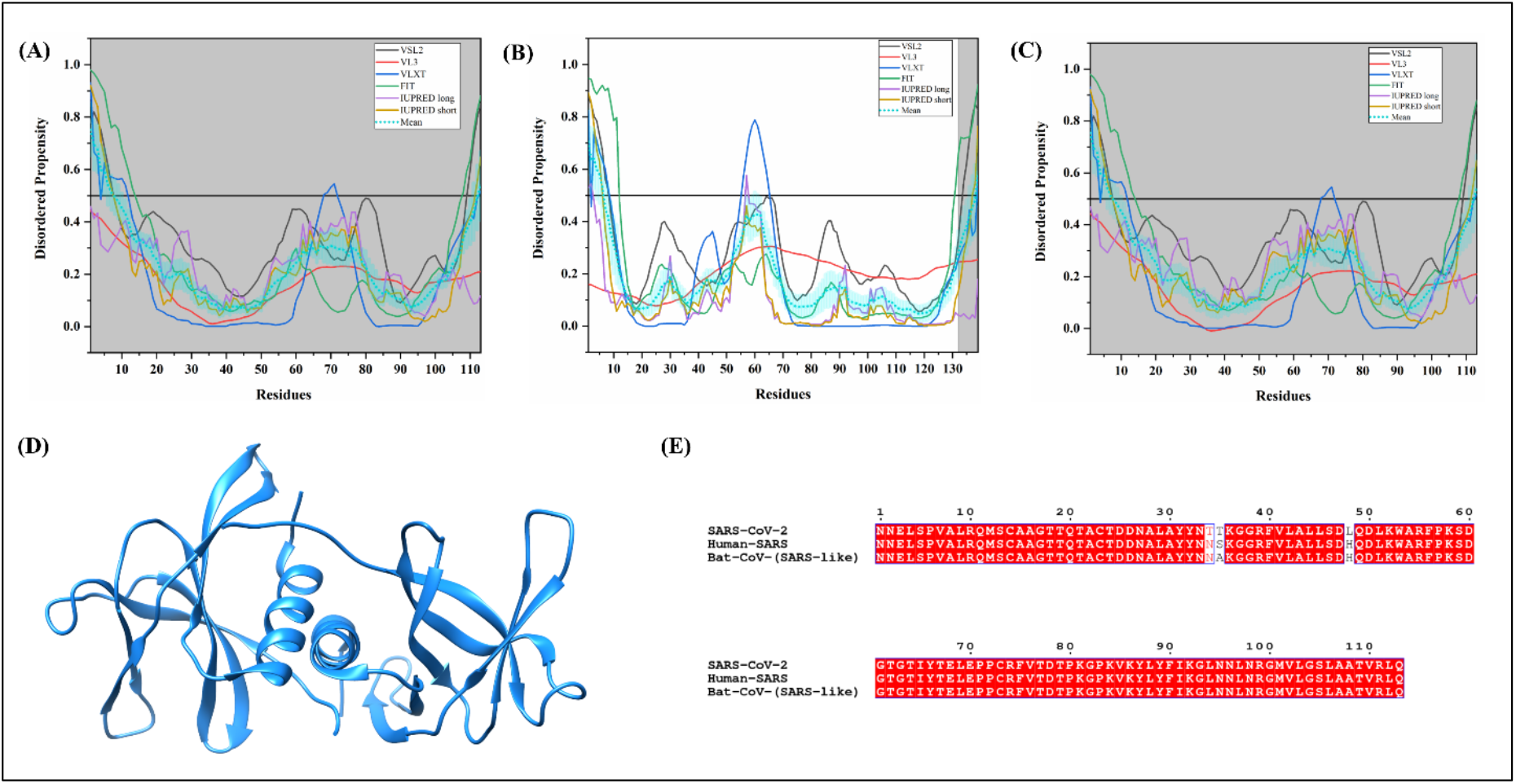
Analysis of intrinsic disorder propensity of Nsp9. Intrinsic disorder profiles generated for Nsp9 from SARS-CoV-2 **(A)**, Human SARS CoV **(B)**, and Bat CoV **(C)**. The colour schemes are the same as in the corresponding plots of **Figure 3. (D)** A 2.70 Å X-ray crystal structure of 3-113 residues of Nsp9 from Human SARS CoV (PDB ID: 1QZ8). **(E)** MSA profile of Nsp9s from SARS-CoV-2, Human SARS, and Bat CoV. Red shaded sequences represent identical regions of these proteins, whereas red characters show similar residues.

Nsp9 of SARS-CoV-2 is equally similar to Nsp9s from both Human SARS and Bat CoV, having a percentage identity of 97.35%. The difference in three amino acids at 34, 35 and 48 positions accounts for these similarity scores (**Figure 27E**). As calculated, the mean PPIDs of Nsp9s of SARS-CoV-2, Human SARS CoV, and Bat CoV are 7.08%, 7.96%, and 7.08% respectively. **Figures 27A, 27B**, and **27C** depict the predicted intrinsic disorder propensity in the Nsp9 protein from SARS-CoV-2, Human SARS, and Bat CoV. According to our analysis, all three Nsp9s are rather structured but contain flexible regions. Nsp9 contains conserved residues (R10, K52, Y53, R55, R74, F75, K86, Y87, F90, K92, R99, and R111) of positively charged side chains suitable for binding with the negatively charged phosphate backbone of RNA and aromatic side-chain amino acids providing stacking interactions [145].

These residues are a part of multiple disorder-based binding sites predicted by MoRFchibi_web server (**Table 2, Supplementary Table 7** and **8**).

#### Non-structural protein 10 (Nsp10)

Nsp10 performs several functions for SARS-CoV. It forms a complex with Nsp14 for dsRNA hydrolysis in 3′ to 5′ direction and activates its exonuclease activity [148]. It also stimulates the methyltransferase (MTase) activity of Nsp14 required during RNA-cap formation after replication [149]. **Figure 28D** represents the X-ray crystal structure of the Nsp10/Nsp14 complex (PDB ID: 5C8T) [150]. In agreement with the results of previous biochemical experimental studies, the structure identified important interactions with the ExoN (exonuclease domain) of Nsp14 without affecting its N7-MTase activity [148,149].

**Figure 28.**
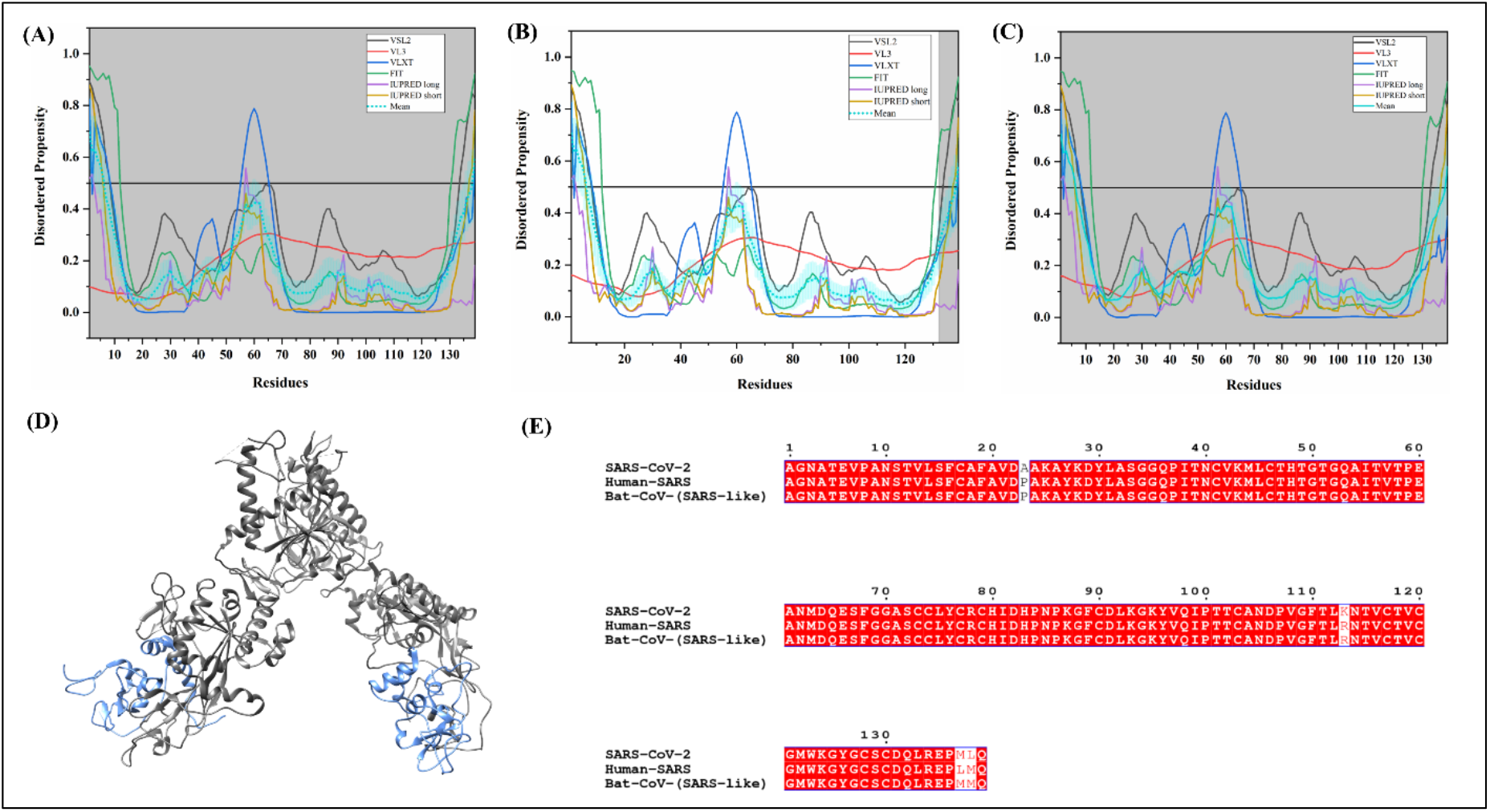
Analysis of intrinsic disorder propensity of Nsp10. Intrinsic disorder profiles generated for Nsp10 from SARS-CoV-2 **(A)**, Human SARS CoV **(B)**, and Bat CoV **(C)**. The colour schemes are the same as in the corresponding plots of **Figure 3. (D)** A 3.20 Å X-ray crystal structure of the Nsp10/Nsp14 complex (PDB ID: 5C8T). In this structure, A and C chain (cornflower blue colour) signify show 1-131 residues of Nsp10 and B and D chains correspond to residues 1-453 and 465-525 of Nsp14 (dim grey colour). **(E)** MSA profile of Nsp10s from SARS-CoV-2, Human SARS and Bat CoV. Red shaded sequences represent identical regions of these proteins, whereas red characters show similar residues.

SARS-CoV-2 Nsp10 protein is quite conserved having a 97.12% sequence identity with Nsp10 of Human SARS and 97.84% with Nsp10 of Bat CoV (**Figure 28E**). Mean PPIDs of all three studied Nsp10 proteins are found to be 5.04%. **Figures 28A, 28B**, and **28C** represent disorder profiles of Nsp10s and signify the lack of long IDPRs but presence flexible regions in these proteins. Furthermore, **Table 2**, and **Supplementary Table 7** and **8** shows that all three virus Nps10 has multiple MoRFs. For SARS-CoV-2, three MoRFs (residues 25-32, 91-99, and 133-138) was identified by MoRFchibi_web server and one MoRF (residues 11-18) was predicted by MoRFPred server. Interestingly, the SARS-CoV Nsp10 residues F16, F19, and V21 form van der Waals interactions with many of the Nsp14 amino acids [150] and one residue (F16) is located in MoRF region which we have identified. Furthermore, many nucleotide-binding residues which are found in all three viruses (**Supplementary Table 9, 10**, and **11**) and above-mentioned residues are not found to interact with DNA/RNA.

#### Non-structural protein 12 (Nsp12)

In coronaviruses, Nsp12 is an RNA-dependent RNA Polymerase (RDRP). It carries out both primer-independent and primer-dependent synthesis of viral RNA with Mn^2+^ as its metallic co-factor and viral Nsp7 and 8 as protein co-factors [151]. As aforementioned, a 3.1Å resolution structure of Human SARS Nsp12 in association with Nsp7 and Nsp8 proteins (PDB ID: 6NUR) has been reported using electron microscopy (**Figure 25D)**. Nsp12 has a polymerase domain similar to “right hand”, finger domain (398–581, 628–687 residues), palm domain (582–627, 688–815 residues) and a thumb domain (816–919) [144].

SARS-CoV-2 Nsp12 protein has a highly conserved C-terminal region **(Supplementary Figure S2E)**. It is found to share a 96.35% sequence identity with Human SARS Nsp12 and 95.60% with Bat CoV Nsp12. Mean PPID values for all three Nsp12s are estimated to be 0.43% (**Table 3**). **Figures 29A, 29B**, and **29C** show that although these proteins are mostly ordered, they have multiple flexible regions. As RDRP protein is observed to be mostly structured, significant MoRFs in disordered regions are not found (**Table 2, Supplementary Table 7** and **8**).

**Figure 29.**
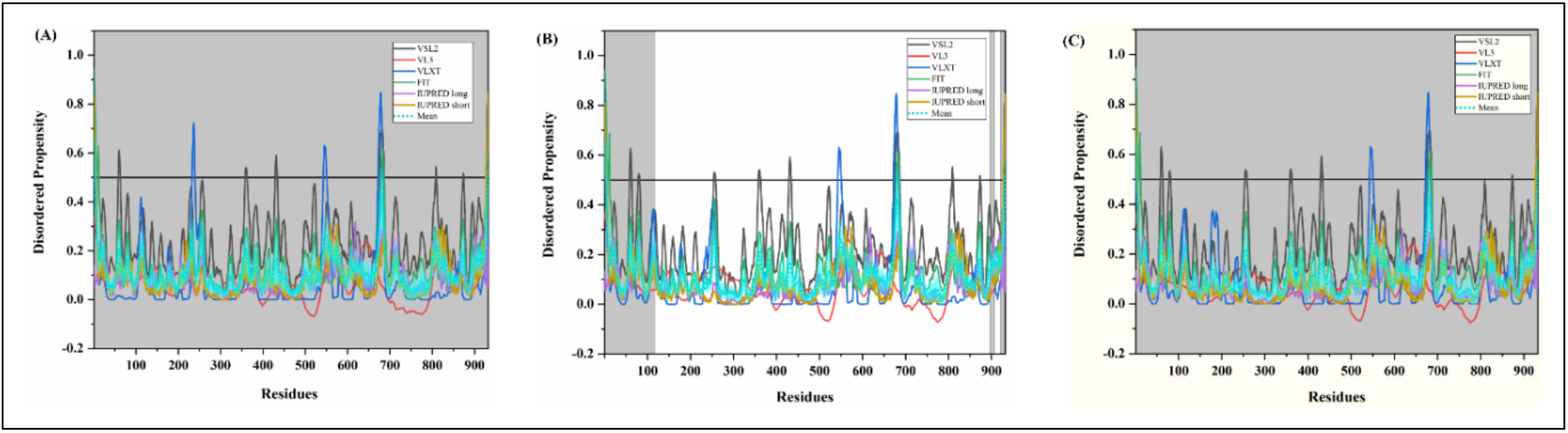
Analysis of intrinsic disorder propensity of Nsp12. Intrinsic disorder profiles generated for Nsp12 from SARS-CoV-2 **(A)**, Human SARS CoV **(B)**, and Bat CoV **(C)**. The colour schemes are the same as in the corresponding plots of **Figure 3**.

#### Non-structural protein 13 (Nsp13)

Nsp13 functions as a viral helicase and unwinds dsDNA/dsRNA in 5’ to 3’ direction [152]. Recombinant viral helicase expressed in *E.coli* Rosetta 2 strain was reported to unwind ~280 bp per second [152]. **Figure 30D** represents a 2.8 Å X-ray crystal structure of Human SARS Nsp13 (PDB ID: 6JYT) [153]. This helicase contains a *β*19-*β*20 loop on 1A domain, which is primarily responsible for its unwinding activity. Furthermore, the study revealed an important interaction of Nsp12 with Nsp13 that further enhances its helicase activity [153].

**Figure 30.**
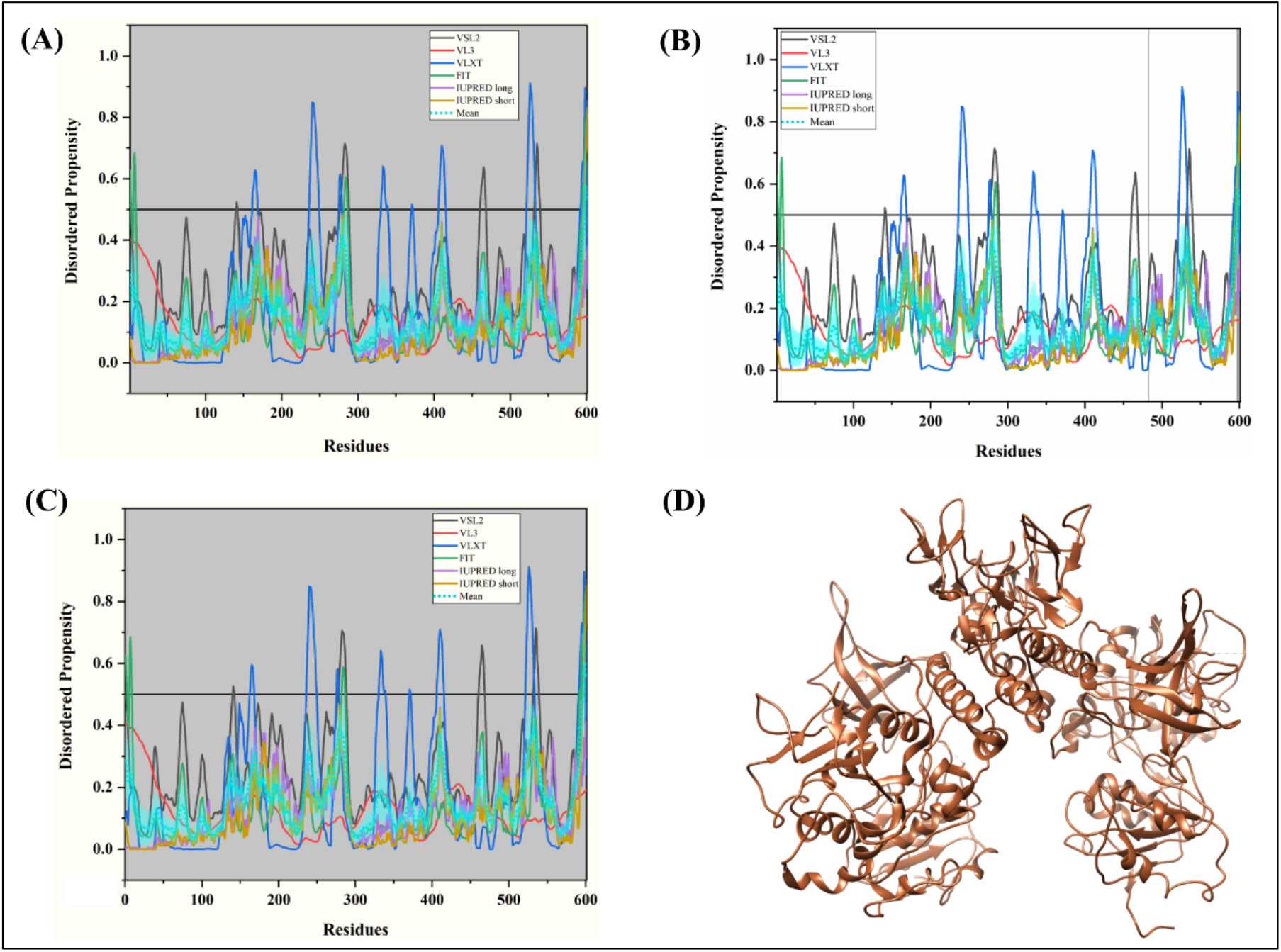
Analysis of intrinsic disorder propensity of Nsp13. Intrinsic disorder profiles generated for Nsp13 from SARS-CoV-2 **(A)**, Human SARS CoV **(B)**, and Bat CoV **(C)**. The colour schemes are the same as in the corresponding plots of **Figure 3. (D)** A 2.80 Å X-ray crystal structure of 1-596 residues of Human SARS Nsp13 (PDB ID: 6JYT).

The 601-amino-acid-long Nsp13 of SARS-CoV-2 is almost completely conserved, as it shares 99.83% with Nsp13 of Humans SARS and 98.84% with Nsp13 of Bat CoV **(Supplementary Figure S2F)**. In accordance with our results, the mean PPIDs of all three Nsp13 proteins are estimated to be 0.67%. **Figures 30A, 30B**, and **30C** show that Nsp13s contain multiple flexible regions but do not possess significant disorder. As expected, being a low disorder protein Nsp13 does not contain any MoRF region and not a single binding-region is located by any server used in all three viruses (**Table 2, Supplementary Table 7** and **8**). It has many nucleotide-binding residues (RNA and DNA) which are tabulated in **Supplementary Tables 9, 10**, and **11**.

#### Non-structural protein 14

Nsp14 is a multifunctional viral protein that acts as an Exoribonuclease (ExoN) and methyltransferase (N7-MTase) in SARS coronaviruses. It’s 3’ to 5’ exonuclease activity lies in the conserved DEDD residues related to exonuclease superfamily [154]. Its guanine-N7 methyltransferase activity depends upon the S-adenosyl-L-methionine (AdoMet) as a cofactor [149]. As aforementioned, Nsp14 requires Nsp10 for activating its ExoN and N7-MTase activity inside the host cells. **Figure 28D** depicts the 3.2 Å crystal structure of human SARS nsp10/nsp14 complex (PDB ID: 5C8T), where amino acids 1-287 form the ExoN domain and 288-527 residues form the N7-MTase domain of nsp14. A loop (residues 288-301) is essential for its N7-MTase activity [150].

SARS-CoV-2 Nsp14 protein shares a 95.07% identity with Human SARS Nsp14 and 94.69% with Bat CoV Nsp14 (**Supplementary Figure S2G**). Mean PPID values for Nsp14s from SARS-CoV-2 and Human SARS is calculated to be 0.38%, while the Nsp14 from Bat CoV has a mean PPID 0.57%. Predicted per-residue intrinsic disorder propensity of Nsp14s from SARS-CoV-2, Human SARS, and Bat CoV is represented in **Figures 31A, 31B**, and **31C**, respectively. As can be observed from these plots and corresponding PPID values, all Nsp14s are found to be highly structured. Likewise, **Table 2** shows Nsp14 contains two protein binding regions (residues 8-13 and 441-445) predicted by the MoRFPred server in all three viruses. As shown in **Supplementary Tables 9, 10**, and **11**, the ORF14 represents multiple nucleotide-binding residues.

**Figure 31.**
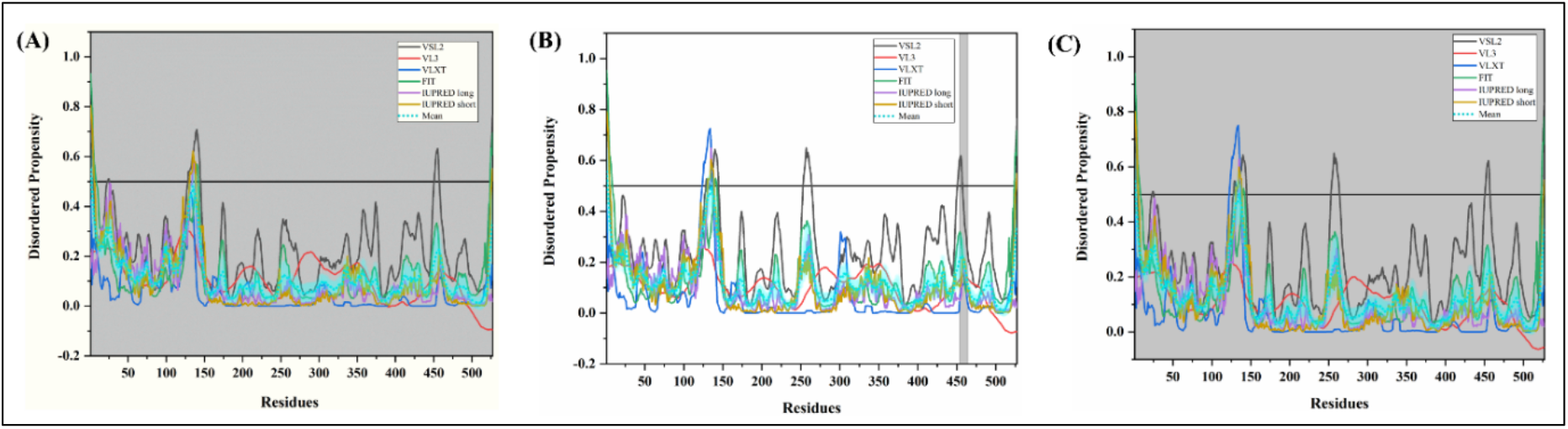
Analysis of intrinsic disorder propensity of Nsp14. Intrinsic disorder profiles generated for Nsp14 from SARS-CoV-2 **(A)**, Human SARS CoV **(B)**, and Bat CoV **(C)**. The colour schemes are the same as in the corresponding plots of **Figure 3**.

#### Non-structural protein 15 (Nsp15)

Nsp15 is a uridylate-specific RNA Endonuclease (NendoU) that creates a 2′–3′ cyclic phosphates after cleavage. Its endonuclease activity depends upon Mn^2+^ ions as co-factors. Conserved in Nidovirus, it acts as an important genetic marker due to its absence in other RNA viruses [155]. **Figure 32D** represents a 2.6 Å crystal structure of Uridylate-specific Nsp15 (PDB ID: 2H85) that was deduced by Bruno and colleagues using X-ray diffraction [156]. The monomeric Nsp15 has three domains: N-terminal domain (1-62 residues) formed by a three anti-parallel *β*-strands and two α-helices packed together; a middle domain (63–191 residues) that contains an α-helix connected via a 39-amino-acid-long coil to an ordered region containing two α-helices and five *β*-strands; and a C-terminal domain (192–345 residues) consisting of two anti-parallel three *β*-strand sheets on each side of a central α-helical core [156].

**Figure 31.**
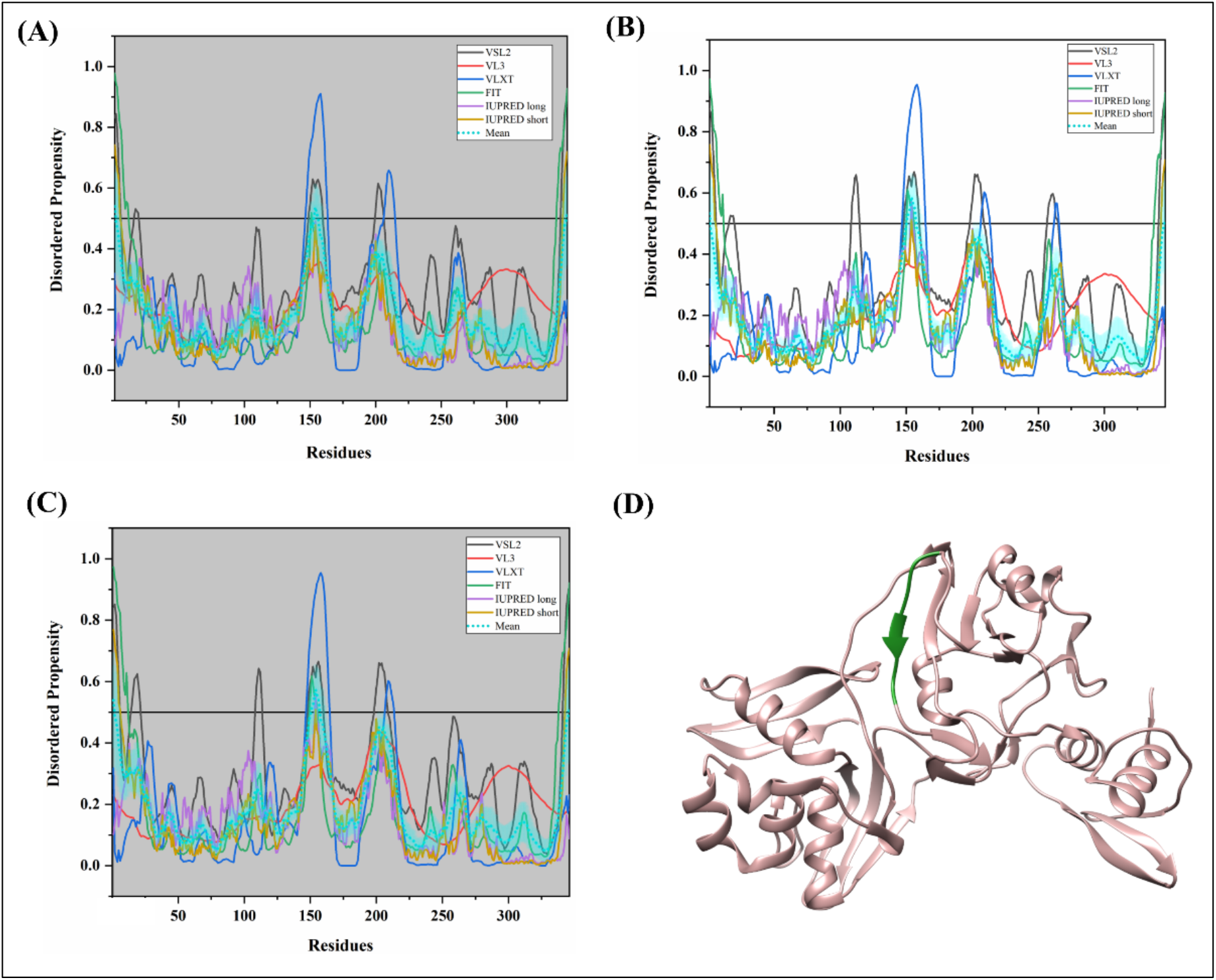
Analysis of intrinsic disorder propensity of Nsp15. Intrinsic disorder profiles generated for Nsp15 from SARS-CoV-2 **(A)**, Human SARS CoV **(B)**, and Bat CoV **(C)**. The colour schemes are the same as in the corresponding plots of **Figure 3. (D)** A 2.60 Å X-ray crystal structure of Nsp15 from Human SARS CoV (PDB ID: 2H85) (rosy brown colour). Residues 151-157 predicted to be disordered are represented in forest-green colour.

The Nsp15 is found to be quite conserved across human SARS and bat CoVs. SARS-CoV-2 Nsp15 shares an 88.73% sequence identity with Nsp15 of Human SARS and 88.15% with Nsp15 of Bat CoV (**Supplementary Figure S2H**). Calculated mean PPIDs of Nsp15s from SARS-CoV-2, Human SARS, and Bat CoV are 1.73%, 2.60%, and 2.60%, respectively. Similar to many other non-structural proteins of coronaviruses, Nsp15s from SARS-CoV-2, Human SARS, and Bat CoV are predicted to possess multiple flexible regions but contain virtually no IDPRs (see **Figures 32A, 32B**, and **32C**). Similarly, no significant disorder-binding regions are predicted in Nsp15 proteins (**Table 2**). SARS-CoV-2 contain one MoRF (residues 9-13) predicted by MoRFPred server. Human SARS do not have a single MoRF while Bat CoV possesses two very short binding regions (**Supplementary Table 7** and **8**). **Supplementary Table 9, 10**, and **11** depicts the presence of many RNA binding residues and few DNA binding residues in Nsp15 of all three viruses.

#### Non-structural protein 16 (Nsp16)

Nsp16 protein is another MTase domain-containing protein. As methylation of coronavirus mRNAs occurs in steps, three proteins Nsp10, Nsp14, and Nsp16 acts one after another. The first event requires the initiation trigger from Nsp10 protein, after which Nsp14 methylates capped mRNAs forming cap-0 (7Me) GpppA-RNAs. Nsp16 protein, along with its co-activator protein Nsp10, acts on cap-0 (7Me) GpppA-RNAs to give rise to final cap-1 (7Me)GpppA(2’OMe)-RNAs [149,157]. A 2 Å X-ray crystal structure of the Human SARS nsp10-nsp16 complex is depicted in **Figure 33D**(PDB ID: 3R24) [158]. The structure consists of a characteristic fold present in class I MTase family comprising of α-helices and loops surrounding a seven-stranded β-sheet [158].

**Figure 33.**
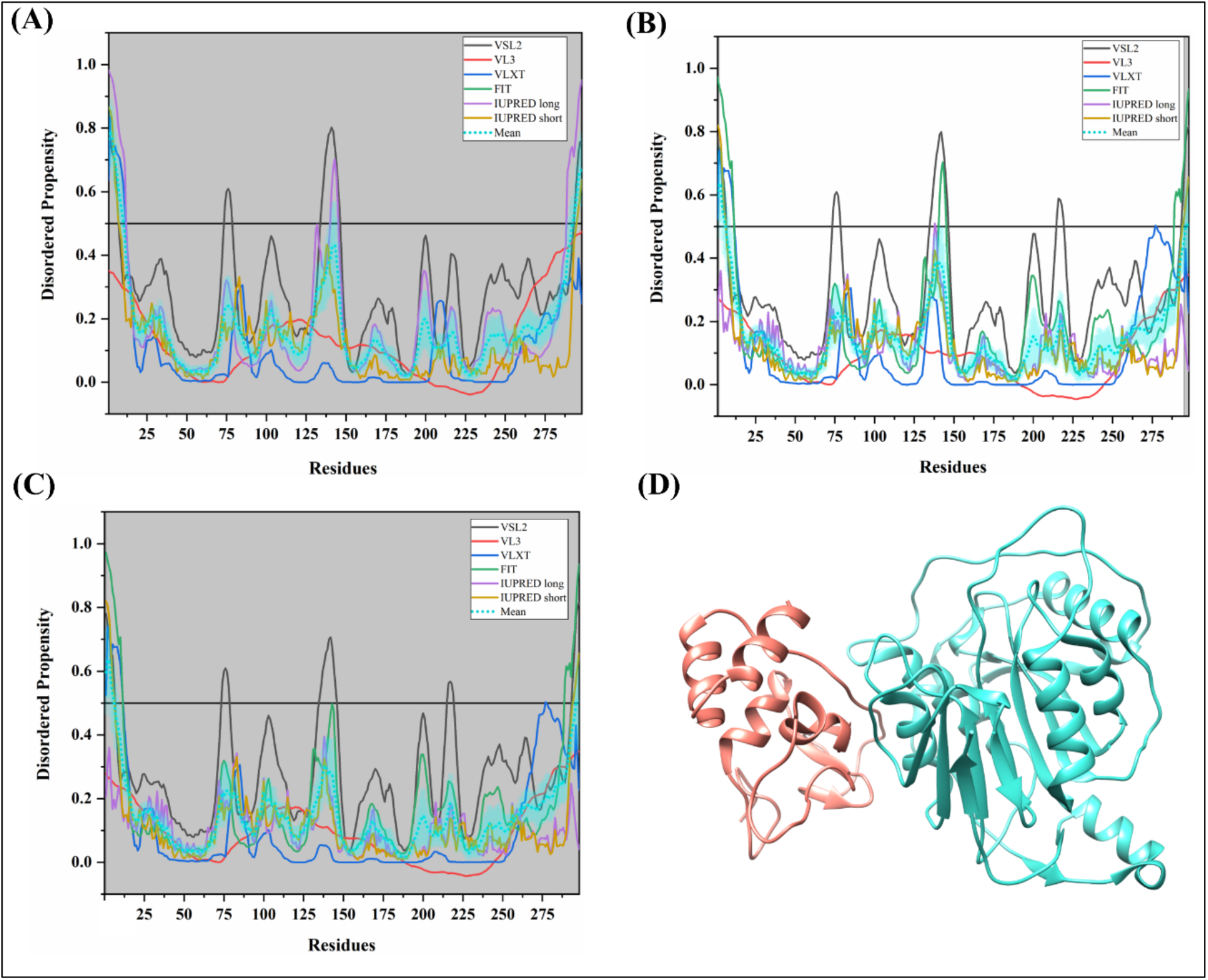
Analysis of intrinsic disorder propensity of Nsp16. **(A)** Intrinsic disorder profiles generated for Nsp15 from SARS-CoV-2 **(A)**, Human SARS CoV **(B)**, and Bat CoV **(C)**. The colour schemes are the same as in the corresponding plots of **Figure 3. (D)** A 2.60 Å X-ray crystal structure of the Human SARS Nsp10-Nsp16 complex (PDB ID: 3R24). Chain A shown in turquoise colour corresponds to the residues 3-294 of Nsp16

Nsp16 protein of SARS-CoV-2 is found to be equally similar to Nsp16s from Human SARS and Bat CoV (93.29%) (**Supplementary Figure S2I**). Mean PPIDs for Nsp16s from SARS-CoV-2, Human SARS, and Bat CoV are 5.37%, 3.02%, and 3.02%, respectively. In line with these PPIDs values, **Figures 33A, 33B**, and **33C** show that Nsp16s are mostly ordered proteins containing several flexible regions. Correspondingly, no significant MoRFs are present in this protein (**Table 2, Supplementary Table 7** and **8**). A single MoRF (residues 151-156) were found with the help of MoRFPred in all three viruses. Further, several RNA-binding and few DNA-binding residues are also identified (**Supplementary Table 9, 10, and 11**).

#### Replicase polyprotein 1a

Since replicase polyprotein 1a contains non-structural proteins 1-10 identical to those found in replicase polyprotein 1ab, we did not perform their disorder analysis separately. However, replicase polyprotein 1a has one additional non-structural protein designated as Nsp11.

#### Non-structural protein 11 (Nsp11)

Nsp11 is a small uncharacterized protein cleaved from the replicase polyprotein 1a. This small protein with unknown function requires experimental insights to further characterize this protein. The intrinsic disorder predicting software used in this study requires amino acid sequences, which are at least 30-residue long. Therefore, because of their short sequences (just 13 residues) Nsp11s from all three studied coronaviruses were not checked for the intrinsic disorder, disorder-based protein binding regions, and nucleotide-binding residues. Based on the MSA outputs, Nsp11 from SARS-CoV-2 was found to have a sequence identity of 84.62% with Nsp11s from Human SARS and Bat CoV (**Figure 34**).

**Figure 34.**
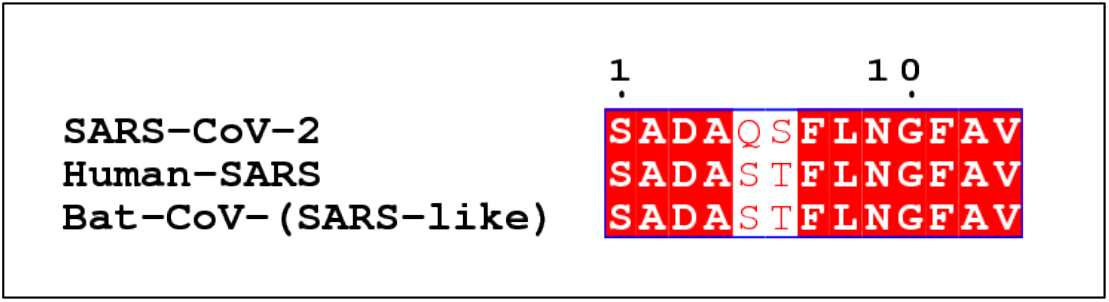
MSA profile of Nsp11 of SARS-CoV-2, Human SARS and Bat CoV. Red shaded sequences represent identical regions of these proteins, whereas red characters show similar residues.

### Concluding remarks

The emergence of new viruses and associated deaths around the globe represent one of the major concerns of modern times. Despite its pandemic nature, there is very little information available in the public domain regarding the structures and functions of SARS-CoV-2 proteins. Based on its similarity with Human SARS CoV and Bat CoV, the published reports have suggested the functions of SARS-CoV-2 proteins. In this study, we utilized information available on SARS-CoV-2 genome and translated proteome from GenBank, and carried out a comprehensive computational analysis of the prevalence of the intrinsic disorder in SARS-CoV-2 proteins. Additionally, a comparison was also made with proteins from close relatives of SARS-CoV-2 from the same group of beta coronaviruses, Human SARS CoV and Bat CoV. Our analysis revealed that in these three CoVs, the N proteins are highly disordered, possessing the PPID values of more than 60%. These viruses also have several moderately disordered proteins, such as Nsp8, Orf6, and Orf9b. Although other proteins have shown lower disorder content, almost all of them contain at least some IDPRs, and all CoV proteins analysed in this study definitely have multiple flexible regions. Importantly, our study provides novel information on presence of intrinsic disorder at the cleavage sites of the replicase 1ab polyprotein of CoVs. This observation confirms the crucial role of IDPRs in maturation of individual proteins. We also established that many of these proteins contain disorder-based binding motifs. Since IDPs/IDPRs might undergo structural transition upon association with their physiological partners, our study generates important grounds for better understanding of the functionality of these proteins, their interactions with other viral proteins, as well as interaction with host proteins in different physiological conditions. This will also guide structural biologists to carry out a structure-based analysis of SARS-CoV-2 proteome to explore the path for the development of new drugs and vaccines.

### Future perspective

The periodical outbreaks of pathogens worldwide always remind the lack of suitable drugs or vaccines for proper cure or treatment. In 2003, nearly 750 deaths were reported due to the SARS outbreak in more than 24 countries. But this time, the outbreak of Wuhan’s novel coronavirus (SARS-CoV-2) has quickly surpassed this number, indicating more causalities soon. The lack of accurate information and ignorance of primary symptoms are major reasons, which cause many infection cases. Although efficient transmission from human to human is confirmed, the actual reasons for fast SARS-CoV-2 spread are still unknown, but some assumptions were made by researchers and Chinese authorities. The fast spread of SARS-CoV-2, COVID-19 pandemic, and associated introduction of quarantine also have made major impacts on economy and education worldwide due to several restrictions, such as limited transportation, restrained or frozen traveling, halted attendance of mass events, the introduction of distant teaching and learning, etc. Due to advancements in sequencing techniques, the full genome sequence of SARS-CoV-2 was made available in a few days of the first infection report from Wuhan, China. However, massive subsequent research needs to be done to identify the actual cause of SARS-CoV-2 infectivity and to design suitable treatment in the coming future. Certain possibilities can be explored with the available information. The mutational pressure study on this virus will be very interesting to see if this virus transforms from Bat SARS to Human SARS to SARS-CoV-2. More in-depth experimental studies using molecular and cell biology techniques to establish structure-function relationships are required for a better understanding of the functioning of SARS-CoV-2 proteins. Additionally, based on the sequence homology and information on protein-protein interactions, the associated viral and host proteins should be explored, for finding means suitable for limiting replication, maturation, and ultimately pathogenesis of this virus. Although structural biology techniques (so-called rational drug design) can be used in drug development utilizing high throughput screening of compounds virtually or experimentally, the applicability of these techniques is limited by the presence of intrinsic disorder in target proteins. Therefore, the thorough disorder analysis of three coronaviruses conducted in this study will help structural biologists to rationally design experiments keeping this information in mind.

## List of Abbreviations

ACE2: Angiotensin-converting enzyme 2
CDF: Cumulative distribution function
CH: Charge hydropathy
COVID-19: Coronavirus disease 2019
CTD: Cterminal domain
DMVs: Double-membrane vesicles
ICTV: International committee on taxonomy of viruses
IDP: Intrinsically disordered proteins
IDPRs: Intrinsically disordered protein regions
IFN: Interferon
MoRFs: Molecular recognition features
MSA: Multiple sequence alignment
Nsps: Non-structural proteins
NTD: N-terminal domain
PONDR: Predictor of natural disordered regions
PPID: Predicted percentage of intrinsic disorder
Pprint: Prediction of Protein RNA-Interaction
RBD: Receptor binding domain
SARS: Severe acute respiratory syndrome
TRS: Transcriptional regulatory sequences
VLPs: Virus-like particles
WHO: World health organization

## Authors Contribution

RG: Conception and design, interpretation of data, writing, and review of the manuscript, and study supervision. VNU: Conception and design, acquisition and interpretation of data, writing, and review of the manuscript. MS, TB, PK, BRG, KG: acquisition and interpretation of data, writing of the manuscript.

## Acknowledgements

All the authors would like to thank IIT Mandi for providing facilities. MS and BRG were supported by MHRD for funding. KG was supported by the Department of Biotechnology (DBT), India (BT/PR16871/NER/95/329/2015). PK was supported by IIT Mandi-IIT Ropar-PGI Chandigarh, BioX consortium grant (IITM/INT/RG/18). TB is grateful to the Department of Science and Technology for her INSPIRE fellowship for Funding.

## Supplementary Figure legends

**Supplementary Table 1.** Evaluation of intrinsic disorder in structural and accessory proteins of SARS-CoV-2.

**Supplementary Table 2.** Evaluation of intrinsic disorder in structural and accessory proteins of Human SARS.

**Supplementary Table 3.** Evaluation of intrinsic disorder in structural and accessory proteins of Bat CoV.

**Supplementary Table 4.** Evaluation of intrinsic disorder in non-structural proteins SARS-CoV-2.

**Supplementary Table 5.** Evaluation of intrinsic disorder in non-structural proteins of Human SARS.

**Supplementary Table 6.** Evaluation of intrinsic disorder in non-structural proteins of Bat CoV.

**Supplementary Table 7:** Predicted MoRF residues in Human SARS proteins.

**Supplementary Table 8:** Predicted MoRF residues in Bat CoV proteins.

**Supplementary Table 9:** Predicted nucleotide-binding residues in SARS-CoV-2 proteins.

**Supplementary Table 10:** Predicted nucleotide-binding residues in Human SARS proteins.

**Supplementary Table 11:** Predicted nucleotide-binding residues in BAT CoV proteins.

**Supplementary Figures S1.** Multiple sequence alignment of structural proteins of all three studied coronaviruses are generated using Clustal Omega. The aligned images are created using Esprit 3.0.

**Figure S1A.** MSA of SARS-CoV-2, Human SARS, and Bat CoV spike glycoproteins.

**Figure S1B.** MSA of SARS-CoV-2, Human SARS, and Bat CoV Nucleoproteins.

**Supplementary Figure S2.** Multiple sequence alignment of non-structural proteins of all three studied coronaviruses are generated using Clustal Omega. The aligned images are created using Esprit 3.0.

**Figure S2A.** MSA of SARS-CoV-2, Human SARS, and Bat CoV Nsp2 proteins.

**Figure S2B.** MSA of SARS-CoV-2, Human SARS, and Bat CoV Nsp3 proteins.

**Figure S2C.** MSA of SARS-CoV-2, Human SARS, and Bat CoV Nsp4 proteins.

**Figure S2D.** MSA of SARS-CoV-2, Human SARS, and Bat CoV Nsp5 proteins.

**Figure S2E.** MSA of SARS-CoV-2, Human SARS, and Bat CoV Nsp12 proteins.

**Figure S2F.** MSA of SARS-CoV-2, Human SARS, and Bat CoV Nsp13 proteins.

**Figure S2G.** MSA of SARS-CoV-2, Human SARS, and Bat CoV Nsp14 proteins.

**Figure S2H.** MSA of SARS-CoV-2, Human SARS, and Bat CoV Nsp15 proteins.

**Figure S2I.** MSA of SARS-CoV-2, Human SARS, and Bat CoV Nsp16 proteins.

