## Supplementary Files for "When Darkness Becomes a Ray of Light in the Dark Times: Understanding the COVID-19 via the Comparative Analysis of the Dark Proteomes of SARS-CoV-2, Human SARS and Bat SARS-Like Coronaviruses"

**Supplementary Table 1.** Evaluation of intrinsic disorder in structural and accessory proteins of SARS-CoV-2.

| **Sr. No.** | **Protein** | **Length** | **PPID_VSL2** | **PPID_VL3** | **PPID_VLXT** | **PPID_FIT** | **PPID_IUPRED_Long** | **PPID_IUPRED_Short** | **PPID_Mean** |
| --- | --- | --- | --- | --- | --- | --- | --- | --- | --- |
| **1.** | Spike Glycoprotein | 1273 | 8.01 | 3.38 | 7.70 | 3.53 | 0.00 | 0.47 | 1.41 |
| **2.** | Envelope | 75 | 25.33 | 0.00 | 12.00 | 29.33 | 0.00 | 8.00 | 5.33 |
| **3.** | Membrane | 222 | 11.26 | 0.00 | 5.86 | 12.61 | 0.00 | 5.41 | 2.70 |
| **4.** | Nucleocapsid | 419 | 69.93 | 58.71 | 48.21 | 65.16 | 68.50 | 61.58 | 60.38 |
| **5.** | ORF3a | 275 | 13.45 | 10.18 | 16.36 | 13.09 | 5.45 | 8.36 | 9.09 |
| **6.** | ORF3b | 22 | 13.64 | 0.00 | 0.00 | 0.00 | 0.00 | 0.00 | 0.00 |
| **7.** | ORF6 | 61 | 36.07 | 18.03 | 18.03 | 40.98 | 0.00 | 14.75 | 22.95 |
| **8.** | ORF7a | 121 | 12.40 | 0.00 | 8.26 | 16.53 | 0.00 | 0.83 | 1.65 |
| **9.** | ORF7b | 43 | 23.26 | 0 | 13.96 | 58.14 | 0 | 6.98 | 9.30 |
| **10.** | ORF8 | 121 | 0.00 | 0.00 | 4.13 | 8.26 | 0.00 | 0.00 | 0.00 |
| **11.** | ORF9b | 97 | 11.34 | 5.15 | 47.42 | 18.56 | 6.19 | 15.46 | 10.31 |
| **12.** | ORF10 | 38 | 10.53 | 0.00 | 0.00 | 26.32 | 2.63 | 2.63 | 0.00 |
| **13.** | ORF14 | 73 | 21.92 | 8.22 | 0.00 | 26.02 | 0.00 | 4.10 | 0.00 |

**Supplementary Table 2.** Evaluation of intrinsic disorder in structural and accessory proteins of Human SARS.

| **Sr. No** | **Protein** | **Length** | **PPID_VSL2** | **PPID_VL3** | **PPID_VLXT** | **PPID_FIT** | **PPID_IUPRED_Long** | **PPID_IUPRED_Short** | **PPID_Mean** |
| --- | --- | --- | --- | --- | --- | --- | --- | --- | --- |
| **1.** | Spike glycoprotein | 1255 | 8.92 | 5.42 | 5.18 | 2.95 | 0.08 | 0.72 | 1.12 |
| **2.** | Envelope | 76 | 26.32 | 0.00 | 13.16 | 30.26 | 0.00 | 9.21 | 6.58 |
| **3.** | Membrane protein | 221 | 7.24 | 0.00 | 8.60 | 9.50 | 0.00 | 4.52 | 1.36 |
| **4.** | Nucleoprotein | 422 | 72.75 | 64.22 | 50.24 | 66.82 | 75.36 | 67.30 | 71.09 |
| **5.** | ORF3a | 274 | 18.98 | 6.20 | 15.69 | 10.22 | 8.39 | 9.12 | 8.76 |
| **6.** | ORF3b | 154 | 28.57 | 24.02 | 14.28 | 16.23 | 0 | 6.4 | 7.14 |
| **7.** | ORF6 | 63 | 31.75 | 25.40 | 19.05 | 41.27 | 0.00 | 14.29 | 20.63 |
| **8.** | ORF7a | 122 | 11.48 | 0.00 | 9.84 | 15.57 | 0.00 | 0.82 | 0.82 |
| **9.** | ORF7b | 44 | 22.72 | 0 | 2.27 | 59.09 | 0 | 4.54 | 4.54 |
| **10.** | ORF8a | 39 | 20.51 | 46.15 | 0.00 | 35.90 | 0.00 | 10.26 | 2.56 |
| **11.** | ORF8b | 84 | 5.95 | 0.00 | 2.38 | 16.67 | 0.00 | 3.57 | 2.38 |
| **12.** | ORF9b | 98 | 32.65 | 21.43 | 50.00 | 34.69 | 22.45 | 24.49 | 26.53 |
| **13.** | ORF14 | 70 | 31.43 | 21.43 | 0.00 | 40.00 | 0.00 | 10.00 | 2.86 |

**Supplementary Table 3.** Evaluation of intrinsic disorder in structural and accessory proteins of Bat CoV.

| **Sr.**  **No** | **Protein** | **Length** | **PPID_VSL2** | **PPID_VL3** | **PPID_VLXT** | **PPID_FIT** | **PPID_IUPRED_Long** | **PPID_IUPRED_Short** | **PPID_Mean** |
| --- | --- | --- | --- | --- | --- | --- | --- | --- | --- |
| **1.** | Spike | 1242 | 10.79 | 5.56 | 8.45 | 4.99 | 0.24 | 0.97 | 1.85 |
| **2.** | Envelope | 76 | 26.32 | 0.00 | 13.16 | 30.26 | 0.00 | 9.21 | 6.58 |
| **3.** | Membrane | 221 | 10.41 | 0.00 | 7.69 | 11.76 | 0.00 | 4.98 | 1.36 |
| **4.** | Nucleocapsid | 421 | 70.31 | 64.61 | 47.98 | 65.08 | 69.60 | 62.71 | 65.80 |
| **5.** | ORF3a | 274 | 16.42 | 5.11 | 12.41 | 10.22 | 4.74 | 5.84 | 6.20 |
| **6.** | ORF3b | 39 | 38.46 | 20.51 | 38.46 | 76.92 | 0.00 | 17.95 | 23.08 |
| **7.** | ORF6 | 63 | 31.75 | 26.98 | 19.05 | 39.68 | 0.00 | 14.29 | 20.63 |
| **8.** | ORF7a | 122 | 11.48 | 0.00 | 1.64 | 14.75 | 0.00 | 0.82 | 0.82 |
| **9.** | ORF7b | 44 | 22.73 | 0.00 | 2.27 | 59.09 | 0.00 | 4.55 | 4.55 |
| **10.** | ORF8 | 121 | 0.00 | 0.00 | 2.48 | 9.09 | 0.00 | 1.65 | 0.00 |
| **11.** | ORF9b | 97 | 16.49 | 11.34 | 32.99 | 19.59 | 5.15 | 14.43 | 9.28 |
| **12.** | ORF14 | 70 | 18.57 | 0.00 | 0.00 | 30 | 0.00 | 5.71 | 0.00 |

**Supplementary Table 4.** Evaluation of intrinsic disorder in non-structural proteins SARS-CoV-2.

| **Sr. No** | **Protein** | **Length** | **PPID_VSL2** | **PPID_VL3** | **PPID_VLXT** | **PPID_FIT** | **PPID_IUPRED_Long** | **PPID_IUPRED_Short** | **PPID_Mean** |
| --- | --- | --- | --- | --- | --- | --- | --- | --- | --- |
| **1.** | Nsp1 | 180 | 33.89 | 17.78 | 39.44 | 14.44 | 13.33 | 15.00 | 12.78 |
| **2.** | Nsp2 | 638 | 12.23 | 8.31 | 15.67 | 5.17 | 0.16 | 1.57 | 5.17 |
| **3.** | Nsp3 | 1945 | 13.93 | 11.31 | 13.06 | 7.66 | 4.68 | 4.83 | 7.40 |
| **4.** | Nsp4 | 500 | 5.60 | 0.00 | 1.40 | 3.60 | 0.00 | 1.40 | 0.80 |
| **5.** | Nsp5 | 306 | 6.21 | 0.00 | 2.94 | 5.88 | 0.00 | 3.27 | 1.96 |
| **6.** | Nsp6 | 290 | 4.83 | 0.00 | 1.03 | 6.21 | 0.00 | 2.76 | 1.03 |
| **7.** | Nsp7 | 83 | 16.87 | 18.07 | 9.64 | 25.30 | 0.00 | 10.84 | 9.64 |
| **8.** | Nsp8 | 198 | 42.42 | 27.78 | 25.76 | 32.32 | 1.01 | 8.08 | 23.74 |
| **9.** | Nsp9 | 113 | 9.73 | 0.00 | 13.27 | 16.81 | 0.00 | 7.96 | 7.08 |
| **10.** | Nsp10 | 139 | 10.07 | 0.00 | 12.95 | 14.39 | 2.16 | 5.76 | 5.04 |
| **11.** | Nsp12 | 932 | 4.40 | 0.00 | 3.11 | 2.36 | 0.00 | 1.18 | 0.43 |
| **12.** | Nsp13 | 601 | 5.99 | 0.00 | 10.98 | 2.83 | 0.00 | 0.67 | 0.67 |
| **13.** | Nsp14 | 527 | 6.45 | 0.00 | 0.00 | 3.04 | 0.76 | 2.85 | 0.38 |
| **14.** | Nsp15 | 346 | 9.25 | 0.00 | 7.23 | 6.07 | 0.00 | 2.60 | 1.73 |
| **15** | Nsp16 | 298 | 10.07 | 0.00 | 3.69 | 9.06 | 9.06 | 3.69 | 5.37 |

**Supplementary Table 5.** Evaluation of intrinsic disorder in non-structural proteins of Human SARS.

| **Sr. No** | **Protein** | **Length** | **PPID_VSL2** | **PPID_VL3** | **PPID_VLXT** | **PPID_FIT** | **PPID_IUPRED_Long** | **PPID_IUPRED_Short** | **PPID_Mean** |
| --- | --- | --- | --- | --- | --- | --- | --- | --- | --- |
| **1.** | Nsp1 | 180 | 28.89 | 13.33 | 31.11 | 14.44 | 11.67 | 12.22 | 14.44 |
| **2.** | Nsp2 | 638 | 15.20 | 10.82 | 17.08 | 3.13 | 0.63 | 1.57 | 2.04 |
| **3.** | Nsp3 | 1922 | 14.62 | 10.82 | 14.57 | 8.01 | 6.76 | 7.08 | 7.91 |
| **4.** | Nsp4 | 500 | 3.80 | 0.00 | 0.80 | 2.80 | 0.00 | 1.40 | 0.60 |
| **5.** | Nsp5 | 306 | 6.86 | 0.00 | 2.94 | 6.21 | 0.00 | 3.27 | 1.96 |
| **6.** | Nsp6 | 290 | 5.52 | 0.00 | 0.69 | 5.86 | 0.00 | 2.41 | 1.03 |
| **7.** | Nsp7 | 83 | 16.87 | 18.07 | 9.64 | 25.30 | 0.00 | 10.84 | 9.64 |
| **8.** | Nsp8 | 198 | 43.43 | 25.25 | 24.24 | 30.81 | 1.01 | 8.08 | 23.74 |
| **9.** | Nsp9 | 113 | 9.73 | 0.00 | 13.27 | 17.70 | 0.00 | 7.96 | 7.96 |
| **10.** | Nsp10 | 139 | 10.07 | 0.00 | 12.95 | 14.39 | 2.16 | 6.47 | 5.04 |
| **11.** | Nsp12 | 932 | 5.36 | 0.00 | 2.36 | 2.25 | 0.00 | 1.07 | 0.43 |
| **12.** | Nsp13 | 601 | 5.99 | 0.00 | 10.98 | 2.83 | 0.00 | 0.67 | 0.67 |
| **13.** | Nsp14 | 527 | 7.59 | 0.00 | 2.66 | 2.85 | 1.52 | 2.85 | 0.38 |
| **14.** | Nsp15 | 346 | 14.74 | 0.00 | 8.38 | 7.51 | 0.00 | 2.89 | 2.60 |
| **15.** | Nsp16 | 298 | 11.07 | 0.00 | 4.03 | 8.72 | 0.00 | 3.36 | 3.02 |

**Supplementary Table 6.** Evaluation of intrinsic disorder in non-structural proteins of Bat CoV.

| **Sr. No** | **Protein** | **Length** | **PPID_VSL2** | **PPID_VL3** | **PPID_VLXT** | **PPID_FIT** | **PPID_IUPRED_Long** | **PPID_IUPRED_Short** | **PPID_Mean** |
| --- | --- | --- | --- | --- | --- | --- | --- | --- | --- |
| **1.** | Nsp1 | 179 | 26.82 | 7.82 | 19.55 | 13.41 | 10.06 | 11.73 | 12.85 |
| **2.** | Nsp2 | 639 | 14.40 | 18.00 | 16.90 | 2.50 | 0.63 | 1.41 | 2.03 |
| **3.** | Nsp3 | 1916 | 15.03 | 11.22 | 15.08 | 6.63 | 6.58 | 5.58 | 7.78 |
| **4.** | Nsp4 | 500 | 3.60 | 0.00 | 0.80 | 2.80 | 0.00 | 1.40 | 0.60 |
| **5.** | Nsp5 | 306 | 6.86 | 0.00 | 2.94 | 6.21 | 0.00 | 3.27 | 1.96 |
| **6.** | Nsp6 | 290 | 5.52 | 0.00 | 0.69 | 5.86 | 0.00 | 2.41 | 4.48 |
| **7.** | Nsp7 | 83 | 16.87 | 18.07 | 9.64 | 25.30 | 0.00 | 10.84 | 9.64 |
| **8.** | Nsp8 | 198 | 45.45 | 30.30 | 20.71 | 28.28 | 1.01 | 8.59 | 22.22 |
| **9.** | Nsp9 | 113 | 9.73 | 0.00 | 13.27 | 17.70 | 0.00 | 7.96 | 7.08 |
| **10.** | Nsp10 | 139 | 10.07 | 0.00 | 12.23 | 14.39 | 2.16 | 6.47 | 5.04 |
| **11.** | Nsp12 | 932 | 5.04 | 0.00 | 2.36 | 2.25 | 0.21 | 1.07 | 0.43 |
| **12.** | Nsp13 | 601 | 5.82 | 0.00 | 10.82 | 3.00 | 0.00 | 0.83 | 0.67 |
| **13.** | Nsp14 | 527 | 8.35 | 0.00 | 2.85 | 2.85 | 0.76 | 2.85 | 0.57 |
| **14.** | Nsp15 | 346 | 13.01 | 0.00 | 7.23 | 7.80 | 0.00 | 3.18 | 2.60 |
| **15.** | Nsp16 | 298 | 11.07 | 0.00 | 4.03 | 7.05 | 0.00 | 3.36 | 3.02 |

**Supplementary Table 7:** Predicted MoRF residues in Human SARS proteins.

| **Sr. No** | **Proteins** | **MoRFchibi_web** | **ANCHOR** | **MoRFPred** | **DISOPRED3** |
| --- | --- | --- | --- | --- | --- |
| **1.** | Spike glycoprotein | 1247-1254 | - | 1-9 | 1-10, 19-24 |
| **2.** | Envelope | 44-76 | - | - | 26-30 |
| **3.** | Membrane protein | 186-211, 215-220 | - | 214-221 | 1-6, 206-221 |
| **4.** | Nucleoprotein | 1-18, 34-44, 84-88, 140-114, 370-376, 389-398, 400-409 | 1-28, 30-77, 104-119, 130-140, 153-188, 202-236, 268-315, 346-364, 380-422 | 11-20, 48-56, 106-117, 163-169, 219-229, 269-275, 398-407 | 1-21, 26-48, 178-208, 233-250, 364-389, 403-408, 410-422 |
| **5.** | ORF3a | 1-17, 257-273 | - | 7-12, 259-263 | - |
| **6.** | ORF3b | 32-37, 41-70, 125-153 | - | - | - |
| **7.** | ORF6 | 31-63 | - | - | - |
| **8.** | ORF7a | 40-47, 72-87 | - | - | 1-10 |
| **9.** | ORF7b | - | - | - | - |
| **10.** | ORF8a | 1-39 | - | - | - |
| **11.** | ORF8b | 1-83 | - | - | - |
| **12.** | ORF9b | 3-95 | 6-14 | 44-48, 90-98 | - |
| **13.** | ORF14 | 1-34, 47-59 | - | - | - |
| **14.** | Nsp1 | 66-73, 99-179 | - | 137-144, 172-178 | 1-6 |
| **15.** | Nsp2 | - | - | - | - |
| **16.** | Nsp3 | - | 105-199 | 102-112, 189-194, 530-534, 957-965 | 515-523 |
| **17.** | Nsp4 | - | - | 495-500 | 1-26 |
| **18.** | Nsp5 | 3-8 | - | - | - |
| **19.** | Nsp6 | - | - | - | 1-22 |
| **20.** | Nsp7 | 1-30, 44-58, 64-83 | - | - | - |
| **21.** | Nsp8 | 181-185 | - | 89-93 | - |
| **22.** | Nsp9 | 40-44, 51-56, 62-77, 83-98 | - | - | - |
| **23.** | Nsp10 | 16-33, 90-99, 132-138 | - | 11-18 | - |
| **24.** | Nsp12 | - | - | 11-15 | 1-6 |
| **25.** | Nsp13 | - | - | - | - |
| **26.** | Nsp14 | - | - | 8-13, 441-445 | - |
| **27.** | Nsp15 | - | - | - | - |
| **28.** | Nsp16 | - | - | 151-156 | - |

**Supplementary Table 8:** Predicted MoRF residues in Bat CoV proteins.

| **Sr. No** | **Proteins** | **MoRFchibi_web** | **ANCHOR** | **MoRFPred** | **DISOPRED3** |
| --- | --- | --- | --- | --- | --- |
| **1.** | Spike | 38-42,1238-1245 | - | 3-12 | 1-10 |
| **2.** | Envelope | 46-75 | - | - | 26-30 |
| **3.** | Membrane | 187-221 | - | 211-221 | 1-6,205-221 |
| **4.** | Nucleocapsid | 1-20,83-88,103-116,395,409 | 1-75,104-116,152-181,201-235,267-280,286-308,345-363,379-421 | 10-21, 49-58, 106-115, 161-167, 219-228, 268-273, 397-409 | 1-19,25-47,179-208,236-249363-388,402-407,409-421 |
| **5.** | ORF3a | 1-11,260-268 | - | 8-12 | - |
| **6.** | ORF3b | 1-38 | - | - | - |
| **7.** | ORF6 | 30-60 | - | - | - |
| **8.** | ORF7a | 39-51,77-88 | - | - | 1-10 |
| **9.** | ORF7b | - | - | - | - |
| **10.** | ORF8 | 26-53,70-91,98-104,113-130 | - | - | - |
| **11.** | ORF9b | 4-21,37-96 | - | 89-97 | - |
| **12.** | ORF14 | 1-20,29-47,52-59,65-69 | - | - | - |
| **13.** | Nsp1 | 65-73,100-179 | - | 137-144, 172-180 | 1-6 |
| **14.** | Nsp2 | 2-6 | - | - | - |
| **15.** | Nsp3 | - | 126-191 | 85-89, 102-110, 183-187, 524-528, 951-959 | - |
| **16.** | Nsp4 | - | - | 495-500 | 1-27 |
| **17.** | Nsp5 | 3-8 | - | - | - |
| **18.** | Nsp6 | - | - | - | 1-21 |
| **19.** | Nsp7 | 1-30,39-58,65-83 | - | - | - |
| **20.** | Nsp8 | 181-185 | - | 89-93 | - |
| **21.** | Nsp9 | 51-56,62-77,83-102 | - | - | - |
| **22.** | Nsp10 | 23-33,90-99,132-138 | - | 11-18 | - |
| **23.** | Nsp12 | - | - | 11-15 | 1-6 |
| **24.** | Nsp13 | - | - | - | - |
| **25.** | Nsp14 | - | - | 8-13, 441-445 | - |
| **26.** | Nsp15 | 2-6 | - | 9-13 | - |
| **27.** | Nsp16 | - | - | 152-156 | - |

**Supplementary Table 9:** Predicted nucleotide-binding residues in SARS-CoV-2 proteins.

| **Sr. No** | **Proteins** | **RNA-binding residues** | | **DNA-binding residues** | |
| --- | --- | --- | --- | --- | --- |
|  |  | **PPRint** | **DisoRDPbind** | **DRNApred** | **DisoRDPbind** |
| **1.** | Spike glycoprotein | 13,28,63,68,70,108-109,122,124,129,142, 151,182,185,207,218,221,233,246,251,255,269,293,321,328,334,349-350,353,354,360,362,364,366,381,383,386,389,394,403,405-407,409-414,430,460,463,485,488,537,540,542,546,550,564-565,567-568,571,575,585,593-596,637-638,641-642,645,667,669,671,673,675-676,678-680,682-689,717-719,730,751,808-815,827-828,838-839,841,843-845,847,904,914,926-929,953,968-969,971-972,978,995,998-1000,1002,1005,1019-1024,1026,1028-1032,1034-1042,1044,1049,1051,1069,1109,1136,1158-1159,1193,1211-1213,1238-1253,1269-1273 | - | 31,35,69,73,74,108,112,147,185,204,313,316,317,319,346,355,378,380,386,396,444,447,449,451,501,528,540-542,544,545,547-549,553,602,603,605,641,682,683,719,887,905,907,912,913,928,929,968,969,974,1037,1039,1045,1047,1101,1102,1105,1107,1238,1249 | 527-529,939-942,12236-1247,1249,1273 |
| **2.** | Envelope | 61,63-64,67-69 | 1,2,49-75 | 1,2,6,7,10,15,38,42,45,48,49,53,56,57,59-64,66-69 | - |
| **3.** | Membrane protein | 4-5,7,36,38-41,58,98-99,101-102,104,107,109-114,127-128,131,138,146,154,158,162-163,172,174-177,179-181,183-194,198-213 | 18-20,28-72,75-78,80 | 1-7,9,20,21,25,36-43,44,47,50,55,58,61,65,68,71-81,88,92,94-101,103-114,116,117,121,123,125-129,131,138,141,143,146-148,150-155,157,158,160,162,166,169,172-205,207-215 | 152-156 |
| **4.** | Nucleoprotein | 2-10,12,14-16,22-46,48-49,54,61,63,65,68-71,75,77-103,115-118,120,122-130,132,134-139,141-143,145-153,171-213,216,232,234-269,272-274,276-290,300,323,340-345,351,355,360,363-388,390-391,405,411-414 | - | 26,32,93,180,189,202,203 | 239-243,246-259,364-369,371,374-383 |
| **5.** | ORF3a | 12,67,74,119,122,125,134,153,161,174-175,189,193-195,239,252-254 | 107,121-124 | 6,9,11,12,14-16,22,24,27,28,30,32,34,38,40,45,58,60,61,66-68,75,76,78,91,92,116,119,122,126,132,134-136,149-151,154,172-176,184,185,189,190,192,193,195,196,198,205,206,208,211,212,216-218,220,221,223,224,227,247,248,251-254,268,270,272 | 58,67,68 |
| **6.** | ORF3b | - | 1-22 | 1- 10,12-18,20 | - |
| **7.** | ORF6 | 20,49 | - | 3,8,10,19-23,27,28,30,31,34,38-43,45-50 | - |
| **8.** | ORF7a | 39,45,78,80-81,85,118 | 97,101 | 1,2,11,14,18-21,24-28,32,36-48,51-54,57,59-65,68-85,87,89,90,94,97,98,109,115,117-120 | 76 |
| **9.** | ORF7b | 38 | 15-28 | 1,2,5,7-10,13,15,16,19,24,27-31,35-39,40,42 | - |
| **10.** | ORF8 | 25,48-56,64,68-70,97,101,115 | 121 | 2,8,11,17,18,23,28,29,40,42-46,48,50-54,67,68,73,76-80,82,89,91,101,103 | - |
| **11.** | ORF9b | 25,32-36,39,50,55,58-59 | 22-27 | 32,34-36,40,47,50,51,53,55,58-60 | - |
| **12.** | ORF10 | 22-24,34,36,38 | 10-28 | - | - |
| **13.** | ORF14 | 3-4,13-18,20-21,67 | 46-59,61-73 | 6,7,10,12,13,15,16,18-21,35 | 19-20,22,25,26 |
| **14.** | Nsp1 | 15,31,49-51,73-74,76-77,79-84,97,99,113,  119,122-137,161,163-169,171,175,179-180 | 41,43,45-47 | 129 | 125,129,132 |
| **15.** | Nsp2 | 1,4,12-14,30,46,57,60-61,63-65,69,71-73,105-108,111-119,121-124,145,148,151,155,189-190,206,218,220-228,237-239,245-250,254-256,258-259,281,283,303,325-328,330,332-333,335,338-341,365-366,370,374,377,397,504,521,535,541-544,554,636 | 337-341,343 | 119,151,222,246,256,303,328-330,332,335,338,342,532-534,538,634,636 | 109,112,327-330,340,544 |
| **16.** | Nsp3 | 1-5,22,24,212-213,216,237,240-256,258-260,262,274-276,278-279,281-282,287-288,290-292,294,297-305,317,332-336,339,345,403,407,415,434,485-486,489-497,516,525,558,563,565,567-569,578,581-582,585-587,590,592-593,632,649-650,677,694,697,711-712,739,755,757-760,762-763,766-767,782,787-792,847-857,869,877,887,888,891,901,903,906-907,909,912,927,936,938,940,941,943,944,947-948,952,955,970-971,973-978,980-984,1014-1018,1021,1031,1034,1037,1044,1052,1084,1092,1118,1130-1131,1155,1157,1160-1165,1182,1185-1187,1189,1194,1198,1206,1240,1241,1267,1291,1294,1296-1297,1301,1303,1317,1319-1320,1323,1325,1341,1348-1349,  1366-1369,1371,1376-1380,1382-1383,1385-1391,1394,1404,1407,1410,1452-1455,1467,1474,1548,1567,1577-1578,1581,1583-1584,1586-1591,1594-1604,  1606-1607,1610-1615,1617-1631,1657,1660-1661,1665-1666,1686,1693-1694,1697,1714-1716,1728-1729,1731-1735,1737-1741,1749,1778,1781-1784,1790-1792,1841,1852-1853,1859,1861,1863-1865,1868-1870,1875-1877,1879-1889,1891-1892,1899,1908-1909,1912,1921,1923,1925-1930,1943. | 20,370,580,1599-1603,1605-1611,1616,1617,1931,1933 | 287,486,586,1366,1382,1595,1602,1614,1621,1624,1782, | 241,242,286-292,407-410,491-495,579,582-584,588,972,1159-1163,1165,1380-1383,1907,1908,1945 |
| **17.** | Nsp4 | 1,5,35,46-48,70,73,75,78,82,84-88,91,97-98,101-109,112,124,169,182,214-226,235,238,266,273-274,305-306,310-312,332-333,355,374,407,409-411,413-414,443,454,456-457,460,467,477,481,500. | 241,242 | 47,51,55,74,78,82-85,134,135,182,211,214,218,220,222,228,236,238,240,241,249,266,303,305,306,396,399,401,408,411,443,450-453,455,457,459,463,464,479 | - |
| **18.** | Nsp5 | 1,2,4-5,8-12,19-27,63,65,72,75-76,80-83,88,90,94-99,101-103,107-109,125-134,137,139,144-148,150,169,172,183-185,215,224,228,263,275-276,278-280,282,303-304,306 | - | 139,147 | 90,92,95 |
| **19.** | Nsp6 | 1-10,95,115,117,133,181,183-185,187,189,210,214,233,239-243,245,249-250,252,254-261,263,278-285,290. | 113,115,117,131-134,209,210,213,284-286 | 1,2,4,5,8,9,13,63,109,129,132,137,151,173-177,187,219,224,232,233,236,242,244,247,252,253,263,264,288 | 1,2,275-280,282-290 |
| **20.** | Nsp7 | 2,5-9,24,26-27,35,36 | - | 1,2,4,7,21,25-27 | 1-12 |
| **21.** | Nsp8 | 29-30,39-40,42-43,46,51,53-58,61,71-72,75,78-80,82-83,96-97,105,109,111-114,136-139,160,162,164,168,179,190,198 | 36,38-40,45-48,53,55,56 | 71,149 | 36-39,41,42-45,77,80,88,89 |
| **22.** | Nsp9 | 1,3,6-19,36-41,55,58-60,62,80-83,86,89,91-100,103,106-109,111,113 | - | 36,38,39,92,95,95,99 | - |
| **23.** | Nsp10 | 1,3,34,40,43-44,47,49-51,53,58,68,70,74,78,80,87-89,91-93,106,113, 124-125,127-130. | - | 36,43,48-53,56,67,68,72,113,114,118,123,125,126 | - |
| **24.** | Nsp12 | 3,5-7,9-16,18,25,33,41,45,47-48,51-55,57-58,74,77,108-110,112-113,115-118,133,138-139,176-177,185,227-228,265-266,293,297,300-301,312,324-325,328,343,345-347,349,358-366,404,408,410-414,416-418,459,474,500-501,503-509,511,545,548-549,551-553,557,559-560,572-574,583,586,595,597-598,621,623-625,627,631,647,680-681,684,695,698,701,712-714,717,721,796,798,800,830,834-836,890-891 | 197,489-503,509 | 24,74,324,500,507,508,551,553,555,556,561,565,568,569,592,593,595,676,679-682,686 | 582,584,631,639-641 |
| **25.** | Nsp13 | 1,2,15,18,47,54,68,76,81,83,94-95,97-98,114,125,128-129,131,170-171,173-178,183,186-187,189,212,214,218,223,230,253,275-277,280-290,311,327-332,343,345-347,352-353,392,409,422-423,427,431,441-444,467-470,473,484-488,497,500,502,504,514-517,518-519,523,526,530-531,553,556-565,567-570,572,594,597 | 1-9,344-357,359,360,583,594-598 | 12,15,21,76,94,100,185,186,192,212,218,286,288,289,567 | 35,36,280,330,458-465,467-475 |
| **26.** | Nsp14 | 2,5-6,9-10,13,20-22,52-53,57,60-61,65,68-69,82,84,86-89,92,95,97,98,100,101,104,111-113,132,135,137,139-145,147,155,157,159,161,163-165,176,179,196,203,205-215,218-219,226-227,230-231,252,254-256,258-260,262-265,268,271-272,286,289,292-293,297,303-304,306-311,313-315,318,335-336,338-339,383,389,391,400,402-406,408,411,414-420,424,432,434-435,465,467,469,472-474,476-482,484-485,511,520 | 182,185-187,314,321-327 | 25,69,71,112,196,197,252,310,376,404,420,424,475,476,478,485 | 313-315,317-321 |
| **27.** | Nsp15 | 1,4,11-14,34,52,61-64,67-68,70,85,92-93,98-101,103-104,13-135,138,147-149,154-155,157,161,177-178,180-181,198,239,242-245,247-250,254,256-259,288-289,316,328,332,344. | 17,142,143,146 | 289 | 99,100 |
| **28.** | Nsp16 | 29,32,39,43,46,66,69,70-74,76-77,130,132-135,137,139,145,166,201,211-212,222,225,232,249,253-254,257,276-277,279,284,286-287,290-291,293-294,297 | - | 232 | - |

**Supplementary Table 10:** Predicted nucleotide-binding residues in Human SARS proteins.

| **Sr. No** | **Proteins** | **RNA-binding residues** | | **DNA-binding residues** | |
| --- | --- | --- | --- | --- | --- |
|  |  | **PPRint** | **DisoRDPbind** | **DRNApred** | **DisoRDPbind** |
| **1.** | Spike glycoprotein | 19,26,30,32,90,92-99,106, 119,120, 122, 126, 137, 139,148,159,166,167,171,210,211,239,240,256,277,280,296,308,315,318,321,336,337,341,347,349,351,353,367-368, 370, 373, 376, 378, 381, 390, 392-394, 396-401, 417, 431-445, 447, 450-451, 477, 480, 482, 488, 516, 526, 528, 532, 536-537, 547, 550, 553-554, 557-558, 560-561, 571, 574, 576, 579-582, 623, 627, 640, 661-662, 664- 672, 674-675, 699-701, 712, 730, 733, 790, 791-797, 809-810, 820-821, 823, 825-827, 829, 886, 896, 909-911, 935, 950-951, 953-954, 960, 977, 980-981, 984, 987, 1001-1006, 1008, 1010-1014, 1016-1024, 1026-1031, 1051, 1091, 1118, 1140, 1141, 1175, 1193-1195, 1221-1222, 1224-1235, 1251-1255. | 956-958 | 35,77,104-106,109-110,146,300,306,333,365,367,383,431,434,435,436,438,484,530,533,534,535,627,869,887,889,894,911,1021,1084,1087,1089 | 1018,1020,1220-1229 |
| **2.** | Envelope | 53, 61, 67-69 | 1,2,58-76 | 6,10,15,38,45,48,49,53,57,59,60,63,64,66,68 | - |
| **3.** | Membrane protein | 4,6, 35,37-40, 57, 97-98, 100-101, 103, 106, 108, 109-113, 126-127, 130, 137, 153, 157, 161-162, 171, 173-176, 179-180, 182-193, 197-211. | 11,16-23,26-82 | 1,2,4,6,20,35-43,46,49,57,64,70,71,73,74,76,80,86,87,91,94,95,97,98,100-116,124-128,145-147,149,152-154,157,161,165,168,171-175,177-192,195,197-204,207-213 | 151-157 |
| **4.** | Nuceloprotein | 1-17,22-50,55,64,66,70-72,78-104,116-119,121,123-140,142-144,146-154,172-214,217,233-270,273-291,301,324,341 | - | 34,95,151,182,190,191,205 | 240,244,246-260,365-368,370,372,376-384 |
| **5.** | ORF3a | 12,67,119,122,125,134,153,161,175,193-194,242,251-253 | 64,70,72,120,122 | 6,9,11,12,14-16,19,22,24,27,28,30,32,34,38,40,45,57,58,61,66,67,69,70,74,75,78,91,92,116,122,126,132,134-136,149-152,154,162,164,166,170,172-174,176,181,184,185,189,190,193-196,198,200,208,211,212,216-218,220,221,223,224,227,229,234,235,238,246,247,250-253,267,269,271 | - |
| **6.** | ORF3b | 19,35,38-39,41-43,51,54,101,105,107-109,126,135,142,145-147,149-152,154 | 128-154 | 1,2,4,5,7,9-19,21,23,24,26,28,31,33,35-45,51,53-55,58-61,63,64,67-73,75,77,78,81,82,86,87,89-92,94,95,98-102,104-109,112,117-119,121-128,135-138,140-142,145-152,154 | 37,38,48-52,68-77,81-101,130-148,153,154 |
| **7.** | ORF6 | 20,23,49 | - | 10,20,21,23,27,35,38,41,42,45-50 | - |
| **8.** | ORF7a | 39,45,72,78,80-81,85,119 | 99,101-116,108-110,112 | 1,2,14,18-21,25-28,32,36-47,52,53,57,59-65,68-76,78,80,81,83-85,89,94,95,98,99,107-121 | - |
| **9.** | ORF7b | - | 11-14,16-28,36,37,39-44 | 1-3,5,8,9,10,13,15,19,21,24,28,30,31,35,42,43 | - |
| **10.** | ORF8a | 27,39 | 12-28 | 2,8,11,14,17,18,19,21,22,25,26,27,29,38,39 | 1,4,5 |
| **11.** | ORF8b | 11-16,30-33,51,55,67,80-84. | 80,81 | 4,8-18,20,26,29,31-36,38,41,43,45,47,49,59,61,64,67-69,73,81-84 | - |
| **12.** | ORF9b | 6,33,35-36,51,56,59-60. | - | - | - |
| **13.** | ORF14 | 4,13-18,20-21,67. | 41-70 | 6,10,12,13,15,16,18-21,67 | - |
| **14.** | Nsp1 | 15,31,49-51,73-74,76-77,79-84,99,103,113,122-137,161,163-165,167-169,171,175,179-180 | 2-4 | 11,96,97,99,124-126,128-130,132-134,163-167 | 132 |
| **15.** | Nsp2 | 1,4,12-14,30,57,59,60-61,63-65,68-69,72,100,106-113,115,119,124,145,148,155,206,216,218-228,237,239,242,246-248,253-256,265,325-326,330,332,338,359,362,366,368-370,397,  420,424,490,496,504,525-526,541-542,545-546,554,619-620,623-624,634,636-638 | 505,507,521-523 | 222,256,329,335 | 102,109,110,330 |
| **16.** | Nsp3 | 1-6,12,23,25,104,222,224,226,228-229,251-256,259-260,265-266,269,275-283,286,295-296,313,317,325,349,353-360,362-364,385,391,415,421-424,426,459-463,466,472-473,476,483,531,534-535,539,541,543-545,554,557-558, 561-563,566,568-569,608,625-626,653,670,673,688,730,732,734-737, 739-740, 743-744,749,753,759,767-768,823-834,836,846,854,864-865,868,878,880-881,883-884,886,889,912-913,915,917,921,925,929,932,947,948,950-952,954-955,957-961,991-995,998,1008,1011,1014,1019-1021,1035,1061,1095,1106-1108,1132,1134,1137-1140,1142,1162-1164,1166,1214,1217-1219,1224,1226,1268,1270-1271,1273-1274,1278,1280,1290,1294-1296,1298-1299,1314,1323,1326,1329-1331,1334,1343-1345,1348,1352-1353,1359-1360,1363,1367,1371,1381,1384,1386-1387,1415,1443-1444,1451,1472,1532,1544-1554-1555,1558,1560-1568,1571-1581,1583-1584,1587-1592,1594-1608,1634,1637-1638,1642-1643,1663,1670-1671,1674,1692,1705-1706,1708-1712,1714-1718,1726,1755,1759-1760,1767-1768,1797,1818,1829-1830,1836,1838,1840-1842,1845-1847,1852-1867,1869,1876,1885-1886,1889-1904, 1906 - 1907,1920. | 1-27,331-332,334-344,346,353-357,359,360 | 237,653,917,1343,1591,1598,1601 | 235,265-270,381-388,466-472,555,558-560,718-721,912-914,1136-1140,1142,1143,1149,1357-1363,1368,1369,1371-1375,1713-1715,1884,1885,1922 |
| **17.** | Nsp4 | 1,47,70,73,75,78,84-88,91, 97-98,101-105,107-109,112, 117,124,169,182,211,214- 226, 235,238,266,273-274,305-306,310-311,317-318, 355, 374,396, 399,402,406-407,409-411,413-414,426-427,443,454,456-457,460,467,477,481,500. | 297-300 | 39,55,66,67,70,74,76-80,82-86,134-135,153,169,182,211,214,218-220-222,228,236,238-241,248,249,303,305,306,309,312,313,392,396,399,401,406,407,408,411,443,444,450-453,455,457,459,463,464,479,481,491,493 | 1-11 |
| **18.** | Nsp5 | 1,2,4-5, 8-12,19-27,63,65,72,75-76,80-83,88,90,94-99,101-103,107-109,112,125-134,137,139,144-148,150,169,172,183-185,215,224,228,263,275-276,278-280,282,303-304,306 | - | 25,95,131,133-135,137,139,147,172,279 | - |
| **19.** | Nsp6 | 1-10,129,133-134,181,183-185,187,214-215,233,239-243,245,249-250,252,254-261,263,278-285,290. | 113,115-117,132,134,141,175,209,220 | 1-5,8-10,12,13,32,38,61-63,85,108,109,111,129,132,137,150,151,153,154,173-177,214,218,219,224,225,229,232,233,236,242,244,245,247,252-255,257,260,263,264,272,274,279-282,287,288,290 | 1-7,12,55-59,271-273,275,277,280,283-290 |
| **20.** | Nsp7 | 1-2,5-9,24,26-27,35-36. | - | 1,2,4,7,21,25-27 | - |
| **21.** | Nsp8 | 30,39,40,42-43,46,51,53-58,61,71-72,75,78-80,82-83,96-97,103,105,109,111-114,136-139,160,162,164,168,190,198 | 39-40,45-48,53,56 | - | 36-39,41,43-46,77,80,88,89 |
| **22.** | Nsp9 | 1,3,6-19,31,34,36-41,55,58-60,62,80-83,86,88-89,91-100,103,106-109,111,113. | - | 34,36,38,39,81,92,95,95,99 | - |
| **23.** | Nsp10 | 1,3,34,40,43-44,47,49-51,53,58,68,70,72,74,78,80,87-89,91-93,106,113,124-125,127-130. | - | 36,43,48-52,56,67,68,72,113,114,118,123,125,126 | - |
| **24.** | Nsp12 | 2,5-7,9-10,12-16,18,25,33,41,45,47-48,51-55,57-58,74,77,108-110,112-113,115-118,133,138-139,176-177,185,227-228,266,293,297,300,301,312,324,325,328,343,345,346,347,  349,358,359,360,361,362,363,364,365,366,404,408,410,411,412, 413,414,416,417,418,459,474,500,501,503,504,505,506,507,508,509,510,511,545,548,549,551,552,553,557,559,560,572,573,574,583,586,592,595,596,597,621,623,624,625,627,631,647,680,681,684,695,698,701,712,713,714,717,721,796,798,800,830,834,835,836,890,891. | 80,81,197-211,213,214,217,218,489-503,509,739 | 24,55,74,324,411,500,507,508,549,551,553,555,556,561,565,568,569,592,593,595,676,679-682,683,686 | 582-584,631 |
| **25.** | Nsp13 | ,2,15,18,47,54,68,76,81,83,94-95,97-98,114,125,128-129,131,170-171,173-178,183,186-187,189,212,214,218,223,230,253,275-277,280-290,311,327, 332,343,345-347,352-353,392,409,422-423,427,431,441-444,467-469,470,484-488,497,500,502,504,514-519,523,526,530-531,553,556-565,567-570,572,594,597 | 1-9,344-357,359,360,595,596-599 | 12,15,21,70,76,94,100,180,185,186,188,192,212,286,288,289,567 | 35,36,280,330,458-465,467-475 |
| **26.** | Nsp14 | 2,5-6,9-10,13,20-22,53,57,60-61,65,68-69,82,84,86-89,92,95,97-98,100-101,104,111-113,132,135,137,139-145,147,155,157,159,161,163-165,176,179,196,203,205-215,218-219,226-227,230-231,252,254-256,258-260,262-265,268,271-272,286,289,292-293,297,303-304,306-311,313-315,318,335-336,339,383,389,391,402-406,408,411,414-420,424,434-435,465,467,469,472-474,476-482,484-485,511,520. | 182-187,311-316,319,321-329 | 13,25,69,71,112,196,197,219,252,254,256,310,404,420,423,424,475,476,478,485 | 171,172,313-321 |
| **27.** | Nsp15 | 1,4,11-14,34,52,61-64,67-68,70,85,92-93,98-100,104,129,132-135,138,147,151,155,157,176-178,180-181,198,202,206,239,242-245,247-250,253-254,256-259,288-289,316,328,332,344. | - | 289 | 251,252,254,255,315 |
| **28.** | Nsp16 | 29,32,39,43,46,66,69-74,76-77,  130,132-135,139,145,166,  201,211,222,225,232,249,253-254,257,277,279,284,286-287,290-291,293-294 | 1,4,5,7-18,298 | 46,46,198,201,231,232,242,253 | - |

**Supplementary Table 11:** Predicted nucleotide-binding residues in BAT CoV proteins.

| **Sr. No** | **Proteins** | **RNA-binding residues** | | **DNA-binding residues** | |
| --- | --- | --- | --- | --- | --- |
|  |  | **PPRint** | **DisoRDPbind** | **DRNApred** | **DisoRDPbind** |
| **1.** | Spike glycoprotein | 14,17-19,38,92-93,96,99-105,125-128,132,142,145-155,175,243-244,246,260,281,284,300,312,322,325,340-341,345,351,353,355,357,371-372,374,377,380,382,385,394,396-398,400-405,421,436,440-441,443,450,452,475-476,503,513,515,523-524,537,540,544-545,548,561,566-569,610,625,648,651-653,656-658,661,665,686-688,699,717,720,777-784,796-797,804,807-808,810,812-814,816,873,883,895-898,922,924,937-938,940-941,947,964,967-969,971,974,988-989,990-993,995,997-1001,1003-1011,1013-1018,1038,1078,1105,1127-1128,1162,1180-1182,1208-1209,1211-1222,1238-1242. | - | 35,115,304,310,369,371,437,440,520,522,526,529,531,856,1008,1070,1074,1076 | 23,24,29-32,1005,1007,1027-1216,1242 |
| **2.** | Envelope | 53,61,67-69. | 1,2,58-76 | 6,10,15,38,45,48,49,53,57,59,60,63,64,66,68 | - |
| **3.** | Membrane protein | 4,6,35,37-40,50,57,97-98,  101,103,106,108-113,  126-127,130,137,  145,153,157,161-162,171,  173-176,179-180,  182-193,197-212,215. | 17-23,26-82 | 1-3,4,6,20,35-43,46,49,57,64,70,71,73-76,80,86,87,91,94,95,97,98,100,102-113,115,116,122,124-128,142,145-147,149,152-154,157,161,165,168,171-175,177-192,195,197-204,207-213 | 151-157 |
| **4.** | Nucleoprotein | 1-10,12,14-15,22-49,54,63,65,69-73,77-103,115-130,132,134-139,141-143,145-153,171-213,216,232-268,269,272- 300,323,340,343,345,351,355,360,364-365,367-385,387-391,405,409,411,413-414. | 107 | 93,95,149,177,179,180,183-185,188,191,195,201-203 | 239-243,246-259,364-367,369,371,375-383 |
| **5.** | ORF3a | 12,67,119,122,125,134,153,161,175,193-194,242,251-253. | 58-76,94,104,107,120-125,127,129-132 | 6,9,11,12,14-16,19,22,24,27,28,30,32,34,38,40,45,57,58,60,61,66,67,69,70,74,75,78,91,122,126,132,134-136,149-151,153,154,170,172-174,176,181,184,185,189,190,193-196,198,200,208,211,212,216-220,224,227,229,234,235,238,246,247,250-253,267,269-271 | 244,245 |
| **6.** | ORF3b | - | 12-28 | - | - |
| **7.** | ORF6 | 20,23,49. | - | 10,20,21,23,27,28,35,38,41,42,45-50 | - |
| **8.** | ORF7a | 39,45,72,78,80-81,85,118-119. | 102,104,112,113 | 1,2,11,14,18-21,24-28,32,36-48,51-54,57,59-65,68-76,78-85,87,89,94,95,98,99,107,110,115,116,118-121 | - |
| **9.** | ORF7b | - | 12,16-28,35-44 | 1-3,5,8-10,13,15,19,21,24,28-31,35,36,38,42-44 | - |
| **10.** | ORF8 | 13,25,48,50-53,55-56,59,64-65,67-69,70,72,101. | 121 | 2,8,11,14,17,18,40-46,48-54,56,66-68,72,73,76-80,82,101,103,104,112 | - |
| **11.** | ORF9b | 32,34-35,55,58-59. | 22,23,25 | - | 49 |
| **12.** | ORF14 | 3-4,13-18,20-21,67 | 41-70 | 6,7,10,12,13,15,16,18-21 | - |
| **13.** | Nsp1 | 15,31,49-51, 73-74,76-77,79-84,99,103,113,122-137,161,163-165,167- 169,171,175. | 2-4 | 99,124-126,128-130,132-134,163-165 | - |
| **14.** | Nsp2 | 1-2,5,13-15,31,58,60-62,64-66,69-70,73,101,107-114,116,120,125,146,149,156,164,207,217,219-238,240,243,246-247,250,257,260-262,326-327,331,333,339,360-361,363,367,369-371,380,385,403,446,491,497,505,526-527,542-543,546-547,555,620-621,624-625,635,637-639. | 503,504,506-510,517-529,538,540 | 257,336 | 103,110,111 |
| **15.** | Nsp3 | 1-6,23,25,214-226,228,232,235,246,248-249,251,254,255,260-261,263-267,270-281,290-291,298,305-306,308-309,312,379-381,385-387,455-456,460,463-467,486,488,495,500,513,528,533,535,537-539,548,551-552,555-557,560,563,602,619-620,647,664,720,722,726,728-731,733-734,737-738,743,747,753,761-762,815,817-828,830,840,848,858-859,862,872,874,877-878,880,882-883,906-907,909,911,915,919,923,926,941-942,944-946,948-949,951-955,985-989,992,1002,1005,1008,1023,1026,1029,1055,1062,1101-1102,1131-1134,1153,  1156,1160,1169,1177,1211-1213,1216,1218,1260-1261,1265,1267,1271,1273-1274,1277-1280,1284-1285,1290-1291,1293,1308-1312,1316-1317,1320,1337,1339,1342,1347-1348,1353-1355,1357-1358,1361-1362,1365,1377-1378,1381,1399,1409,1425-1426,1434,1438,1466,1526,1538,1548-1549,1552,1554-1555,1557-1562,1565-1578,1581-1586,1588,1602,1628,1631-1632,1635-1637,1655,1657,1664-1665,1668,1684-1687,1699,1704-1706, 1708-1712,1720,1749,1753-1754,1761,1763,1791,1823, 1824,1830,1832,1834-1836,1839-1841,1846-1848,1850-1860,1862-1863,1870,1879,1880,1883-1892,1894-1901,1914. | 3-27,344,346-355,550,834 | 232,911,1337,1353,1585,1592,1595 | 377,378,460,549,552-554,624,625,713,906-908,1131-1134,1136,1347,1351-1357,1362,1363,1365-1369,1558,1771,1878,1879,1916 |
| **16.** | Nsp4 | 1,4-5,47,70,73,75,78,84-88,91,97-98,101-105,107-109,112,117,124,169,182,211,214-226,235,238,266,273-274,305-306,310-311,317-318,355,374,396,406-407,409-411,413-414,426-427,443,454,456-457,460,467,477,481,500 | - | 36,39,55,66,67,70,73-80,82-86,134,135,153,169,182,183,211,214,218-220,222,228,229,236,238,240,241,249,299,303,305,306,309,312,313,354,355,392,396,399,401,406-408,411,443,444,450-453,455,457,459,463,464,479-481,491,493 | - |
| **17.** | Nsp5 | 1-2,4-5,8-12,19-27,62-63,65,72,75-76,80-83,88,90,94- 99,101-103,107-109,125-134,137,139,144-146,148,150,169,172,183-185,215,224,228,263,275-276,278-280,282,303-304,306. | - | 131,134,137,139,147,172,279 | - |
| **18.** | Nsp6 | 1-10,129,133-134,181,183-185,187,214-215,233,239-243,245,249,250,252,254-261,263,278-285,290. | 113,117,209-220 | 1-5,8,9,12,13,32,61-63,108,109,111,129,132,137,151,153,154,173-177,214,218,219,224,225,229,232,233,236,242,244,245,247,252-255,257,260,263,264,272,274,279,281,282,287,288,290 | 1-7,54-59,271-273,275,277,280,283-290 |
| **19.** | Nsp7 | 1-2,5-9,24,26-27,35-36. | - | 1,2,4,7,21,25-27 | 1-12 |
| **20.** | Nsp8 | 28-30,39-40,42-43,46,51,53-58,61,71-72,75,78-80,82-83,96-97,103,105,109,111-114,136,139,160-162,164,168,190,198 | - | 0 | 35-39,41,43-49,77,80,88,89 |
| **21.** | Nsp9 | 1,3,6-19,31,34,36-41,55,58-60,62,80-83,  86,89,91-100,103,106-109,111,113. | - | 36,92,95,96,99 | - |
| **22.** | Nsp10 | 1,3,34,40,43-44,47,49-51,53,58,68,70,72,74,78,80,87-89,91-93,106,113,124-125,127-130 | - | 48,51,67,72,113,114,118,123,125,126 | - |
| **23.** | Nsp12 | 2,5-7,9-10,12-16,18,25,33,41,45,47-48,  51-55,57-58,74,77,108-110,112-113,115-118,133,138-139,176-177,184-185,188,227-228,266,293,297,300-301,312,324-325,328,343-347,349,358-366,404, 408,410-416,417, 418, 459, 474, 500-501, 503-509, 511,545,548-549,551-553,557,559-560,572-574,583-586, 595, 597, 621, 623-627, 631,647,680-681,684,695,698,701,712-714,717,721,796,798,800,830,834-836,890-891. | 80,192-214,217,218,489-503,509,739 | 24,50,55,74,324,411,500,507,508,549,551,553,555,556,561,565,568,569,592,593,595,676,679-683,686 | 257,582-584,631,641 |
| **24.** | Nsp13 | 1,4,11-14,34,52,61-64,67-68,70,85,92-93,98-100,104,129,132-135,138-139,147,151,155,157,176-178,180-181,198,202,206,239,248-250,253-254,256-259,288-289,316,328,332,344. | 1-9,181-187,344-357,359,360,582,594-601 | 12,15,21,70,76,94,100,185,186,188,192,212,286,288,289,567 | 35,36,280,329,459,461,462,463,466-472 |
| **25.** | Nsp14 | 2,5-6,9-10,13,20-22,52-53,57,60-61,65,68-69,82,84,86-89,92,95,97-98,100-101,104,111-113,132,135,137,139,140-145,147,155,157,159,161,163-165,176,179,196,203,205-215,218-219,226-227,230-231,252,254-256,258-260,262-265,268,271-272,286,289,292-293,297,303,306-311,313, 315,318,334-336,338-339,383,389,391,402-406,408,411,414-420,424,434-435,465,467,469,472-474,476-485,511,520. | 182,183,185-187,313,314,321,323-325,327 | 25,69,71,112,196,197,219,252,256,310,404,420,423,424,475,476,478,485 | 172,173,310,312-321 |
| **26.** | Nsp15 | 1,2,15,18,47,54,68,76,81,83,94-95,97-98,114,125,128-129,131,171,173-178,183,186-187,189,212,  214,218,223,230,253,275-277,280-290,311,327-332,343,345-347,352-353,392,409,422-423,  427,431,441-444,467-470,484-488,497,500,502,504,514-519,523,526,530-531,553,556-565,567-570,572,594,597. | - | 289 | 251,252,254,255,315 |
| **27.** | Nsp16 | 29,32,39,43,46,66,69,70-74,76-77,131-135,161,165-166,201,211,222,225,232,249,253-254,  257,277,279,284,286-287,290-291,293-294. | 1,4,5,7-18,298 | 74,198,232 | 136 |

**Supplementary Figures S1.** Multiple sequence alignment of structural proteins of all three studied coronaviruses are generated using Clustal Omega. The aligned images are created using Esprit 3.0.

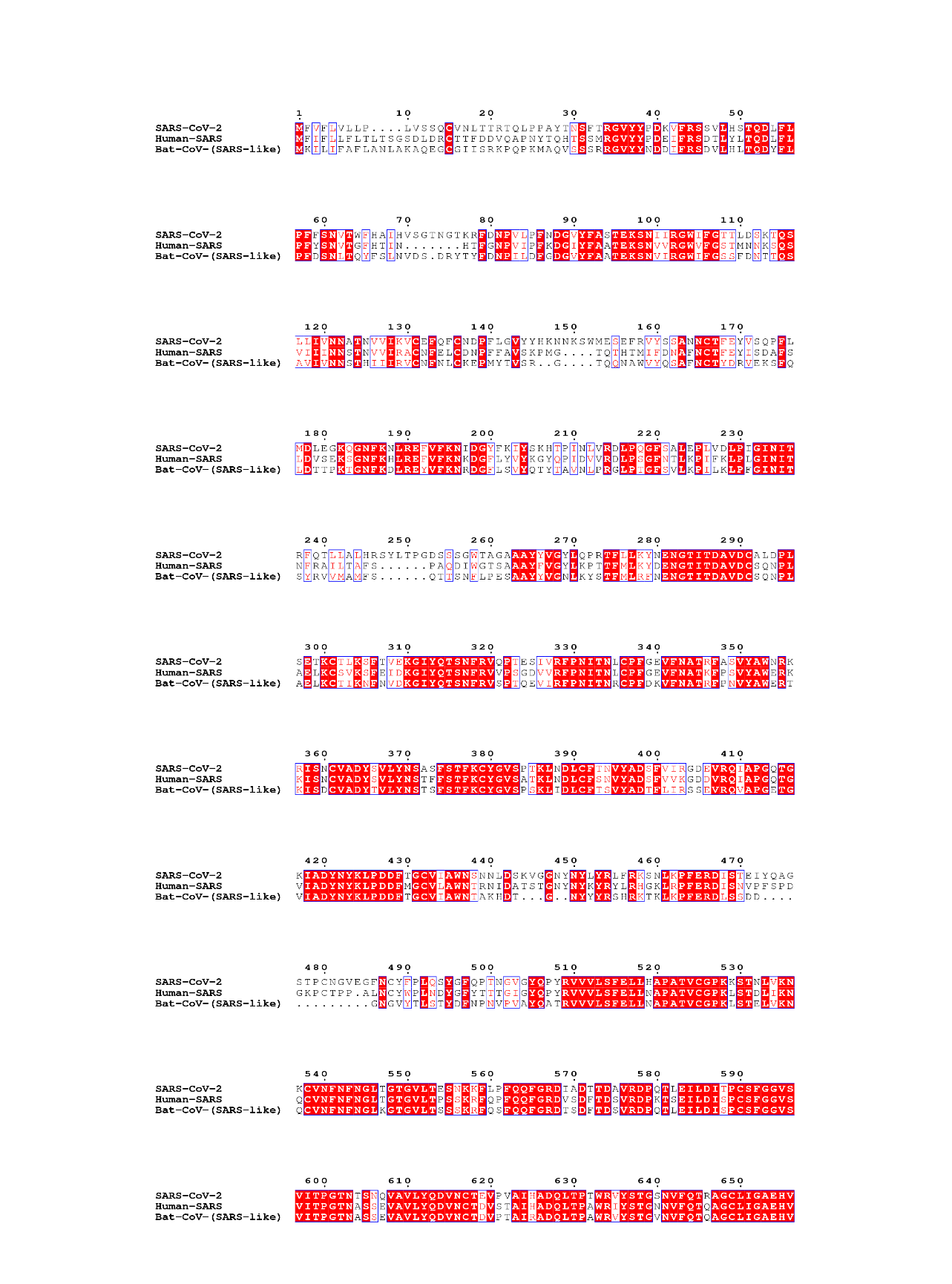

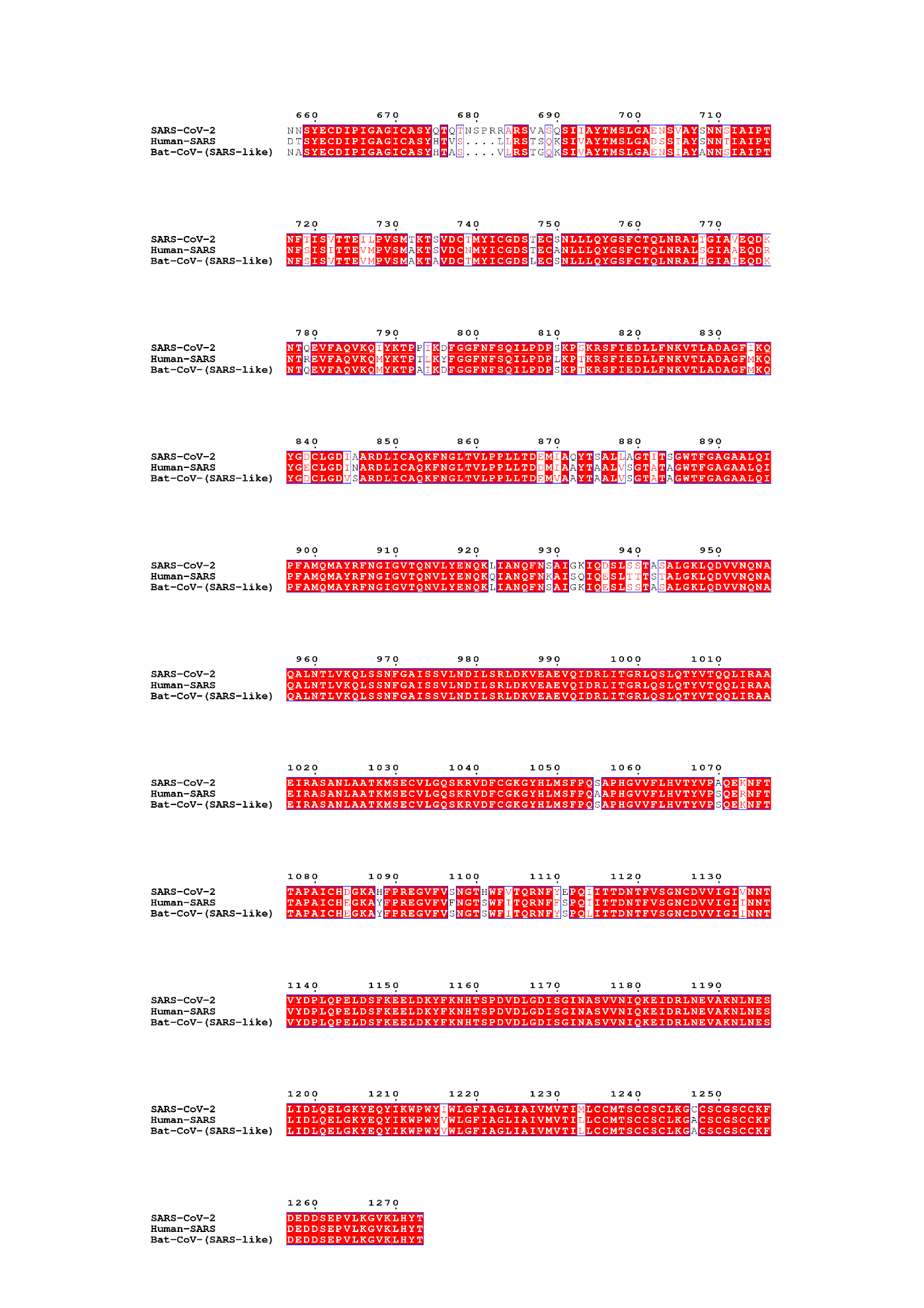

**Figure S1A.** Multiple sequence alignment of SARS-CoV-2, Human SARS, and Bat CoV spike glycoproteins.

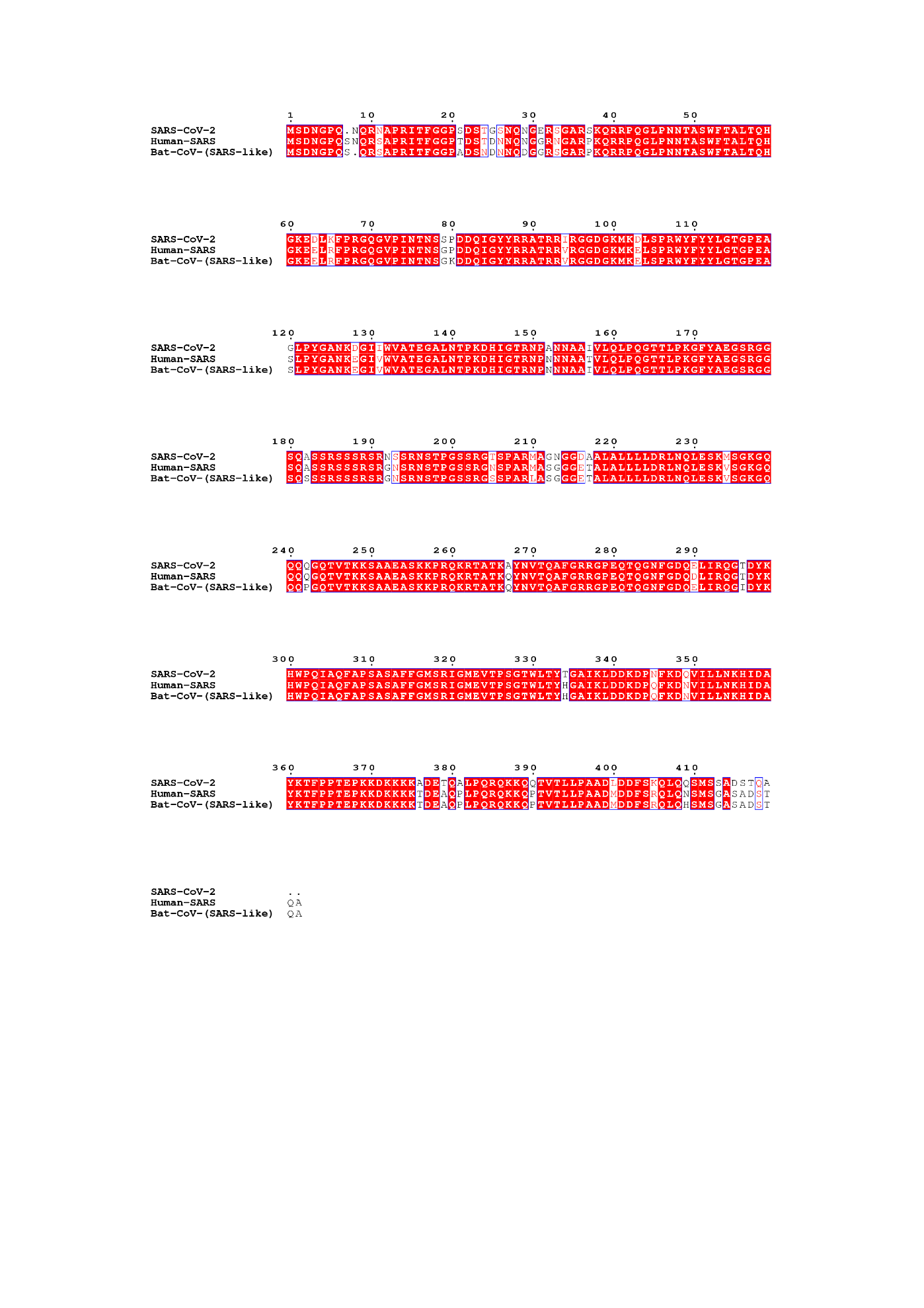

**Figure S1B.** Multiple sequence alignment of SARS-CoV-2, Human SARS, and Bat CoV Nucleoproteins.

**Supplementary Figure S2.** Multiple sequence alignment of non-structural proteins of all three studied coronaviruses are generated using Clustal Omega. The aligned images are created using Esprit 3.0.

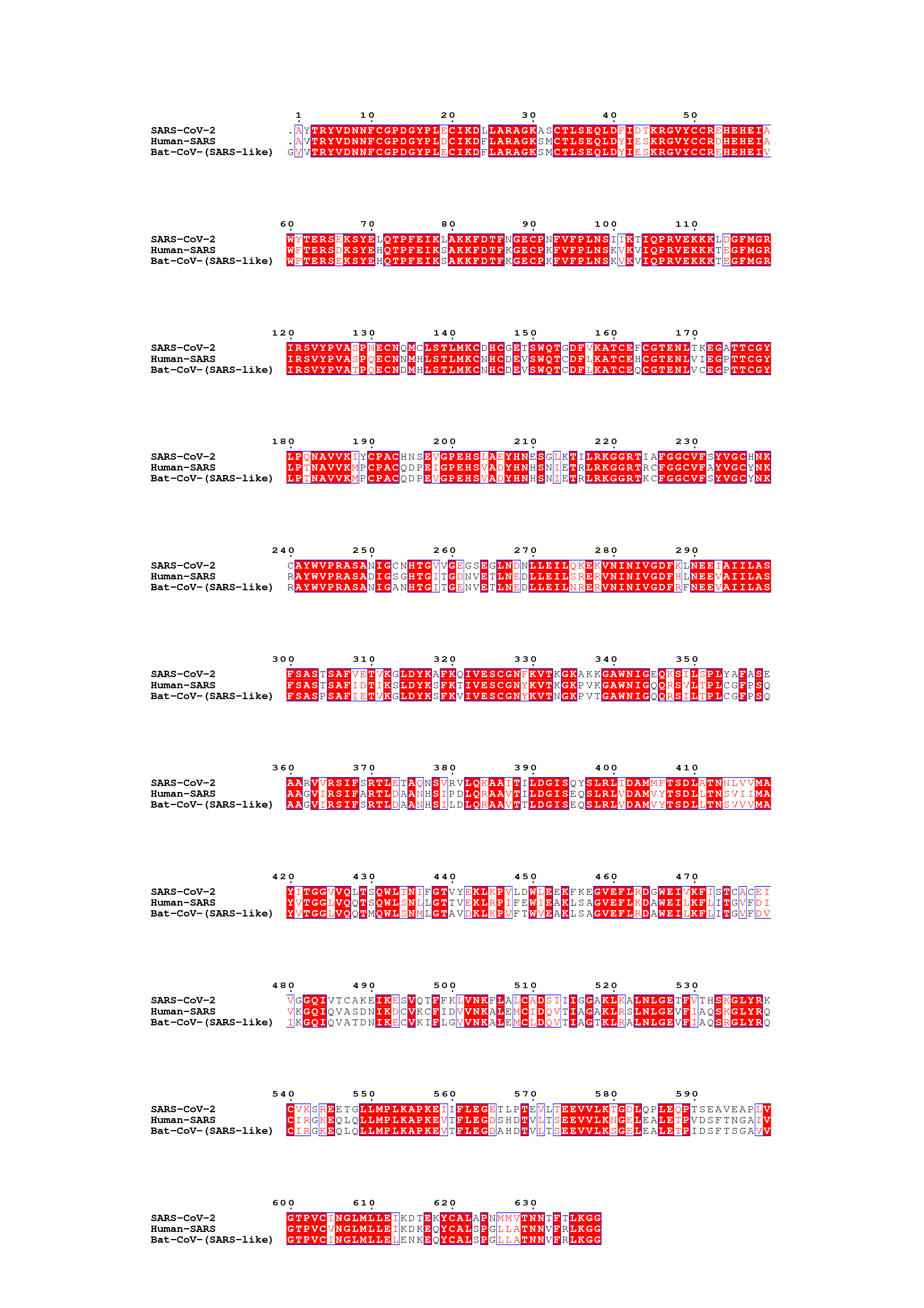

**Figure S2A.** Multiple sequence alignment of SARS-CoV-2, Human SARS, and Bat CoV Nsp2 proteins.

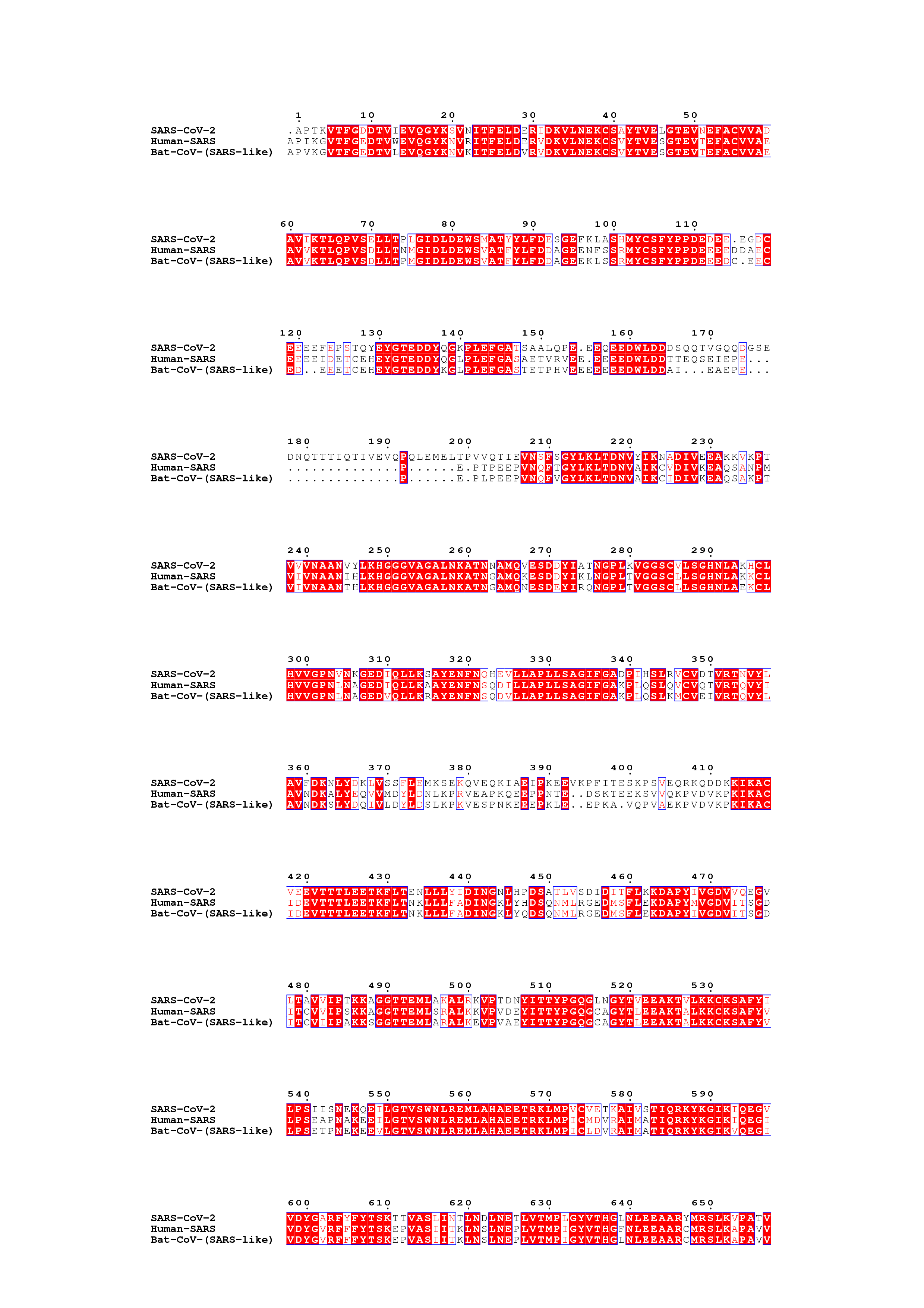

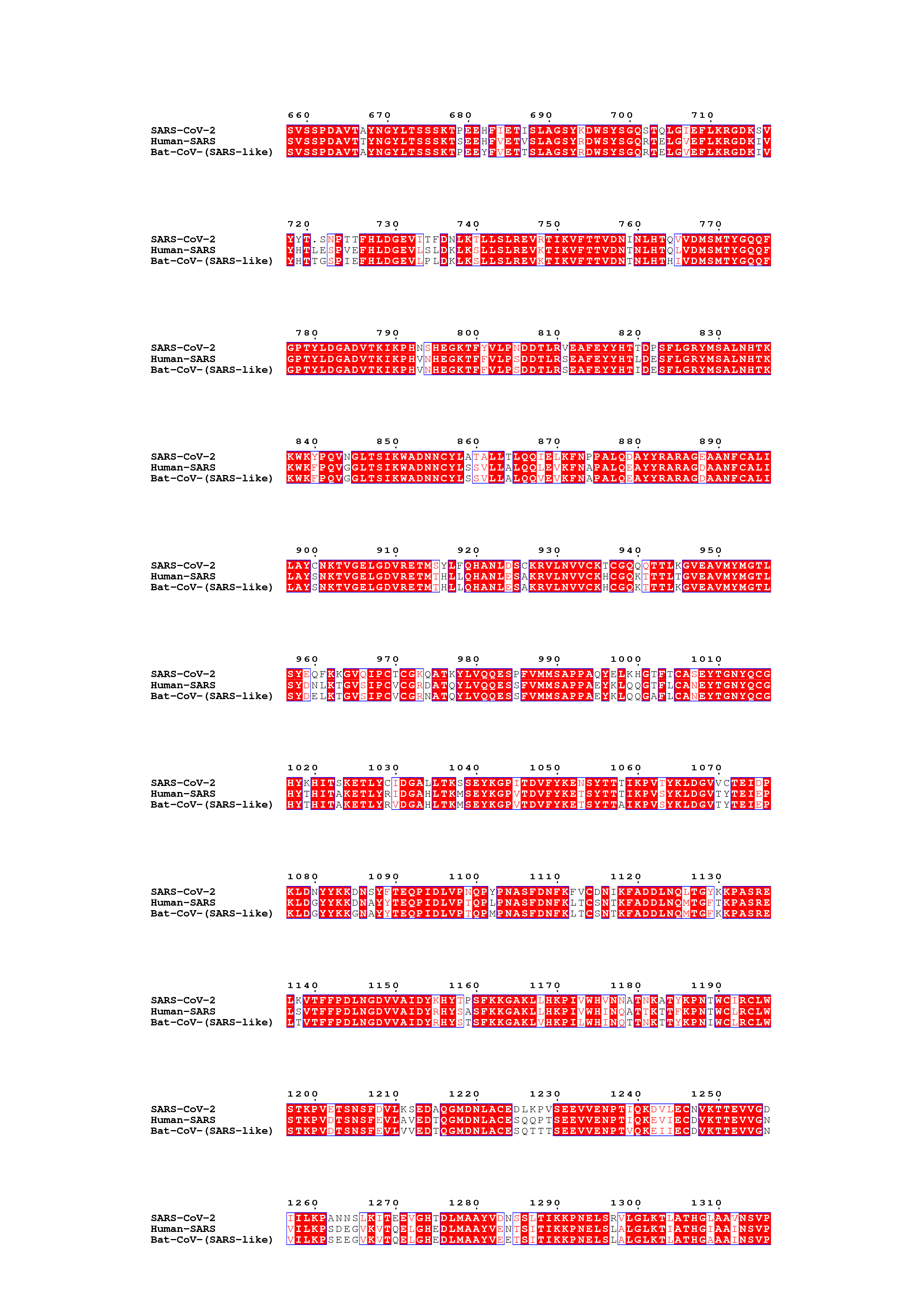

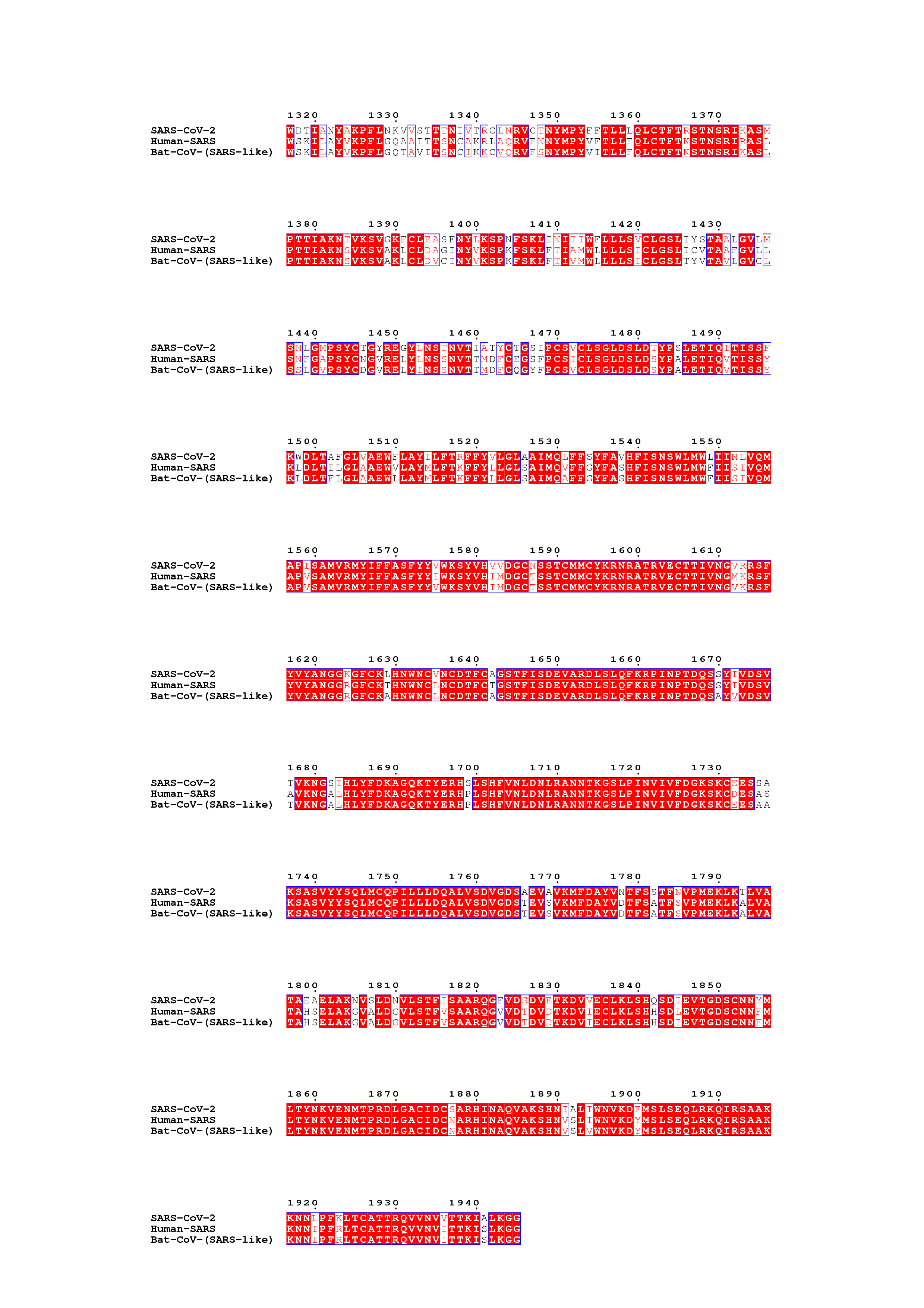

**Figure S2B.** Multiple sequence alignment of SARS-CoV-2, Human SARS, and Bat CoV Nsp3 proteins.

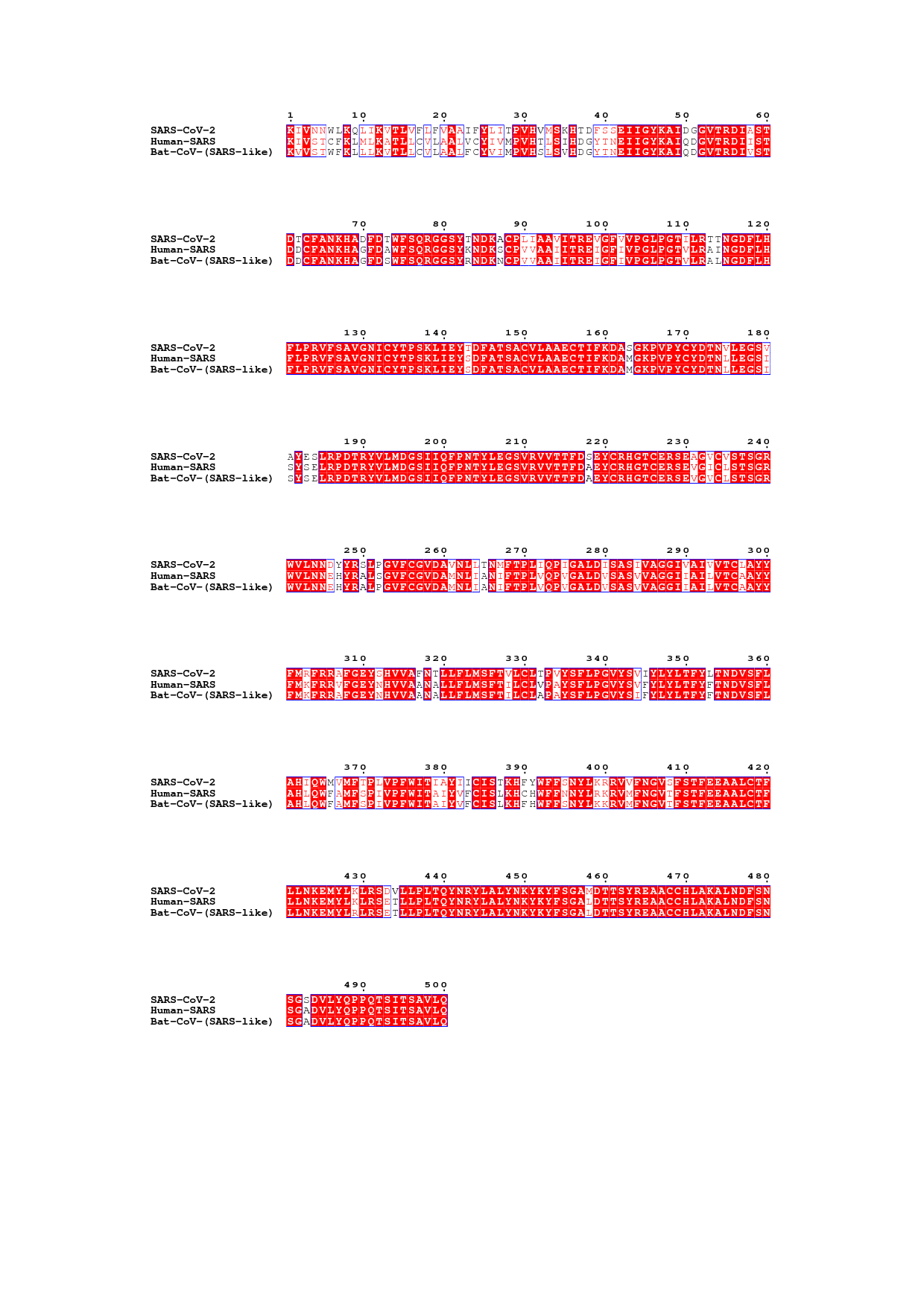

**Figure S2C.** Multiple sequence alignment of SARS-CoV-2, Human SARS, and Bat CoV Nsp4 proteins.

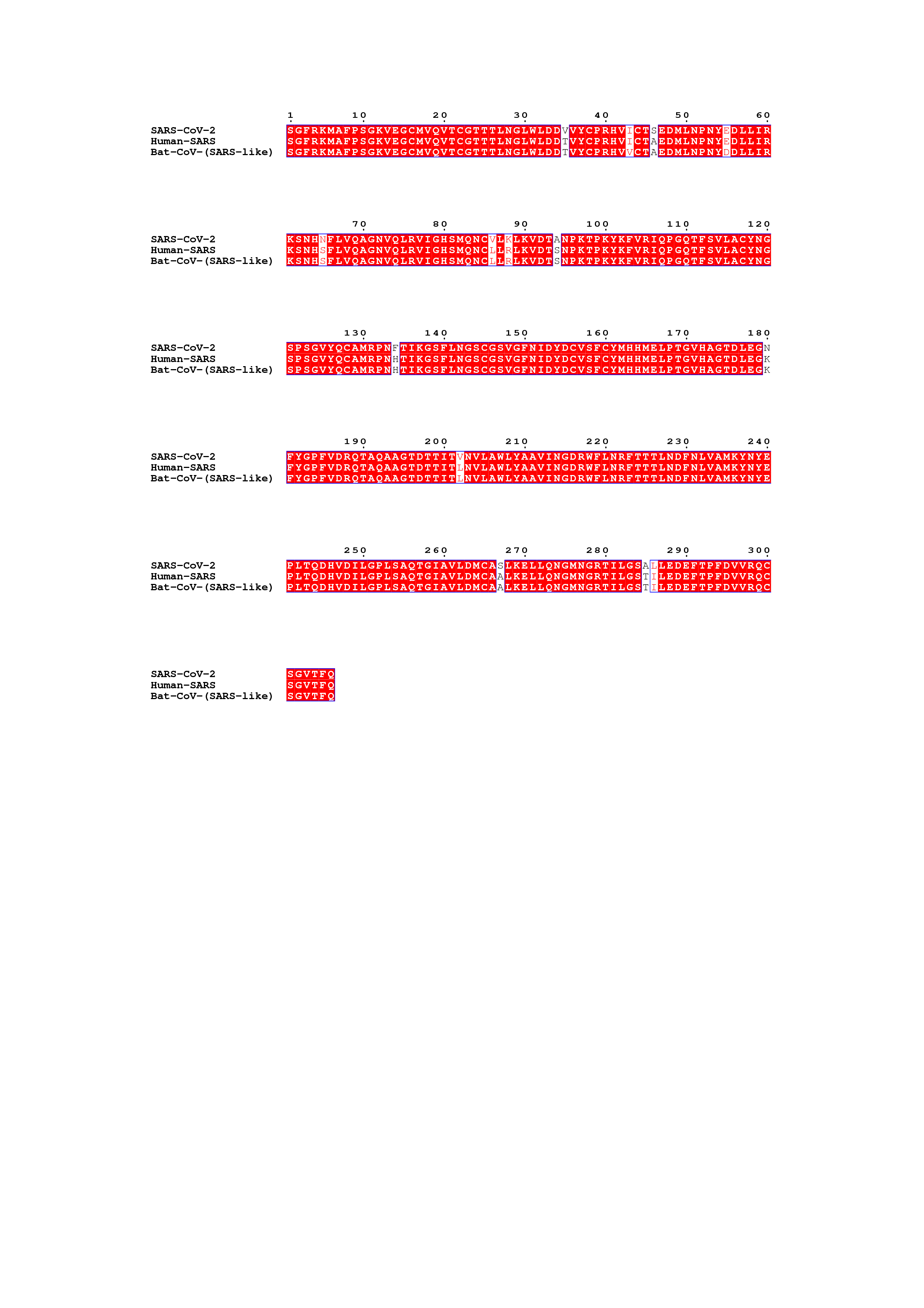

**Figure S2D.** Multiple sequence alignment of SARS-CoV-2, Human SARS, and Bat CoV Nsp5 proteins.

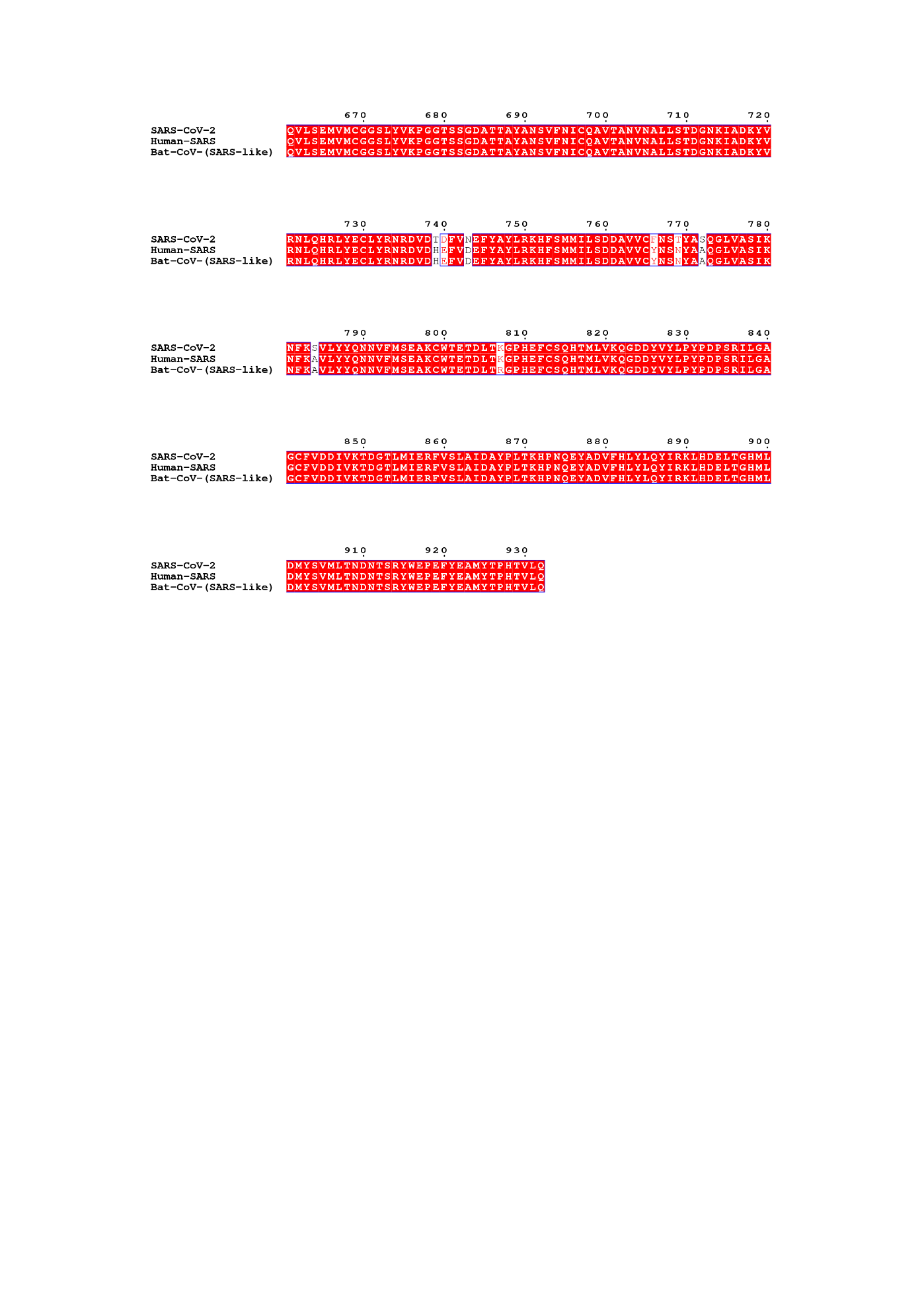

**Figure S2E.** Multiple sequence alignment of SARS-CoV-2, Human SARS, and Bat CoV Nsp12 proteins.

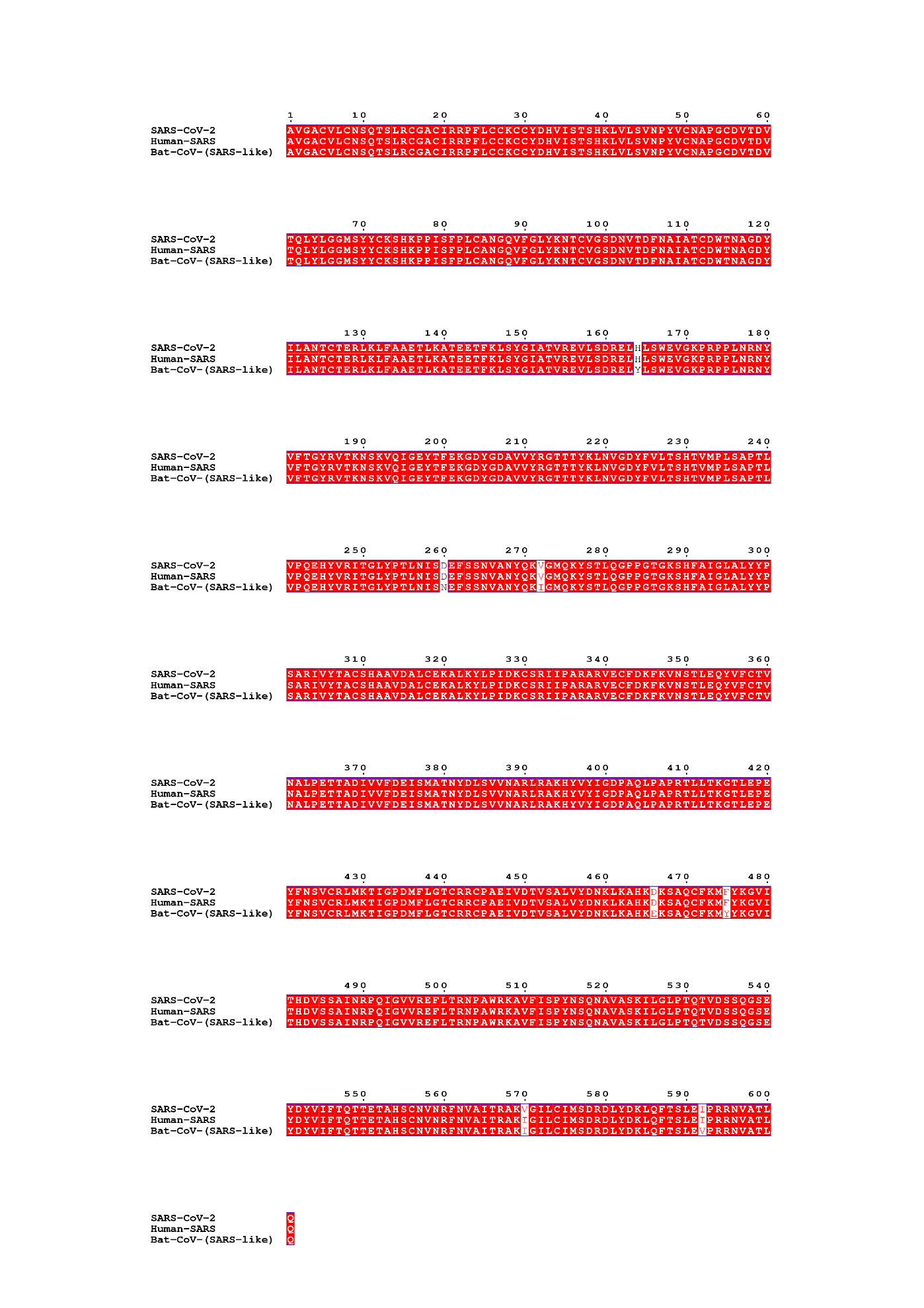

**Figure S2F.** Multiple sequence alignment of SARS-CoV-2, Human SARS, and Bat CoV Nsp13 proteins.

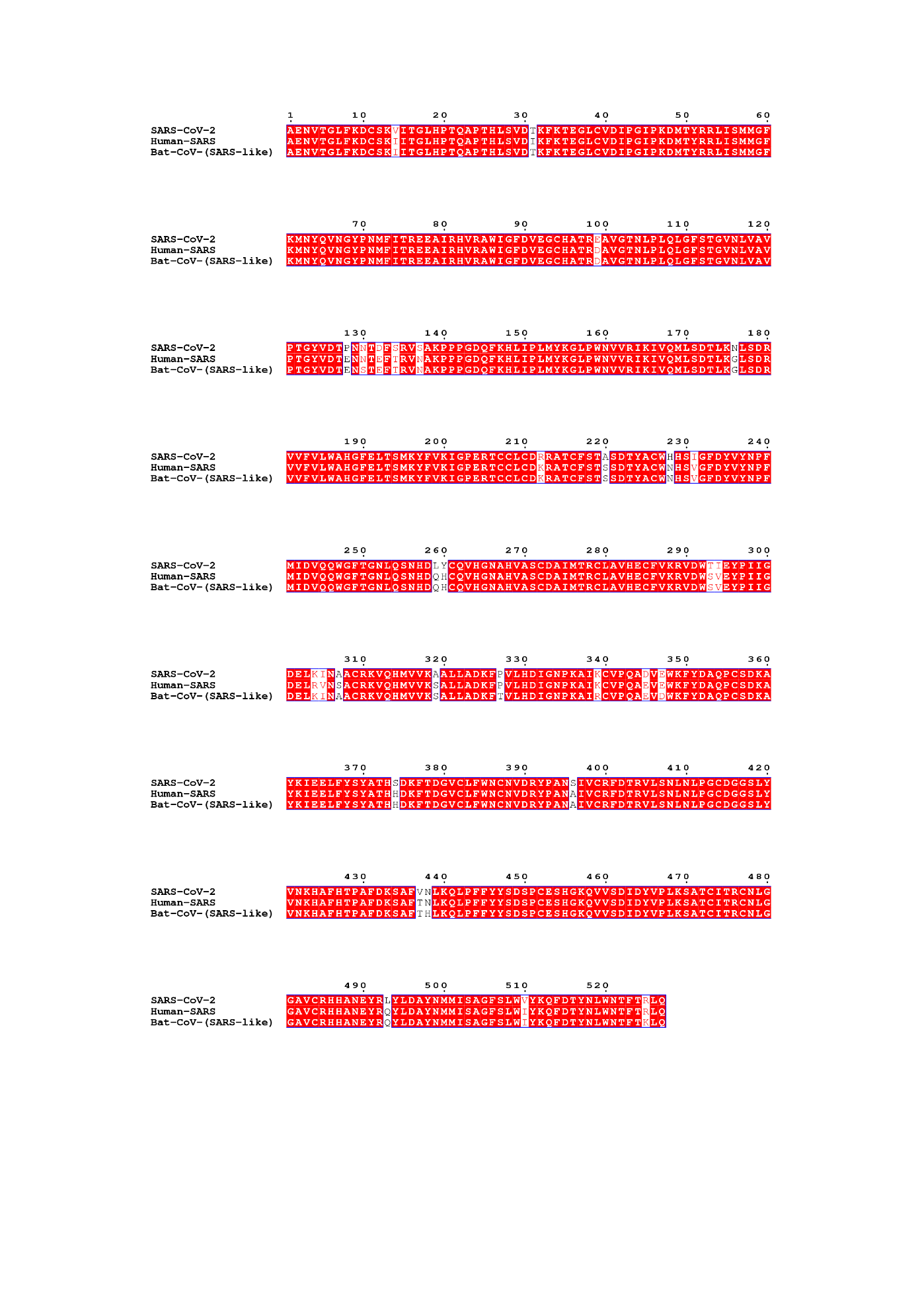

**Figure S2G.** Multiple sequence alignment of SARS-CoV-2, Human SARS, and Bat CoV Nsp14 proteins.

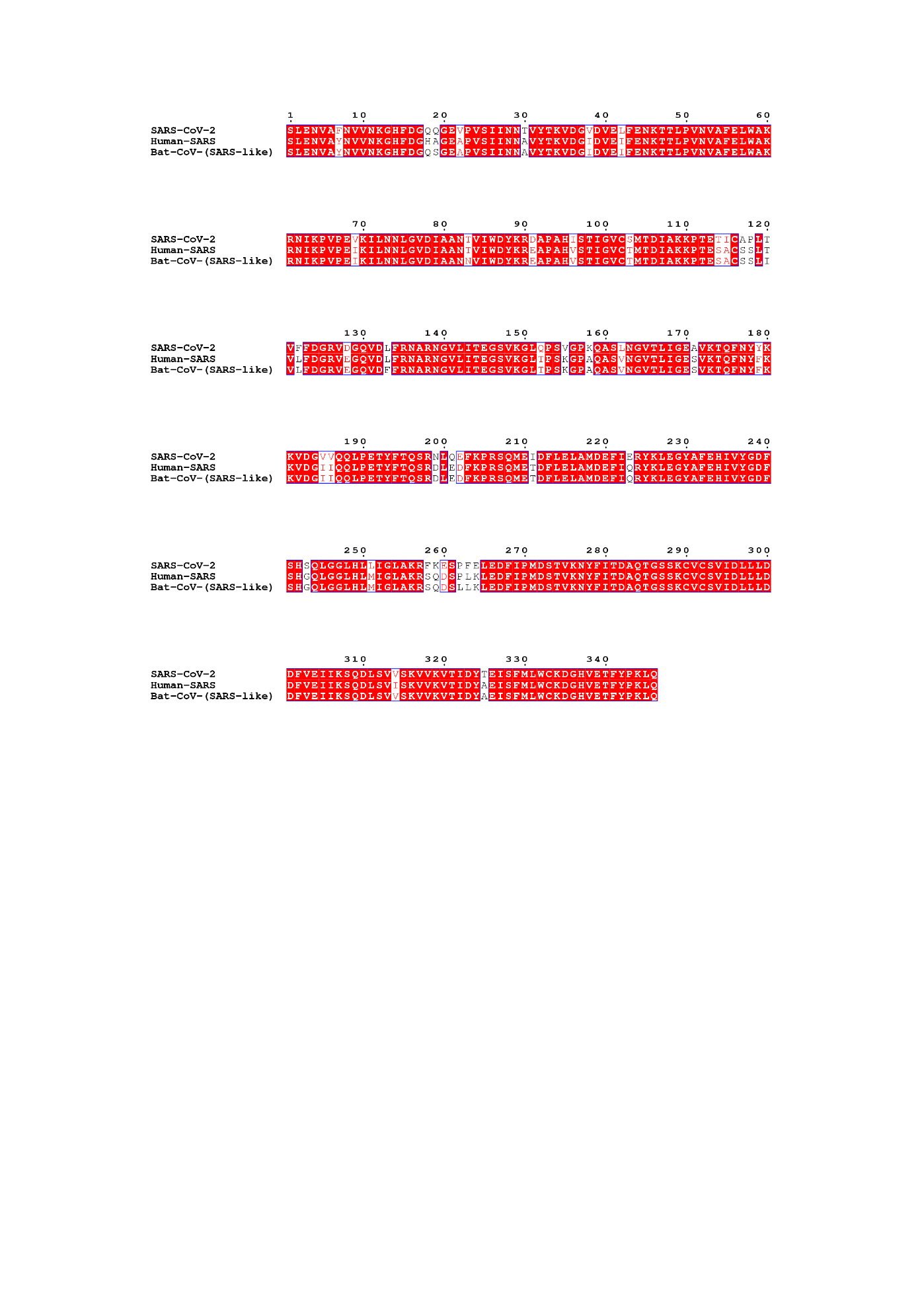

**Figure S2H.** Multiple sequence alignment of SARS-CoV-2, Human SARS, and Bat CoV Nsp15 proteins.

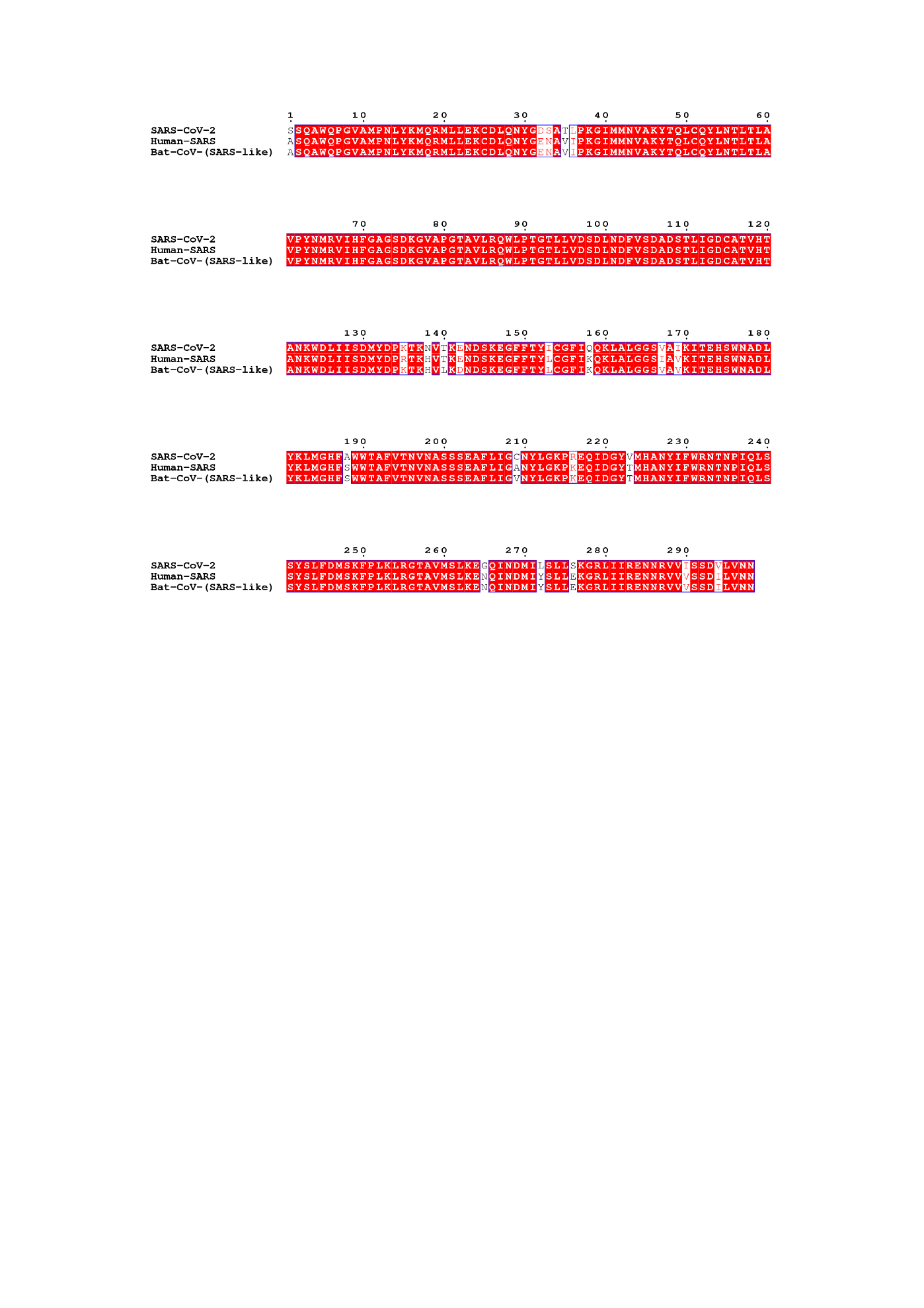

**Figure S2I.** Multiple sequence alignment of SARS-CoV-2, Human SARS, and Bat CoV Nsp16 proteins.
